# Designing a Minimal Artificial Glycolytic Pathway

**DOI:** 10.1101/2022.09.22.508994

**Authors:** Yiqun Yang, Yuwan Liu, Jie Zhang, Qiaoyu Yang, Jian Cheng, Huanyu Chu, Haodong Zhao, Mengting Luo, Xiaoyun Lu, Dingyu Liu, Xiang Sheng, Yi-Heng P. J. Zhang, Huifeng Jiang, Yanhe Ma

## Abstract

The canonical glycolysis generates two molecules of acetyl-coenzyme A (acetyl-CoA) from one glucose through eleven cascade biochemical reactions. Here, we designed and constructed a Minimal Artificial Glycolytic (MAG) pathway consisting of only three types of biochemical reactions, with phosphoketolase as the core, combined with phosphatase and isomerase as auxiliary enzymes. It could theoretically achieve a 100% carbon yield to acetyl-CoA from any monosaccharide by integrating one-carbon condensation reaction. We tested the MAG pathway *in vitro* and *in vivo*, demonstrating the catabolism of typical C1-C6 carbohydrates to acetyl-CoA with yields from 82% to 95%. This novel glycolytic pathway provides a promising route for biomanufacturing with stoichiometric productivity from multiple carbon sources in the future.

## Main Text

Glycolysis is a catabolic pathway with ten cascade biochemical reactions to convert glucose to pyruvate, which is further decarboxylated to produce acetyl-CoA, providing energy and building blocks for life survival^1,2^. Billions of years of natural selection has evolved complicated networks for glycolysis to maintain metabolic and physiological homeostasis^3-5^. Intensive efforts in the past decades have been made to reform metabolic homeostasis toward compound of interest with the maximum stoichiometry^6-10^. However, the synthesis of cellular metabolites via glycolysis has been evolved with the optimality principle in nature^11,12^, partially engineering of the central carbon metabolism is not enough to untie the complicated networks of catabolism within cells^13,14^. Here depending on chemical principles, we *de novo* designed and implemented a Minimal Artificial Glycolytic (MAG) pathway, which could completely convert any ketose into acetyl-CoA only based on three core reactions.

The thiamin diphosphate (ThDP)-dependent phosphoketolase (PK) is one of three key enzymes in nature for C-C bond cleavage^15,16^. The C2-C3 bond of diverse ketoses can be broken by PKs to produce one molecular of acetyl-phosphate (AcP), which can be further converted into acetyl-CoA by phosphate acetyltransferase (PTA)^17^. According to the catalytic mechanisms of PK, it is possible that any ketose could produce a covalent intermediate, which will produce an AcP and release a free aldose (Fig. 1A and Fig. S1)^15^. At the same time, interconversion between aldoses and ketoses can be conducted by isomerases^18,19^. Therefore, we hypothesized that all sugars could be completely converted into acetyl-CoA through multiple rounds of the carbon cleavage by PK and isomerization (Fig. 1A). Inspired by these natural and hypothetical catalytic processes, we proposed to construct a MAG pathway to achieve the production of stoichiometric amounts of acetyl-CoA from any sugar.

**Fig. 1.**
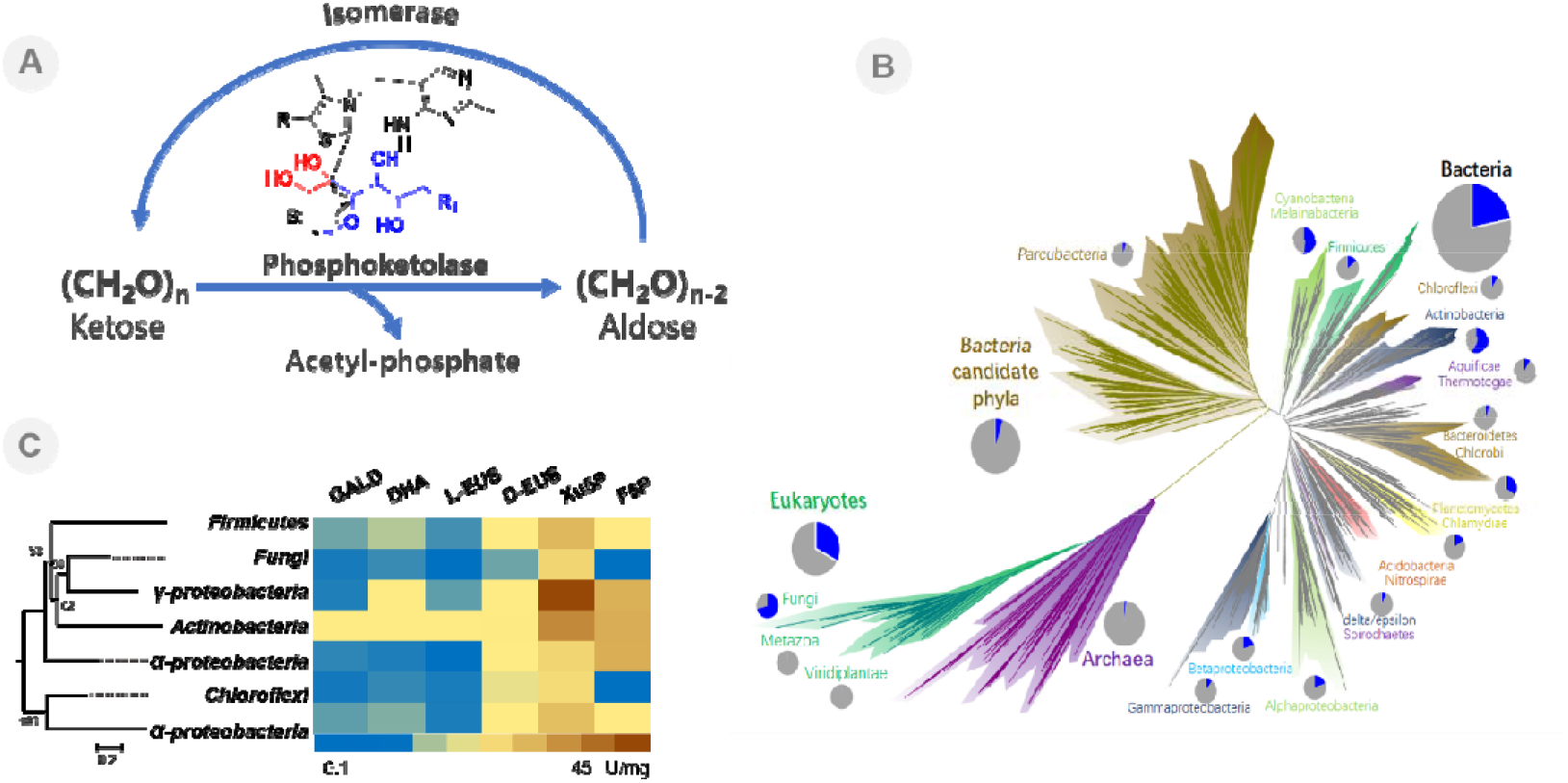
Design of the Minimal Artificial Glycolytic (MAG) pathway. (**A**) Schematic illustration of the MAG pathway. (**B**) The distribution of PKs in the tree of life. The tree was referenced the Jillian F. Banfield’s study. The blue color in pie chart represents the species with PKs in all genome sequenced species. (**C**) The catalytic activity of PKs from different species on different substrates. Seven PKs were selected from different phylum based on the phylogenetic analysis. The right table represents the catalytic activity of all candidate PKs on six classes of ketose or ketose phosphate. Different colors represent specific enzyme activity (U/mg).

In fact, PK gene originates very early in nature and widely distributes in three kingdoms of life, indicating extensively application for carbon utilization during the evolutionary processes (Fig. 1B and Fig. S2). In order to confirm the activity of PK on different sugars, we expressed 7 candidate genes from different species based on the phylogenetic tree (Fig. S3). The function of carbon cleavage on different sugars by PKs are highly conserved (Fig. 1C). Almost all PKs could catalyze short chain ketoses without phosphorylation to AcP but displayed no activity on phosphorylated short chain ketoses and aldoses (Fig. 1C and Fig. S4-S6). We used quantum-chemical analysis to determine the catalytic processes of PK on short ketoses (Fig. S7-S12 and Table S1-S3). As seen from the energy profiles, it is feasible to form 2-α, β-dihydroxyethylidene-ThDP (DHEThDP) from short chain ketoses catalyzed by PK (Fig. S9). However, the strong anchoring effect of the phosphate moiety of dihydroxyacetone phosphate (DHAP) and D-erythrulose-4-phosphate (Eu4P) made the carbonyl groups far away from the active C2 position of ThDP, which made them less prone to interact with ThDP to produce the covalent intermediate (Fig. S13). Among all of the studied PKs, PK from Actinobacteria (*Bifidobacterium*) (BbPK) displayed better activities on all of the tested ketoses.

In order to further improve the activity of BbPK on short sugars, we intended to conduct directed evolution experiment. Due to the difference between the forming process of DHEThDP from glycolaldehyde and dihydroxyacetone (DHA)^20^, we decided to independently screen mutants with better activities. Since the catalytic center of PK is formed at the interface of homodimer, we selected the residues located at the active cavity and protein-dimer interface to perform saturation mutagenesis (Fig. 2A). After screening around 3060 mutants (Fig. S14 and S15), we obtained three beneficial mutants. PK-E520I and PK-Q321A displayed 2.3-fold and 2.8-fold catalytic efficiency for DHA than wide-type BbPK, respectively (Fig. 2B and Table S4). PK-H142N showed 4.9-fold catalytic efficiency for glycolaldehyde and a 3.8-fold higher catalytic efficiency for D-erythrulose (D-EUS) than the wild-type of BbPK (Fig. 2C and Table S4). Molecular dynamics analysis indicated that the decrease of pocket volume in mutants could be the major reason for the improvement of activities (Fig. 2D).

**Fig. 2.**
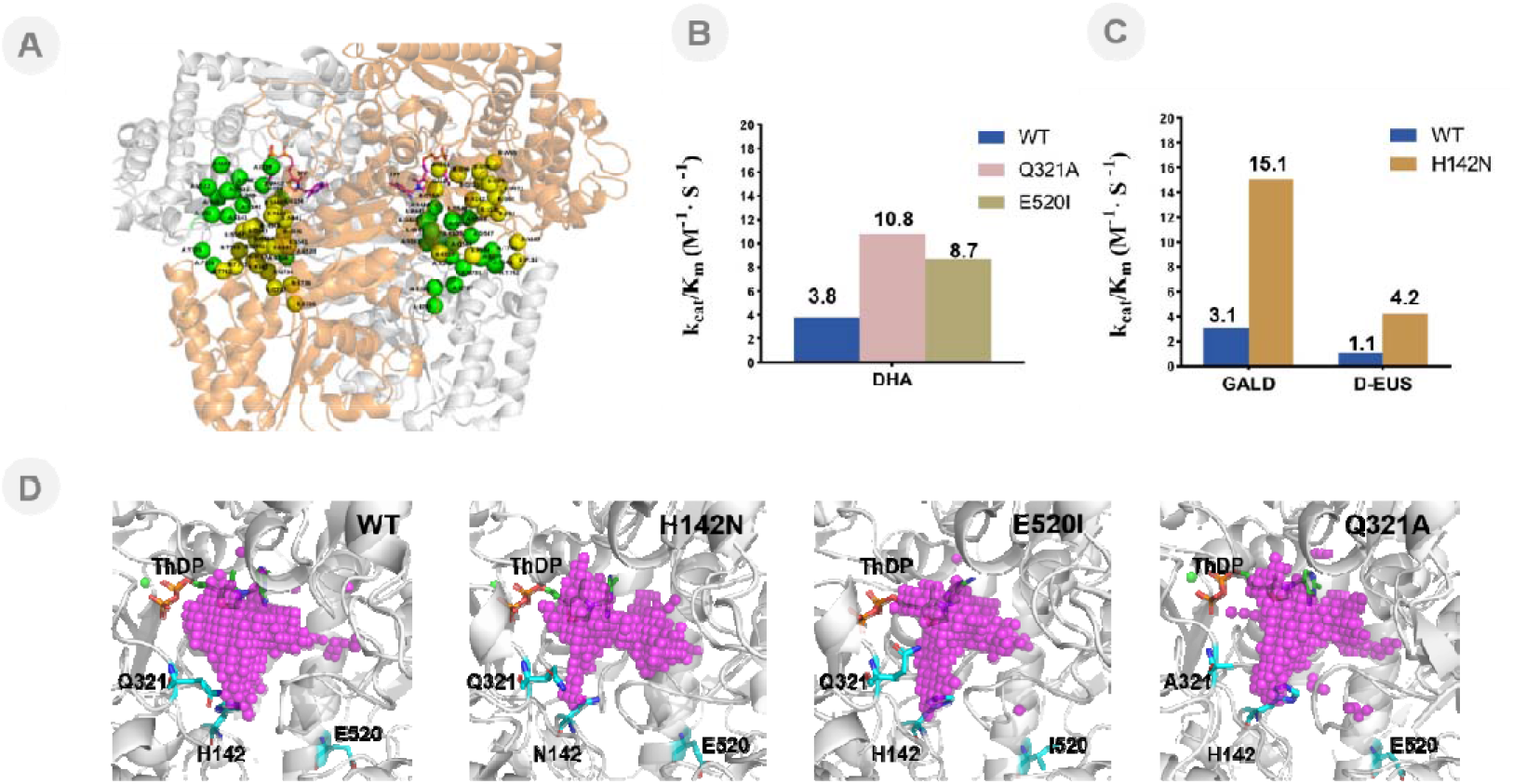
The directed evolution of BbPK. (**A**) The structure model of BbPK and GALD complex. The binding pockets of substrates located at the interface of the BbPK homodimer. The two chains of the BbPK homodimer are indicated in white and orange, respectively. Residues that make up the pockets and key interfacial residues near the pockets are represented by green and yellow spheres based on their chain IDs. The chain ID and residue ID of these residues are labelled. (**B**) The kinetic parameters of wide-type BbPK and mutants for DHA. DHA, dihydroxyacetone. (**C**) The kinetic parameters of wide-type BbPK and mutants for GALD and D-EUS. GALD, glycolaldehyde; D-EUS, D-erythrulose. (**D**) The pocket volumes of WT, H142N, E521I, and Q321A after 50 ns molecular dynamics simulations. Pink dots represent the volumes of the binding pockets, which are 366.6, 269.3, 254.5, and 346.6 Å^3^, respectively. The improved mutants tend to have smaller pockets, which may increase the likelihood of small substrates being in a reactive state. The figures were rendered using Pymol software version 2.3.0. The pockets were calculated using POVME 3.0.

To validate the entire MAG pathway *de novo*, we tested the most abundant hexose D-glucose and pentose D-xylose as examples, which were converted into fructose-6-phosphate (F6P) and xylulose-5-phosphate (Xu5P) after they were transported into cells. Therefore, we used F6P and Xu5P as substrates. F6P and Xu5P are converted into AcP and erythorse-4-phosphate / glyceraldehyde-3-phosphate (E4P/G3P) by BbPK^21^. Two possible pathways could be available for the further conversion of E4P/G3P into AcP according to the order of dephosphorylation and isomerization (Fig. S16). Considering the substrate promiscuity of most phosphatases, it is more efficient to reduce phosphorylated intermediates in the MAG pathway. Thus, E4P was first dephosphorylated and then isomerized into D-EUS (Fig. 3A). However, if G3P was dephosphorylated first, there was no ready enzyme to isomerize glyceraldehyde to DHA^22^. Therefore, we proposed to convert G3P into DHAP and then produce DHA by phosphatase, followed the reaction catalyzed by BbPK (Fig. 3B).

**Fig. 3.**
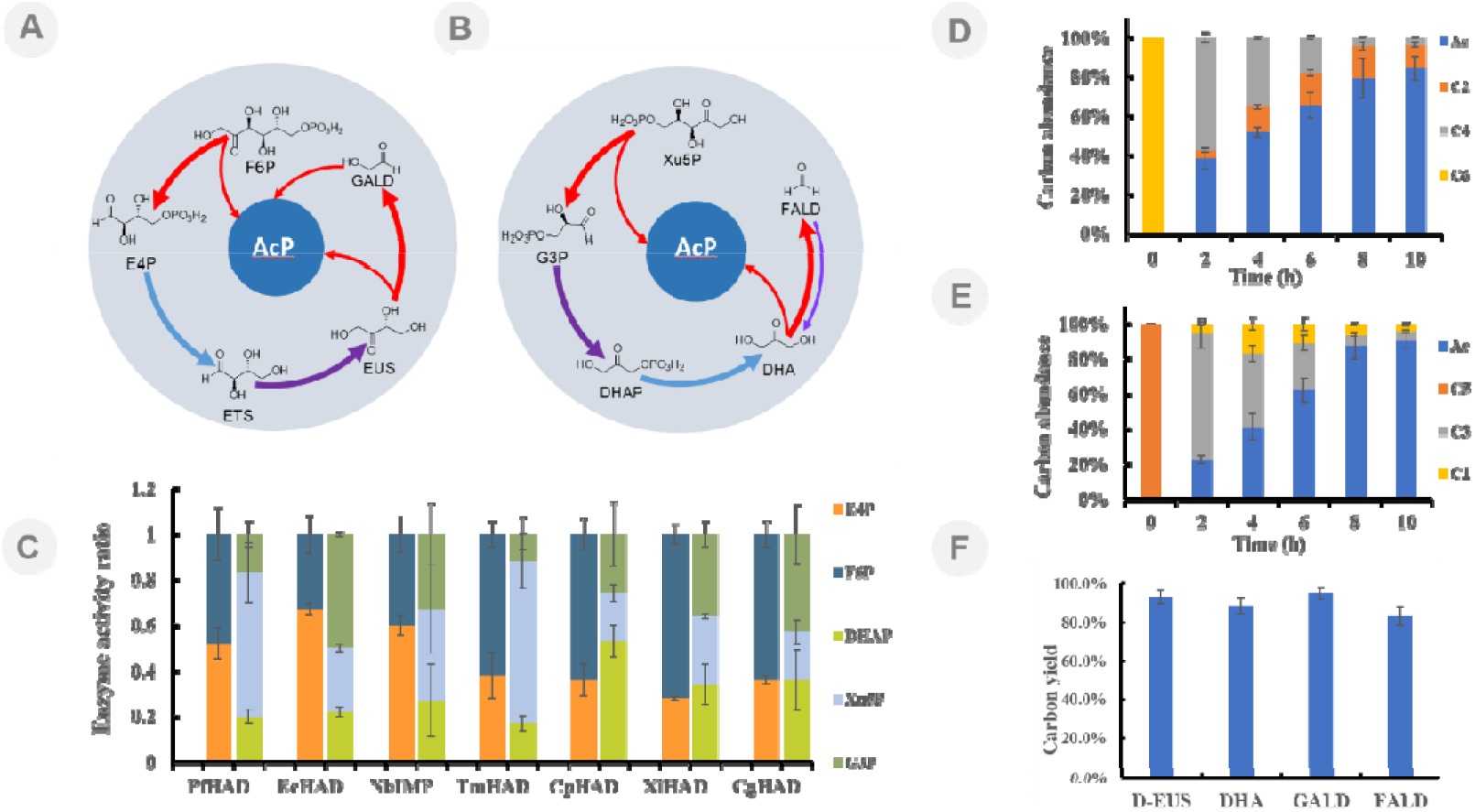
The MAG pathway for utilization of multiple carbon sources *in vitro*. (**A**) The MAG pathway for the synthesis of acetyl phosphate from F6P. (**B**) The MAG pathway for the synthesis of acetyl phosphate from Xu5P. (**C**) The screening of specific phosphatases for E4P and DHAP. Enzyme activity refers to the conversion of substrates in unit time. E4P, D-erythrose-4-phosphate; F6P, fructose-6-phosphate; DHAP, dihydroxyacetone phosphate; Xu5P, D-xylulose-5-phosphate; G3P, D-glyceraldehyde-3-phosphate. Detailed information of phosphatases sees Table S7 and Table S8. (**D**) Metabolite analysis of the MAG pathway for F6P. The starting F6P concentration was 10 mM. Ac, acetic acid; C2, glycoaldehyde; C4 refer to D-erythrose, D-erythrulose, and D-erythrose-4-phosphate. (**E**) Metabolite analysis of the MAG pathway for Xu5P. The starting Xu5P concentration was 10 mM. C1 refer to formaldehyde, C3 refer to D-glyceraldehyde, dihydroxyacetone, dihydroxyacetone phosphate, and D-glyceraldehyde-3-phosphate. Carbon abundance refers to the ratio of carbon moles of an intermediate or product to the total carbon moles. (**F**) The MAG pathway for the utilization of C1, C2, C3, and C4 carbon sources. D-EUS, D-erythrulose; DHA, dihydroxyacetone; GALD, glycoaldehyde; FALD, formaldehyde. The carton yields from C1, C2, C3, and C4 carbon sources are 83%, 95%, 88%, 93%, respectively. Error bars represent s.d. (standard deviation), n = 3.

Considering the multiple phosphate intermediates in the MAG pathway and low selectivity of most phosphatases, it was necessary to find specific phosphatases highly selective for E4P and DHAP. Seven phosphatases from different species were screened (Fig. S17). HAD-like hydrolase (EcHAD) from *Escherichia coli* displayed the best specificity for E4P, with 2-fold activity higher than on F6P (Fig. 3C). Sugar phosphatase from *Candida parapsilosis* (CpHAD) displayed the best specificity on DHAP, with more 2-fold activity higher than on Xu5P and G3P (Fig. 3C). In addition, L-rhamnose isomerase (Ps-LRhI) from *pseudomonas stutzeri* was used to catalyze the isomerization of erythrose^23^, and triose phosphate isomerase (EcTIM) from *E. coli* was used to isomerize G3P.

In addition, ketoses with an even number of carbon atoms could be completely converted into a half of the even number of AcP by PK and isomerase. But ketoses with an odd number of carbon atoms could remain one molecule of formaldehyde at the end, which could result in carbon loss. In order to address this problem, we proposed to introduce formose reaction to condense formaldehyde into glycolaldehyde or DHA^20,24^, which would be further converted into AcP by BbPK (Fig. S18). We compared the catalytic activities of BbPK on glycolaldehyde and DHA. Owing to the higher affinity of BbPK for DHA (Table S4), we chose formolase (FLS) to recycle formaldehyde in the system. With the input of 10 mM DHA, the residual formaldehyde in the system was always lower than 1 mM. Finally, the yield of acetic acid reached 82% (Fig. S19). Thus, it was validated to recycle most of formaldehyde generated in the MAG pathway.

With all available key enzymes, we assembled the MAG pathway *in vitro*. The MAG pathway for F6P included BbPK, EcHAD, and PsLRhI, while it consisted of BbPK, EcTIM, CpHAD, and FLS for Xu5P (Fig. S20). Under the optimal reaction conditions, the final yields of acetic acid from F6P and Xu5P reached 84% and 90%, respectively (Fig. 3D and 3E), which were higher than that of the glycolytic pathway^13^. Metabolite analysis showed that F6P and Xu5P were quickly cleaved by BbPK within the first two hours (Fig. 3D and 3E), while the conversion of E4P or G3P to AcP was slow due to the low catalytic efficiency of BbPK on glycolaldehyde, DHA, and D-EUS (Table S4). After 10 hours, there were still small amounts of glycolaldehyde and D-EUS remaining in the F6P reaction system, as well as formaldehyde and DHA in the Xu5P reaction system. Furthermore, we also tested the MAG pathway for C1, C2, C3, and C4 carbon source *in vitro* (Fig. S21). The results demonstrated that the MAG pathway enabled to achieve the nearly stoichiometric synthesis of acetyl-CoA from all tested carbon sources, where the highest yield of acetic acid reached 95% (Fig. 3F).

Lastly, we evaluated the feasibility of the MAG pathway *in vivo* in an engineered *E. coli* stain, in which the influence of the Embden-Meyerhof-Parnas pathway (EMP) was mitigated (Fig. 4A)^25^: the *ldhA, frdBC, pflB* and *adhE* genes were knocked out to abolish the consumption of pyruvate^26^; the *zwf* and *epd* genes were knocked out to avoid the consumption of G6P and E4P^27,28^; the *fucO, yqhD* and *aldA* genes were knocked out to block the consumption of glycolaldehyde^29-31^; the pathways that consume DHAP, DHA and formaldehyde (encoded by *glpD, gldA* and *frmABR*) were also blocked; the key enzyme MgsA was blocked to prevent the conversion of G3P into pyruvate^32^; the *acs* and *ptaI* genes were knocked out to prevent the conversion of acetate and acetyl phosphate into acetyl-CoA from the EMP pathway^33,34^. After an initial aerobic growth phase for cell growth and protein induction, anaerobic conditions were used to further prevent the intermediates from being oxidized to CO_2_. Our results showed that the carbon yield of acetic acid from glucose reached 82%, which was significantly higher than that of glycolytic pathway (Fig. 4B). These results indicated that the MAG pathway worked well *in vivo*.

**Fig. 4.**
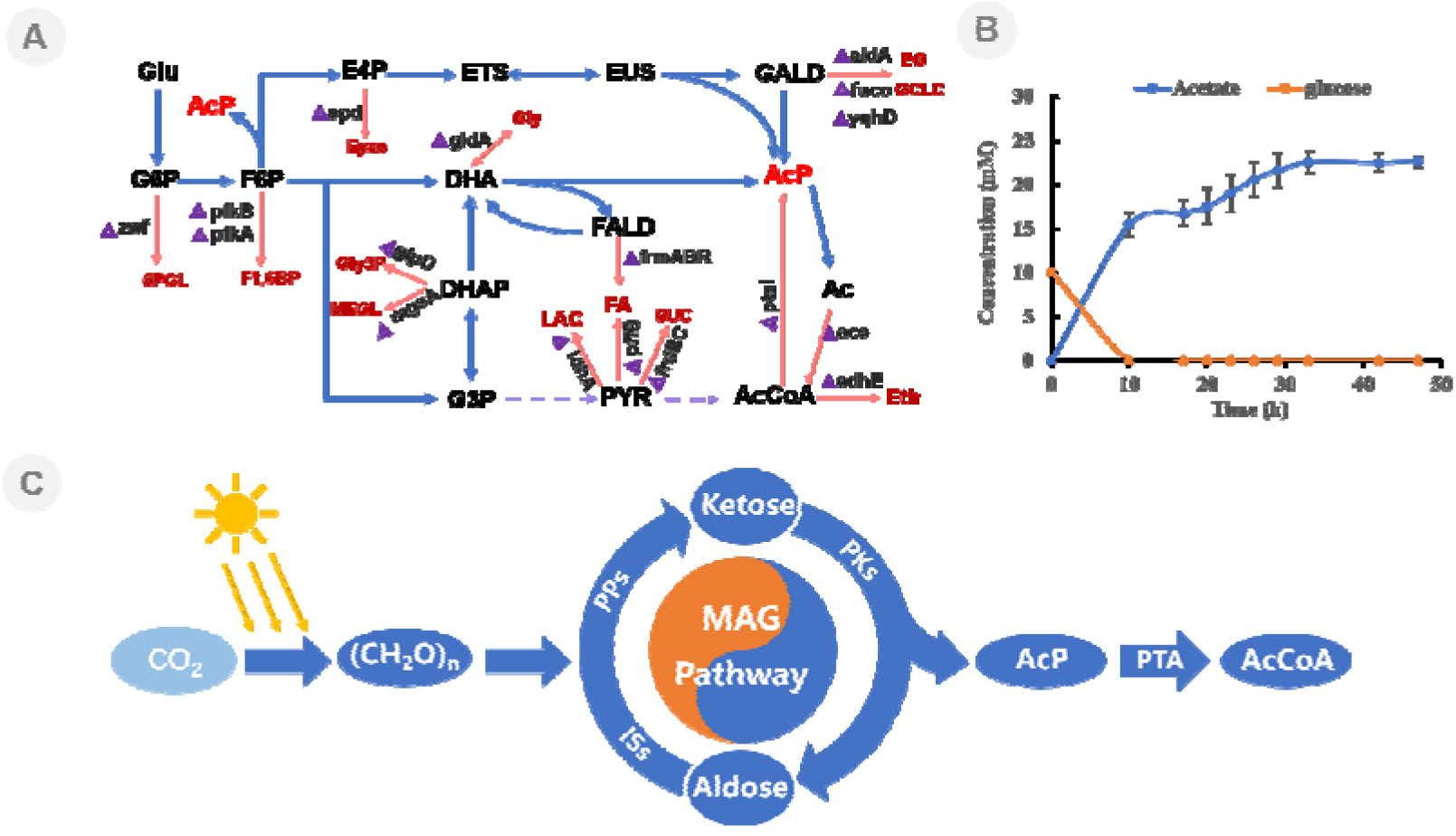
The implementation of MAG pathway *in vivo*. (**A**) Metabolic engineering strategy for MAG pathway in *E. coli* strains. The red arrows represent the pathway that need to be blocked, and the related genes are marked. Blue arrows represent the MAG pathway. Glu, glucose; G6P, D-glucose-6-phosphate; F6P, D-fructose-6-phosphate; E4P, D-erythrose-4-phosphate; ETS, D-erythrose; EUS, D-erythrulose; GALD, glycoaldehyde; DHA, dihydroxyacetone; DHAP, dihydroxyacetone phosphate; G3P, D-glyceraldehyde-3-phosphate; PYR, pyruvate; Ac, acetic acid; 6PGL, 6-phospho-gluconic acid; F1,6BP, D-fructose-1,6-diphosphate; Eyac, erythronic acid; Gly, glycerin; EG, ethylene glycol; GCLC, glycolic acid; Gly3P, glycerol-3-phosphate; MEGL, methylglyoxal; LAC, lactic acid; SUC, succinic acid; FA, formic acid; Eth, ethanol; AcP, acetyl phosphate. (**B**) Engineered strains for acetic acid production from glucose via MAG pathway. (**C**) Conceptual diagram of the conversion of CO_2_ to AcCoA via the MAG pathway. “(CH_2_O)_n_” refer to carbohydrate. Iss, isomerase; PPs, phosphatase. Error bars represent s.d. (standard deviation), n = 3.

Depending on the versatile PK, it is available to convert various carbon sources to AcP and then generate acetyl-CoA (Fig. S2). However, in nature the PK mainly cooperates with acetate kinase (AK), and even forms gene clusters that convert AcP into acetate and ATP for energy supply (Fig. S22 and S23)^35,36^. Since the intricate sugar molecules produced through formose reactions provided an important source of carbohydrate for early life^37-39^, the ability to catabolize various carbohydrates by PK might have been of great significance for life survival during the ancient reductive world. Therefore, until now most of PKs still retained these ancient functions (Fig. 1C). However, with the advent of photosynthesis and oxygen accumulation, the main carbon sources changed in the world and the energy production efficiency of PK pathway is much lower than that of the aerobic respiration process, resulting in the loss of PK genes in most of species (Fig. 1B). By reviving the ancient function of PK on different ketoses, the MAG pathway provides the simplest way for carbohydrates metabolism (Fig. 4C), which is not only advantageous for the utilization of complex carbon sources, but also has an important advantage in terms of atom economy compared with other pathways (Fig. S24 and Table S5). The design, build and test of MAG pathway indicates the possibility of using multiple carbon sources with higher efficiency for biomanufacturing in the future.

## Acknowledgments

This work was supported by National Key R&D Program of China (grant no. 2018YFA0901600) (Y.W.L) and National Key R&D Program of China (grant no. 2021YFC2103500) (Y.W.L) and Strategic Priority Research Program of the Chinese Academy of Sciences-Precision Seed Design and Breeding (grant no. XDA24020103-3) (H.F.J) and Tianjin Synthetic Biotechnology Innovation Capacity Improvement Project (grant no. TSBICIP-KJGG-007) (H.F.J) and Tianjin Outstanding Scholar Program (Y.H.M). We thank YH. Yao for GC-MS analysis. We thank the core facility center at Tianjin Institution of Industrial Biotechnology, CAS, for instrument and technology support.

## Author contributions

H.F.J, Y.H.M, Y.Q.Y, and Y.W.L designed the experiments. Y.Q.Y, Y.W.L, J.Z, Q.Y.Y, H.Y.C, H.D.Z, M.T.L, J.C, D.Y.L, X.Y.L, X.S performed the experiments. Y.Q.Y and Y.W.L processed data. Y.Q.Y, Y.W.L, and H.F.J wrote the manuscript. Y.H.M, H.F.J, Y.W.L, J.C, Y.H.Z reviewed the manuscript.

## Competing interests

There is no competing financial interest

## Notes

### Competing Interest Statement

The authors have declared no competing interest.

