## supplemental figure S1-S24 for "Designing a Minimal Artificial Glycolytic Pathway"

**Materials and Methods**

**Quantum-chemical analysis for PK from *Bifidobacterium* (BbPK)**

The computational model was obtained using AlphaFold^1^. The complex of BbPK with ThDP and substrate was generated with PyMOL^2^. The model contains 218 atoms with a total charge of +1, including the side chains of His64, His553, Glu479, Tyr501, Gln321, Ser440, His142, Gly155, His320, Asn549, Gln546, His548, His97, and the cofactor ThDP. Generally, the glutamate (Glu479) is modeled in protonated state for forming a hydrogen bond with the N1’ atom of ThDP. His97 was considered as the most possible candidate of proton donor for dehydration process, and it was modeled in the doubly protonated state. According to the interaction mode of pocket residues, His64, His553, His320, His142, His548, were modeled in their singly protonated states. The structural models of short chain ketoses were obtained from the PubChem.

All the calculations were performed using the Gaussian 09 package^3^ and the B3LYP method. The 6-31G (d, p) basis set was used for the geometry optimizations, and the electronic energies of the stationary points were refined by single-point calculations with the 6-311++G (2d, 2p) basis set. Solvation energies were calculated with the SMD^4^ implicit solvent method and a dielectric constant of ε = 4. Previous studies have shown that the effect of the solvation diminishes rapidly with the size of the active site model, rendering the particular value used for the dielectric constant less critical. The zero-point energy corrections were done at the same level of theory as the geometry optimizations.

As used in the cluster approach^5,6^, a number of atoms were kept fixed to their crystallographic positions in the geometry optimizations (indicated by asterisks in the Fig. S7). This coordinate-fixing protocol is very important to avoid large unrealistic movements of the various groups at the active site. This approach deals only with the chemical steps of the enzymatic reactions, implying that the substrate binding and product release are usually not explicitly considered by the calculations. Therefore, an implicit assumption in the current model is that neither of these events are rate- or selectivity-determining.

**Molecular docking of substrate to BbPK**

The molecular docking tool of GNINA^7^ was used to predict the potential interaction of different ligands (DHAP, Eu4P, Xu5P, F6P, D/L-EUS, and DHA) with PK from *Bifidobacterium*. A structural model of BbPK was obtained using AlphaFold^1^ and a 20-nanosecond molecular dynamics simulation was performed to optimize the side chain conformations. The refined PK structural model was treated as the receptor for docking. The structural models of the ligands and ThDP were obtained from the PubChem. The binding site was determined by the structural alignment to the crystal structure of a known phosphoketolase (PDB ID：3AHE)^8^. The docking models were ranked by GNINA’s CNN pose score. For each ligand, the top 1 docking model was selected and energy minimization was performed by OpenMM7^9^ with the Amber14^10^ forcefield to optimize the complex model.

**Phosphoketolase selection**

To investigate the function of PK genes in distant evolutionary branches, we screened and analyzed all potential PKs in the NCBI database. First, we predicted all potential PKs by search against the non-redundant database with the Pfam domain ID PF03894 (hmmscan --cpu 10 --domtblout output.txt -E 1e-4 PF03894.hmm NR.fasta)^11^. Second, we retrieved all PKs from KEGG database (https://www.genome.jp/entry/pf:xfp). Third, we made a local blastp search using PKs from NCBI as the query sequences, and PKs from KEGG as the BLAST database. After blastp search, 12185 potential PKs were screened with three standards: the best hit is a D-xylulose 5-phosphate/D-fructose 6-phosphate phosphoketolase (XFP, EC:4.1.2.9 4.1.2.22, KO: K01621), the identity is more than 40, and the align length is more than 600. Fourth, all PKs were classified into 4101 groups by using OrthoMCL with the amino acid identity more than 90 in a group. For each group, we selected a PK gene, which is closest approximation to supposed optimal sequence, consisting of the highest frequency residues in multiple sequences alignment. Using the similar strategy, 23 PKs were screened based on the standard of the identity of 60. Finally, we totally synthesized 7 PK genes (Firmicutes: WP_125748362; Fungi: RDK44081; Gammaproteobacteria: WP_154224703; Actinobacteria: WP_011743105; Alphaproteobacteria: WP_038497079; Chloroflexi: WP_008479084; Alphaproteobacteria: WP_156359173) for next functional evaluation.

**Protein engineering of BbPK**

To obtain full mutations, oligonucleotide primers were designed with the degenerate codon NNK, which cover almost all mutations with only 96 clones. Hence, a total of 3264 clones were screened against 34 single-site saturation mutation libraries. Each single-site saturation mutant library was generated based on PCR. The PCR product was degraded the template with *Dpn*I restriction endonuclease, and then transformed into *E. coli* BL21 (DE3) competent cells for library construction. Each colony was incubated in 200 μL of LB medium for 24 hours to plateau at 37 °C and then transferred to 1 mL of the same medium for protein expression. The cells, which induced by IPTG (isopropyl-β-D-thiogalactopyranoside) and cultured overnight at 16 °C, were harvested by centrifugation. The bacterial pellet was washed and resuspended in reaction buffer (50 mM potassium phosphate buffer, 5 mM GALD or 20 mM DHA, pH 7.4). After 3 hours of reaction at 37 °C, the supernatant was collected by centrifugation for detection of substrate or product.

For GALD, we determined the activity of the mutants by detecting the reduction of substrate. The detection method was as follows: Added 120 μL chromogenic reagent to 60 μL sample and heated at 90 °C for 15 minutes. Subsequently, the residual substrate concentration was measured spectrophotometrically at 650 nm. The chromogenic reagent: 1.5 g diphenylamine was dissolved in 100 mL acetic acid, and then 1.5 mL pure sulfuric acid was added.

For DHA, we screened for highly active mutants by detecting the product formaldehyde. The formaldehyde detection method was as follows: 40 μL sample was mixed with 160 μL chromogenic reagent, and then heated at 60 °C for 10 minutes. Subsequently, formaldehyde production was measured spectrophotometrically at 440 nm. 100 mL chromogenic reagent (pH 6.0) contains 25 g ammonium acetate, 3 mL acetic acid, and 0.25 mL acetylacetone solution.

**Activity assay and kinetic properties of PK and mutants**

The standard reaction mixture (100 µL) contained 50 mM potassium phosphate buffer (pH 7.5), 5 mM MgSO_4_, 1 mM ThDP, 10 mM GALD (DHA or D-EUS), 1 mM ADP, 0.2 mg mL^-1^ AckA, 5 U hexokinase, 2.5 U Glucose-6-phosphate dehydrogenase, 1mM NADP^+^, and 10 mM glucose. 0.5mg mL^-1^ PK was added into the reaction system. The reactions conducted at 37 °C. NADPH was detected spectrophotometrically at 340 nm. Enzyme kinetics with GALD (DHA or D-EUS) as substrate were determined in assays with GALD (DHA or D-EUS) concentrations of 0-110 mM. Kinetic parameters k_cat_ and K_m_ were determined by measuring the initial velocities of the enzymic reaction and curve-ﬁtting according to the Michaelis-Menten equation, using GraphPad Prism 5 software.

**Investigation of the distribution of PK, AK, and PTA genes**

To comprehensively investigate the distribution of PK, AK, and PTA genes in the tree of life, we respectively identified PK, AK, and PTA genes in about 50,000 sequenced genomes collected in NCBI database. Using the similar method for PK selection, we totally identified 12185 PKs, 29003 AKs, and 21686 PTAs in all sequenced genomes. Furthermore, we downloaded the gff files of all sequenced genomes from NCBI to detect the gene cluster between three classes of genes. Only the two or three genes were closely adjacent to each other in the same chromosome, or the number of interval genes between two genes were less than 4, the two or three genes were considered as the potential gene cluster.

**Chemicals and agents**

Common chemicals were bought from Sigma-Aldrich (Shanghai, China), SolarBio (Beijing, China), Zhenzhun Biotech (Shanghai, China) and Yuanye Biotech (Shanghai, China). Standard acetic acid, formaldehyde (FALD), glycoaldehyde (GALD), dihydroxyacetone (DHA), L-erythrulose (L-EUS), D-erythrulose (D-EUS), D-xylulose, D-fructose, and acetyl phosphate (AcP) were purchased from Yuanye Biotech (Shanghai, China). Fructose-6-phosphate (F6P), D-xylulose-5-phosphate (Xu5P), D-erythrose-4-phosphate (E4P), D-glyceraldehyde (GCD), and dihydroxyacetone phosphate (DHAP) were purchased from Zhenzhun Biotech (Shanghai, China). Restriction enzymes and DNA polymerase were purchased from Thermo Fisher Scientific (Shanghai, China), New England Biolabs and TransGen Biotech (Beijing, China). Kits for DNA manipulation were purchased from Axygen (Shanghai, China) and TransGen Biotech (Beijing, China). Alkaline phosphatase was purchased from TaKaRa (Dalian, China). Primers and synthesized genes were obtained from Genecreat (Wuhan, China) or GENEWIZ (Suzhou, China). Materials and equipment for protein purification were obtained from GE Healthcare (Beijing, China) and BioRad (Beijing, China).

**Bacterial strains and growth condition**

*Escherichia coli* *DH5α* strain (TransGen^TM^) was grown at 37 ℃ in LB medium for gene cloning and other DNA manipulations. *E. coli* BL21 (DE3) (TransGen^TM^) was grown at 37 ℃ or 16 ℃ in 2YT medium for protein expression. Antibiotics for selection purposes were used with 100 μg mL^-1^ spectinomycin, 100 μg mL^-1^ ampicillin, 34 μg mL^-1^ chloramphenicol or 100 μg mL^-1^ kanamycin.

**Activity assay of enzymes**

**Acetate kinase (AckA, EC 2. 7. 2.1)**

The activity of AckA was determined by monitoring the product ATP^12^, which is detected by coupling hexokinase and glucose-6-phosphate dehydrogenase. Hexokinase converts ATP and glucose to G6P and ADP. Glucose-6-phosphate dehydrogenase converts G6P and NADP^+^ to 6-phosphate gluconate and NADPH. NADPH was detected spectrophotometrically at 340 nm. The standard reaction mixture (100 µL) contained 50 mM potassium phosphate buffer (pH 7.5), 5 mM MgSO_4_, 1 mM ThDP, 5 mM AcP, 1 mM ADP, 0.2 mg mL^-1^ AckA, 0.5 U hexokinase, 0.25 U glucose-6-phosphate dehydrogenase, 1mM NADP^+^ and 10 mM glucose.

**Erythrose isomerase (Ps-LRHI, EC 5. 3. 1. 14)**

The activity of Ps-LRHI was determined by monitoring the product erythrulose, which was detected by HPLC. The standard reaction mixture (100 µL) contained 50 mM potassium phosphate buffer (pH 7.5), 5 mM MgSO_4_, 10 mM erythrose, and 1 mg mL^-1^ Ps-LRHI. HPLC conditions: column, Aminex HPX-87H (Bio-Rad); detection wavelength, 277 nm; mobile phase, 5 mM sulphuric acid; flow rate, 0.6 mL min^-1^; sample volume, 20 µL; column temperature, 40 ℃.

**Formolase (FLS, EC 4. 1. 2. -)**

The activity of FLS was determined by monitoring the product dihydroxyacetone, which was detected by HPLC. The standard reaction mixture (100 µL) contained 50 mM potassium phosphate buffer (pH 7.5), 5 mM MgSO_4_, 1 mM ThDP, 30 mM formaldehyde, and 1mg mL^-1^ FLS. The reactions were conducted at 37 ℃ for 2 hours and were terminated by adding 100 μL acetonitrile. HPLC conditions: column, Aminex HPX-87H (Bio-Rad); detection wavelength, 200 nm; mobile phase, 5 mM sulphuric acid; flow rate, 0.6 mL min^-1^; injection volume, 20 µL; column temperature, 40 ℃.

**Glycoaldehyde synthase (GALS, EC 4. 1. 2. -)**

The activity of GALS was determined by monitoring the product GALD, which was detected by HPLC. The standard reaction mixture (100 µL) contained 50 mM potassium phosphate buffer (pH 7.5), 5 mM MgSO_4_, 1 mM ThDP, 30 mM formaldehyde, and 1mg mL^-1^ GALS. The reactions were conducted at 37 ℃ for 2 hours and were terminated by adding 100 µL acetonitrile. HPLC conditions: column, Aminex HPX-87H (Bio-Rad); detection wavelength, 200 nm; mobile phase, 5 mM sulphuric acid; flow rate, 0.6 mL min^-1^; injection volume, 20 µL; column temperature, 40 ℃.

**Triose-phosphate isomerase (TIM, EC 5. 3. 1. 1)**

The activity of TIM was determined by monitoring the product DHAP, which is detected by coupling with glycerophosphate dehydrogenase. NAD^+^ was detected spectrophotometrically at 340 nm. The standard reactions contained 50 mM potassium phosphate buffer (pH 7.5), 0.05 mg mL^-1^ TIM, 1U glycerophosphate dehydrogenase, 1 mM NADH, and 5mM glycerophosphate.

**Erythrose-4-phosphate isomerase (RpiB, 5. 3. 1. 34)**

The activity of RpiB was determined by monitoring the product D-erythrulose-4-phosphate, which was detected by coupling with alkaline phosphatase. D-EUS was detected by HPLC. The standard reactions contained 50 mM potassium phosphate buffer (pH 7.5), 5 mM MgSO_4_, 0.5 mg mL^-1^ RpiB, and 10 mM erythrose-4-phosphate. The reaction system was terminated by heating at 95 °C for 5 minutes, and then cool down to 37 °C. Adding 5 U alkaline phosphatase to above reaction system and maintained at 37 ℃ for 4 hours. 100 µL acetonitrile was used to terminate the reaction. D-EUS was detected by HPLC.

HPLC conditions: column, Aminex HPX-87H (Bio-Rad); detection wavelength, 277 nm; mobile phase, 5 mM sulphuric acid; flow rate, 0.6 mL min^-1^; injection volume, 20 µL; column temperature, 40 ℃.

**Erythrose-4-phosphate dehydrogenase (E4PDH, 1. 2. 1. 72)**

The activity of E4PDH was determined by monitoring the product NADH, which was detected spectrophotometrically at 340 nm. The standard reactions contained 50 mM potassium phosphate buffer (pH 7.5), 0.1 mg mL^-1^ E4PDH, 5 mM E4P, and 1 mM NAD^＋^.

**Enzyme activity of phosphoketolase from *Bifidobacterium* for DHA, L-EUS and D-EUS**

The coding gene of phosphoketolase from *Bifidobacterium* was ligated into the expression vector pET-28a via NdeІ and XhoI restriction sites. *E. coli* BL21(DE3) cells carrying recombinant plasmid were inoculated into 5 mL LB (Luria Broth) medium with Kanamycine (100 μg mL^-1^) and cultured overnight at 37 °C, and then scaled up to 800 mL 2YT medium (16 g L^-1^ Tryptone, 10 g L^-1^ yeast extract, 5 g L^-1^ NaCl) containing Kanamycine (100 μg mL^-1^). Gene expression was induced by adding IPTG (isopropyl-β-D-thiogalactopyranoside) to a final concentration of 0.5 mM when OD_600_ reached 0.6. The cell cultures continued to grow overnight at 16 °C before being harvested by centrifugation at 6,000×g and then was resuspended in 50 mL lysis buffer (50 mM potassium phosphate buffer (pH 7.4), 5 mM MgSO_4_, 0.5 mM ThDP). The bacterial pellet was lysed by using a high-pressure homogenizer (JNBIO, China), and the cell debris was removed by centrifugation at 10,000×g for 60 mins at 4 °C. The soluble protein sample was loaded onto a nickel affinity column (GE Healthcare), which was rinsed with 50 mL wash buffer (50 mM potassium phosphate buffer (pH 7.4), 5 mM MgSO_4_, 0.5 mM ThDP, and 50 mM imidazole) and then eluted with 20 mL elution buffer (50 mM potassium phosphate buffer (pH 7.4), 5 mM MgSO_4_, 0.5 mM ThDP, and 200 mM imidazole). The eluted protein was concentrated and dialyzed against lysis buffer (50 mM potassium phosphate buffer (pH 7.4), 5 mM MgSO_4_, and 0.5 mM ThDP) by ultrafiltration with an Amicon Ultra centrifugal filter device (Millipore, USA) with a 30 kDa molecular-weight cutoff. The protein concentration was determined using a BCA Protein Assay Reagent Kit (Pierce, USA) with BSA as the standard.

Activity of PKs on DHA was determined with 1 mg mL^-1^ purified recombinant protein in 200 µL reaction mixtures. The reaction system comprised 10mM DHA, 1 mM ThDP, 5 mM MgSO_4_, and 50 mM phosphate buffer (pH 7.4). After incubation at 37 ℃ for 0.5 hours, the reaction was stopped by adding 200 µL of acetonitrile. Acetyl-phosphate in samples is determined by chromogenic and liquid chromatography, the by-product formaldehyde was confirmed by GC-MS.

Chromogenic detection of acetyl-phosphate (AcP): add 100 μL hydroxylamine hydrochloride solution (2M, pH 6.5) to 100 μL sample, conduct at 30 ℃ for 10 mins. Then add 200 μL of chromogenic solution, which is prepared with 10 mL FeCl_3_.6H_2_O (FeCl_3_.6H_2_O powder dissolved in 0.1 M hydrochloric acid, 5% (m/v)), 10 mL trichloroacetic acid (15% (m/v)), and 10 mL HCl (4 M), conduct at 30 ℃ for 5 mins, then detect the absorbance at 505 nm.

HPLC detection for AcP: add equal volume of 5% sulfuric acid to the sample for completely decomposing AcP into acetic acid. Acetic acid was then detected by HPLC. HPLC conditions: column, Aminex HPX-87H (Bio-Rad); detection wavelength, 210 nm; mobile phase, 5 mM sulphuric acid; flow rate, 0.6 mL min^-1^; sample volume, 20 µL; column temperature, 40 ℃.

Formaldehyde detection by GC-MS. Sample derivatization：100 μL 2.4-dinitrophenylhydrazine solution was added to 100 μL samples, the mixture was performed at 60 °C for 60 mins in the dark. Then added 400 μL of n-hexane to the mixed solution, centrifuge at 5000×g for 2 minutes. Separated the upper solution and then added appropriate amount of anhydrous sodium sulfate powder, centrifuged at 10000×g for 10 minutes. Separated the upper solution, and was detected by GC-MS. GC-MS conditions: Electron ionization (EI) GC-MS analyses were performed with a model 7890A GC (Agilent) with a DB-5 fused silica capillary column (30 cm length, 0.25 mm inner diameter, 0.25 μm film thickness) coupled to an Agilent 7200 Q-TOF mass selective detector; Injections were performed by a model 7683B autosampler. The GC oven was programmed from 160 ℃ (held for 1 min) to 240 ℃ at 10 ℃ min^-1^, and then held for 5 mins; the injection port temperature was 250 ℃, and the transfer line temperature was 280 ℃. The carrier gas, ultra-high purity helium, flowed at a constant rate of 1.2 mL min^-1^. For full-scan data acquisition, the MS scanned from 35 to 550 atomic mass units. Data analysis for GC-MS was performed with Mass Hunter software (Agilent, USA) and NIST Database.

The activity of PKs on L-EUS and D-EUS were determined with 1 mg mL^-1^ purified recombinant protein in 200 µL reaction mixtures. The reaction system comprised 10 mM L-EUS or D-EUS, 1 mM ThDP, 5 mM MgSO_4_, and 50 mM phosphate buffer (pH 7.4). After incubation at 37 ℃ for 0.5 hours, the reaction was stopped by adding 200 µL of acetonitrile. AcP in samples was detected by chromogenic and liquid chromatography. The by-product glycoaldehyde was confirmed by GC-MS. Chromogenic and liquid chromatography detection of AcP were same as DHA.

Glycoaldehyde was confirmed by GC-MS. Samples were freeze-dried, then 60 μL of 0.2 M PFBOA solution was added and mixed, the reaction was performed at 30 °C for 90 minutes. 300 μL of n-hexane was added and centrifuge at 5000×g for 2 minutes. Separated the upper solution and then added appropriate amount of anhydrous sodium sulfate powder, centrifuged at 10000×g for 10 minutes. Separated 100 μL upper solution，then 30 μL N-Methyl-N-(trimethylsilyl)-trifluoroacetamide with 1% trimethylchlorosilane was added and mixed, the reaction system conducted at 30 °C for 30 minutes. The product was detected by GC-MS. GC-MS conditions: Electron ionization (EI) GC-MS analyses were performed with a model 7890A GC (Agilent) with a DB-5 fused silica capillary column (30 cm length, 0.25 mm inner diameter, 0.25 μm film thickness) coupled to an Agilent 7200 Q-TOF mass selective detector; Injections were performed by a model 7683B autosampler. The GC oven was programmed from 60 ℃ (held for 1 min) to 100 ℃ at 5 ℃ min^-1^, to 300 ℃ at 25 ℃ min^-1^ and then held for 5 mins; the injection port temperature was 250 ℃, and the transfer line temperature was 280 ℃. The carrier gas, ultra-high purity helium, flowed at a constant rate of 1.2 mL min^-1^. For full-scan data acquisition, the MS scanned from 35 to 550 atomic mass units. Data analysis for GC-MS was performed with Mass Hunter software (Agilent, USA) and NIST Database.

Activity of PKs on E4P, D-ETS, D-GCD and DHAP were determined with 1 mg mL^-1^ of purified recombinant protein in 200 µL reaction mixtures. The reaction system included substrate (10 mM E4P, D-ETS, D-GCD, or DHAP), 1 mM ThDP, 5 mM MgSO_4_, and 50 mM phosphate buffer (pH 7.4). After incubation at 37 ℃ for 0.5 hours, the reaction was stopped by adding 200 µL of acetonitrile. AcP in samples was detected by chromogenic and liquid chromatography. Chromogenic and liquid chromatography detection of AcP were same as DHA.

Activity of PKs on erythrulose-4-phosphate (Eu4P) were determined with 1 mg mL^-1^ purified recombinant protein in 200 µL reaction mixtures. The reaction system included 10 mM Eu4P, 0.5 mg mL^-1^ RpiB, 1 mM ThDP, 5 mM MgSO_4_, and 50 mM phosphate buffer (pH 7.4). After incubation at 37 ℃ for 0.5 hours, the reaction was stopped by adding 200 µL of acetonitrile. AcP in samples was detected by chromogenic and liquid chromatography. Chromogenic and liquid chromatography detection of AcP were same as DHA.

**Expression, purification, and enzyme kinetics of PKs** **from different species**

The PKs coding genes from different species were ligated into the expression vector pET-28a via *Nde*І and *Xho*I restriction sites. All genes were expressed in BL21 (DE3) and purified on the Ni-NTA column. Large-scale purification (800 mL) typically produced about 50 mg enzyme. The protein concentration was determined using the BCA Protein Assay Reagent Kit (Pierce, USA) with BSA as the standard.

Determination of kinetics of PKs on GALD, DHA, L-EUS, D-EUS. The standard reaction mixture (200 µL) contained 50 mM potassium phosphate buffer (pH 7.5), 5 mM MgSO_4_, 1 mM ThDP, 10 mM GALD (or, DHA, L-EUS, D-EUS), 1 mM ADP, 0.2 mg mL^-1^ AckA, 1 U hexokinase, 0.5 U glucose-6-phosphate dehydrogenase, 1mM NADP^+^, and 10 mM glucose. Various PKs (0.25-1 mg mL^-1^) from different species were added into the reaction system. The reactions conducted at 37 ℃. The production of NADPH was detected at 340 nm. Enzyme kinetics with DHA, L-EUS, and D-EUS as substrates were determined in assays with concentrations of 0-110 mM. Kinetic parameters were determined from triplicate experiments using GraphPad Prism 5 (GraphPad Software, USA).

Determination of kinetics of PKs on Xu5P. The standard reaction mixture (200 µL) contained 50 mM potassium phosphate buffer (pH 7.5), 5 mM MgSO_4_, 1 mM ThDP, 5 mM Xu5P, 0.05 mg mL^-1^ TIM, 1U glycerophosphate dehydrogenase, and 1 mM NAD^＋^. Various PKs (0.1mg mL^-1^) from different species were added into the reaction system. The reactions conducted at 37 ℃. The production of NADH was detected at 340 nm.

Determination of kinetics of PKs on F6P. The standard reaction mixture (200 µL) contained 50 mM potassium phosphate buffer (pH 7.5), 5 mM MgSO_4_, 1 mM ThDP, 5 mM F6P, 0.3 mg mL^-1^ erythrose-4-phosphate dehydrogenase, and 1 mM NAD^＋^. Various PKs (0.1mg mL^-1^) from different species were added into the reaction system. The reactions conducted at 37 ℃. The production of NADPH was detected at 340 nm.

**Expression, purification, and enzyme kinetics of phosphatases from different species**

The phosphatases coding genes of different species were ligated into the expression vector pET-28a via NdeІ and XhoI restriction sites. All genes were expressed in BL21 (DE3) and purified on the Ni-NTA column. Large-scale purification (800 mL) typically produced about 5–50 mg enzyme. The protein concentration was determined using a BCA Protein Assay Reagent Kit (Pierce, USA) with BSA as the standard.

The kinetics of phosphatases on F6P, Xu5P, E4P, G3P, and DHAP were determined by monitoring the production of fructose, xylulose, ETS, GCD and DHA by HPLC. The standard reaction mixture (100 µL) contained 50 mM potassium phosphate buffer (pH 7.5), 5 mM F6P (or Xu5P, E4P, G3P, DHAP). Various phosphatases (0.1mg mL^-1^) from different species were added into the reaction system. The reactions conducted at 37 ℃ for 0.5-2 hours. 100 μL acetonitrile was added to the samples to terminate the reactions, and then analyzed by HPLC.

HPLC condition for GCD or DHA: column, Aminex HPX-87H (Bio-Rad); detection wavelength, 210 nm; mobile phase, 5 mM sulphuric acid; flow rate, 0.6 mL min^-1^; sample volume, 20 µL; column temperature, 40 ℃.

HPLC detection for fructose, xylulose, and erythrose. Sample derivatization: Reaction samples (50 µL) were mixed with a solution of O-benzylhydroxylamine hydrochloride (50 µL, 0.14 mmol mL^-1^) (pyridine: methanol: water = 33:15:2). After incubation at 50 ℃ for 60 mins, samples were diluted in methanol (100 µL) and directly analyzed by HPLC chromatography. HPLC condition: column, X-BridgeTM C18, 5 μm, 4.6×250 mm column from Waters (Milford, USA); sample volume, 30 μL; solvent system, (A) aqueous trifluoroacetic acid (TFA) (0.1 % (v/v) and (B) TFA (0.095% (v/v)) in CH_3_CN/H_2_O (4:1), gradient elution from 20-60 % B in 16 mins; flow rate, 1 mL min^-1^; detection wavelength, 215 nm; column temperature, 35 ℃.

**Formaldehyde circulation system verification**

Comparison of PK-GALS formaldehyde recycling system and PK-FLS formaldehyde recycling system：The 100 μL reaction system contained 50 mM potassium phosphate buffer (pH 7.5), 5 mM MgSO_4_, 2 mg mL^-1^ PK, 2 mg mL^-1^ GALS or FLS, 10 mM DHA, and 1 mM ThDP. The reactions conducted at 37 ℃ for 2 hours. 100 μL sulfuric acid (5%) was added to the samples to terminate the reactions. Acetic acid was then detected by HPLC. HPLC conditions: column, Aminex HPX-87H (Bio-Rad); detection wavelength, 210 nm; mobile phase, 5 mM sulphuric acid; flow rate, 0.6 mL min^-1^; sample volume, 20 µL; column temperature, 40 ℃.

Acetic acid and DHA detection method is the same as above. Formaldehyde was detected by chromogenic method. Chromogenic detection of formaldehyde is as follows: dilute the sample to the appropriate concentration (0.1-1 mM), add 80 μL chromogenic solution to 120 µL of the diluted sample. The reaction conducted at 60 ℃ for 10 mins, and centrifugal at 12000×g for 5 mins. Take 150 μL to measure the absorbance at 414 nm. Calculate the concentration of formaldehyde in the sample according to the standard curve. Chromogenic solution is prepared as follows: Dissolve 250 g of ammonium acetate in 900 mL of ddH_2_O, then add 30 mL of acetic acid and 2.5 mL of acetylacetone, adjust the pH of the solution to 6 with acetic acid, and finally add ddH_2_O to a volume of 1L.

**Process analysis of the MAG pathway for F6P**

The 1 mL reaction system contained 50 mM potassium phosphate buffer (pH 7.5) 5 mM MgSO_4_, 2 mg mL^-1^ PK, 0.5 mg mL^-1^ EcHAD, 1 mg mL^-1^ Ps-LRHI, 1 mM ThDP, 10 mM F6P. The reactions conducted at 37 ℃ for 10 hours. Samples were taken out every two hours for analysis. The reactions were terminated by heating at 95 ℃ for 5 minutes, and then cooled down to 37 ℃. Added the 5U alkaline phosphatase, and conducted at 37 ℃ for 4 hours, equal volume acetonitrile was used to terminate the reaction. Fructose, D-ETS, D-EUS, GALD, and acetic acid were detected by HPLC.

Sample derivatization: Reaction samples (50 μL) were mixed with a solution of O-benzylhydroxylamine hydrochloride (50 μL, 0.14 mmol mL^-1^) (pyridine: methanol: water = 33:15:2). After incubation at 50 ℃ for 60 mins, samples were diluted in methanol (100 µL) and directly analyzed by HPLC. HPLC conditions: column, X-Bridge TM C18, 5 µm, 4.6×250 mm column from Waters (Milford, USA); sample volume, 30 µL; mobile phase, solvent system (A) aqueous trifluoroacetic acid (TFA) (0.1 % (v/v) and (B) TFA (0.095% (v/v)) in CH_3_CN/H_2_O (4:1), gradient elution from 20-60 % B in 16 mins; flow rate, 1 mL min^-1^; detection wavelength, 215 nm; column temperature, 35 ℃.

**Process analysis of the MAG pathway for Xu5P**

The 1 mL reaction system contained 50 mM potassium phosphate buffer (pH 7.5) 5 mM MgSO_4_, 2 mg mL^-1^ PK, 1 mg mL^-1^ CpHAD, 0.1 mg mL^-1^ TIM, 2 mg mL^-1^ FLS, 1 mM ThDP, and 10 mM Xu5P. The reactions conducted at 37 ℃ for 10 hours. And samples were taken out every two hours for analysis. The reactions were terminated by heating at 95 ℃ for 5 minutes then cooled down to 37 ℃. Added the 5U alkaline phosphatase, and conducted at 37 ℃ for 4 hours, equal volume acetonitrile was used to terminate the reaction. D-xylulose, GCD, DHA, and acetic acid were detected by HPLC. Formaldehyde was detected by chromogenic method as before.

HPLC detection for D-xylulose, GCD and DHA. Sample derivatization: Reaction samples (50 µL) were mixed with a solution of O-benzylhydroxylamine hydrochloride (50 µL, 0.14 mmol mL^-1^) (pyridine: methanol: water = 33:15:2). After incubation at 50 ℃ for 60 mins, samples were diluted in methanol (100 μL) and directly analyzed by HPLC chromatography. HPLC conditions: column, X-BridgeTM C18, 5 μm, 4.6×250 mm column from Waters (Milford, USA); sample volume, 30 μL; mobile phase, solvent system (A) aqueous trifluoroacetic acid (TFA) (0.1 % (v/v) and (B) TFA (0.095% (v/v)) in CH_3_CN/H_2_O (4:1), gradient elution from 20-60 % B in 16 mins; flow rate, 1 mL min^-1^; detection wavelength, 215 nm; column temperature, 35 ℃. HPLC condition of acetic acid were the same as before.

**MAG pathway for the utilization of C1-C4 carbon sources**

The MAG pathway for D-EUS and GALD. The 200 µL reaction system contained 50 mM potassium phosphate buffer (pH 7.5), 5 mM MgSO_4_, 2 mg mL^-1^ PK, 1 mM ThDP, 10 mM GALD or 5 mM D-EUS. The reactions conducted at 37 ℃ overnight.

The MAG pathway for FALD and DHA. The 200 µL reaction system contained 50 mM potassium phosphate buffer (pH 7.5), 5 mM MgSO_4_, 2 mg mL^-1^ PK, 2 mg mL^-1^ FLS, 1 mM ThDP, 20 mM FALD or 10 mM DHA. The reactions conducted at 37 ℃ overnight.

HPLC detection for AcP: add equal volume of 5% sulfuric acid to the sample for completely decomposing AcP into acetic acid. Acetic acid was then detected by HPLC. HPLC conditions: column, Aminex HPX-87H (Bio-Rad); detection wavelength, 210 nm; mobile phase, 5 mM sulphuric acid; flow rate, 0.6 mL min^-1^; sample volume, 20 µL; column temperature, 40 ℃.

**The production of acetic acid using glucose *in vivo***

The *E.coli* MG1655(DE3)(△*ldhA*, △*frdBC*, △*pflB*, △*adhE*, △*zwf*, △ *epd*, △*fucO*, △*yqhD*, △*aldA*, △*ptaI* and △*acs*) strain containing bi-plasmid (pET-28a-*yidA* and pACYC-Duet-*pk*-*l-rhi*) was used. The cells that had induced the protein in 2YT were re-inoculated into reaction system, containing 50 mM potassium phosphate, 5 mM MgSO_4_, and 10 mM glucose. Cells from different culture times were collected and used to detect metabolites. Organic acid was monitored by HPLC. Glucose was monitored by biochemical analysis instrument. HPLC condition of acetic acid was the same as before.

**Fig. S1.**

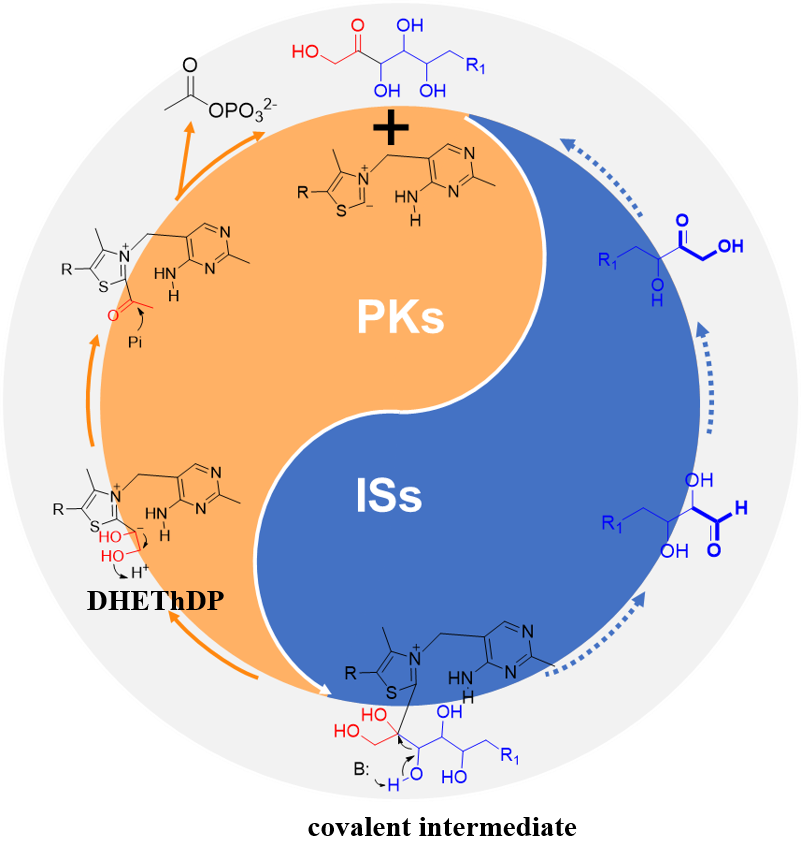

**Supplementary Figure 1. The design of the Minimal Artificial Glycolytic (MAG) pathway.** The orange part indicates the forming process of AcP from ketoses by phosphoketolase (PK). The blue part indicates the conversion of aldose molecules to ketose molecules by isomerases (ISs).

**Fig. S2**

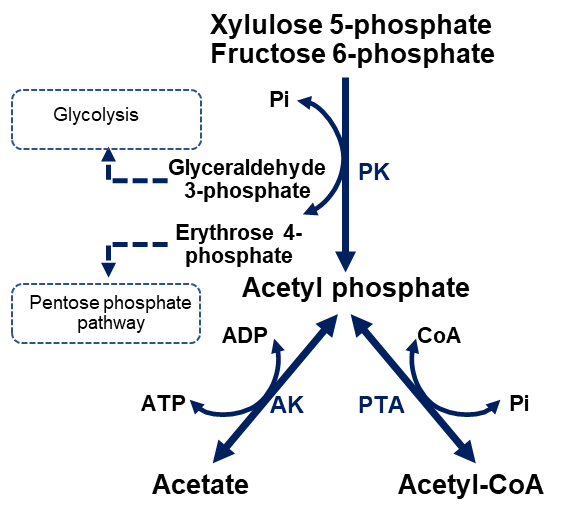

**Supplementary Figure 2.** **The phosphoketolase (PK) pathway in nature.** Fructose-6-phosphate (F6P) and xylulose-5-phosphate (Xu5P) are converted into and erythorse-4-phosphate / glyceraldehyde-3-phosphate (E4P/G3P) and acetyl-phosphate (AcP), which not only can be converted into acetyl-CoA by phosphate acetyltransferase (PTA), but also can be converted to ATP and acetate by acetate kinase (AK).

**Fig. S3**

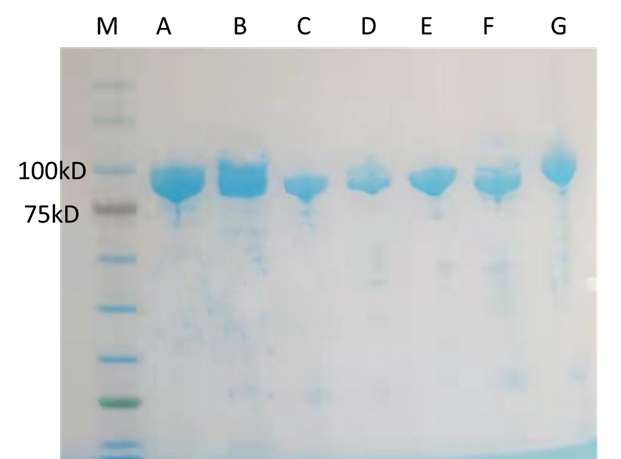

**Supplementary Figure 3. SDS-PAGE analysis of PK from different species.** A, PK1; B, PK2; C, PK3; D, PK4; E, PK5; F, PK6; G, PK7. The detailed information of PK sees Table S7 and Table S8.

**Fig. S4.**

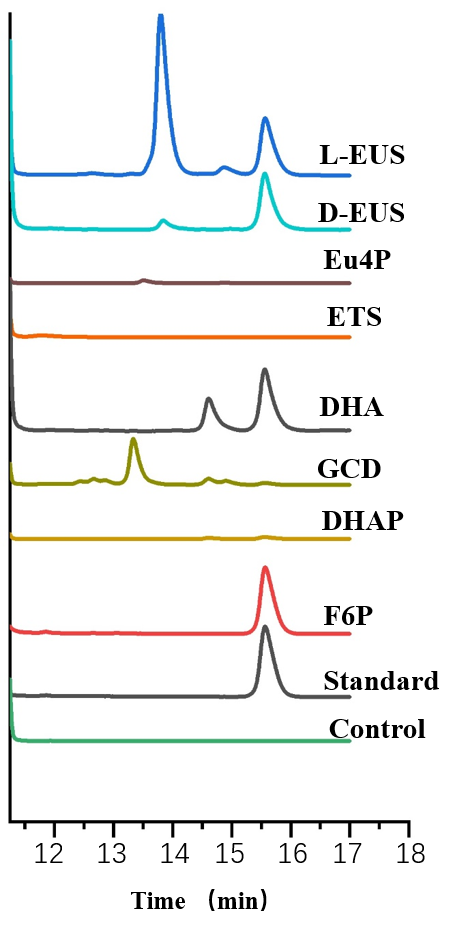

**Supplementary Figure 4.** **The activity test of PK from *Bifidobacterium* (BbPK) on different sugars.** The product acetyl phosphate was converted to acetic acid, which was detected by HPLC. The reaction system without PK was used as the control. L-EUS, L-erythrulose; D-EUS, D-erythrulose; Eu4P, D-erythrulose-4-phosphate; ETS, D-erythrose; DHA, dihydroxyacetone; GCD, D-glyceraldehyde; DHAP, dihydroxyacetone phosphate; F6P, D-fructose-6-phosphate**.**

**Fig. S5.**

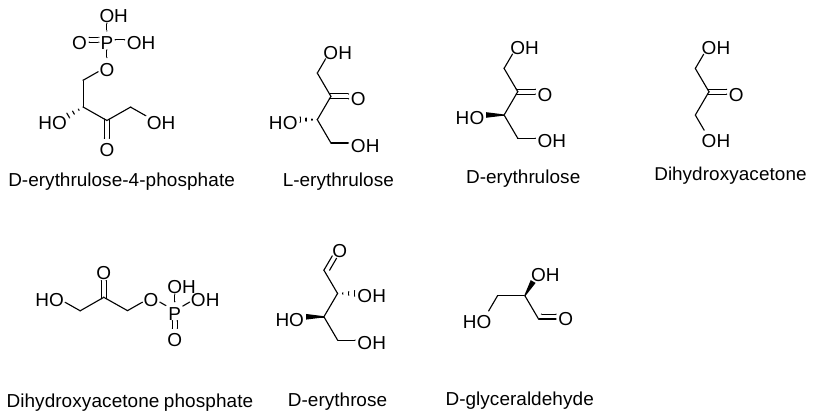

**Supplementary Figure 5**. **The** **structure of tested substrates.**

**Fig. S6.**

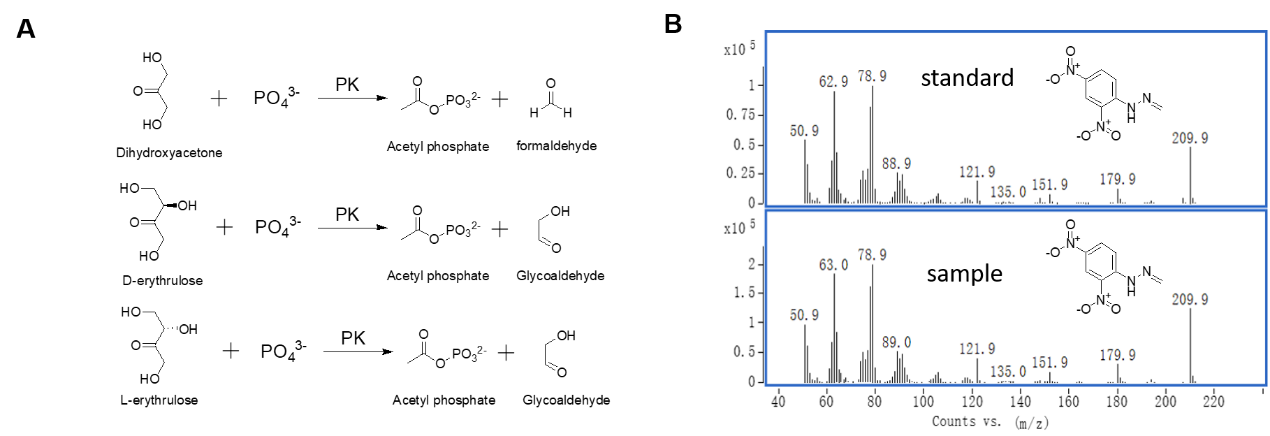

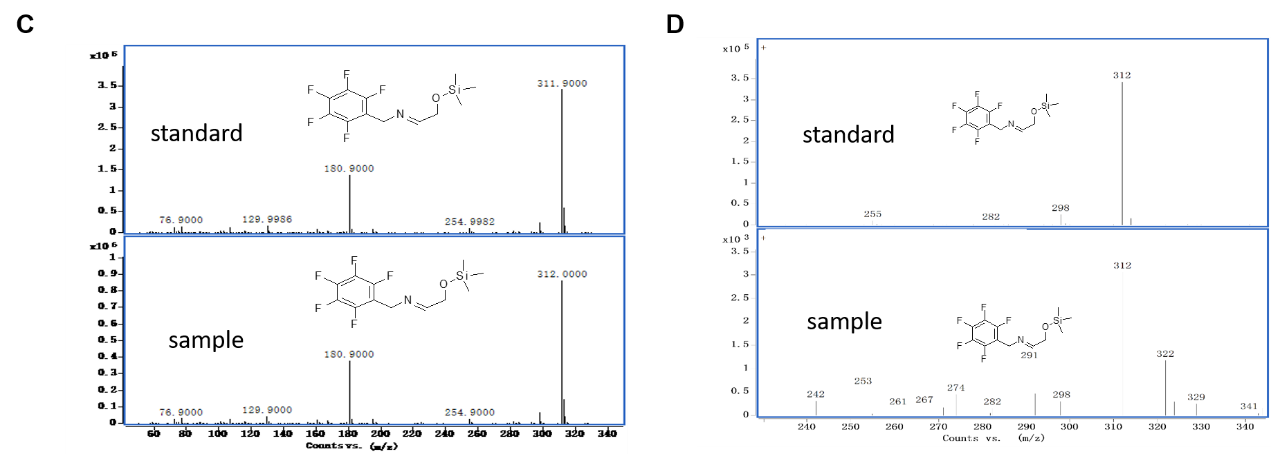

**Supplementary Figure 6.** **Detection of formaldehyde and glycoaldehyde in PK-** **catalyzed reaction system. A** Schematic illustration of the reaction of dihydroxyacetone, D-erythrulose, and L-erythrulose catalyzed by PK. **B** Formaldehyde was detected by GC-MS in PK-catalyzed dihydroxyacetone reaction system. **C** Glycoaldehyde was detected by GC-MS in PK-catalyzed D-erythrulose reaction system. **D** Glycoaldehyde was detected by GC-MS in PK-catalyzed L-erythrulose reaction system. Sample derivatization methods see materials and methods.

**Fig. S7.**

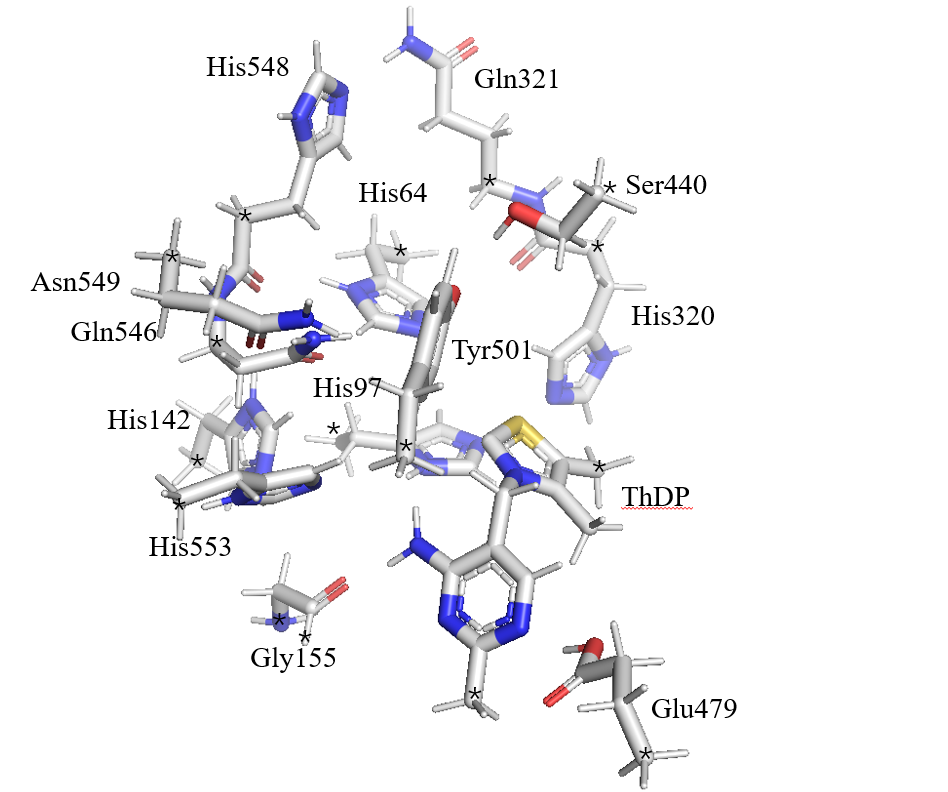

**Supplementary Figure 7.** **Computational model for calculation**. The model contains 218 atoms with a total charge of +1, including the side chains of His64, His553, Glu479, Tyr501, Gln321, Ser440, His142, Gly155, His320, Asn549, Gln546, His548, His97, and the cofactor ThDP. The fixed atoms are labeled by asterisks.

**Fig. S8.**

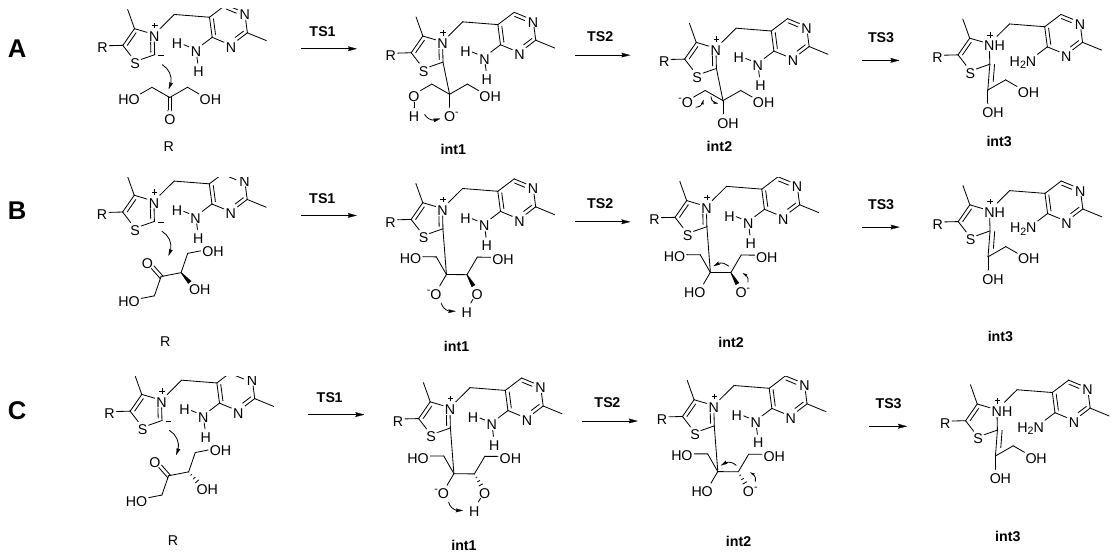

**Supplementary Figure 8.** **Proposed catalytic mechanism of PK**. **A** The forming process of 2-α, β-dihydroxyethylidene-ThDP (DHEThDP) from 1,3-dihydroxyacetone. **B** The forming process of DHEThDP from D-erythrulose. **C** The forming process of DHEThDP from L-erythrulose. Upon binding of the substrate to ThDP, the first step is a C−C bond formation that leads to an alkoxide tetrahedral intermediate. Next, an intramolecular proton transfer takes place from the hydroxyl group in C3 to the alkoxide. The last step is a C−C bond cleavage to form the DHEThDP. R, reactant; int1, intermediate 1; TS1, transition state 1.

**Fig. S9.**

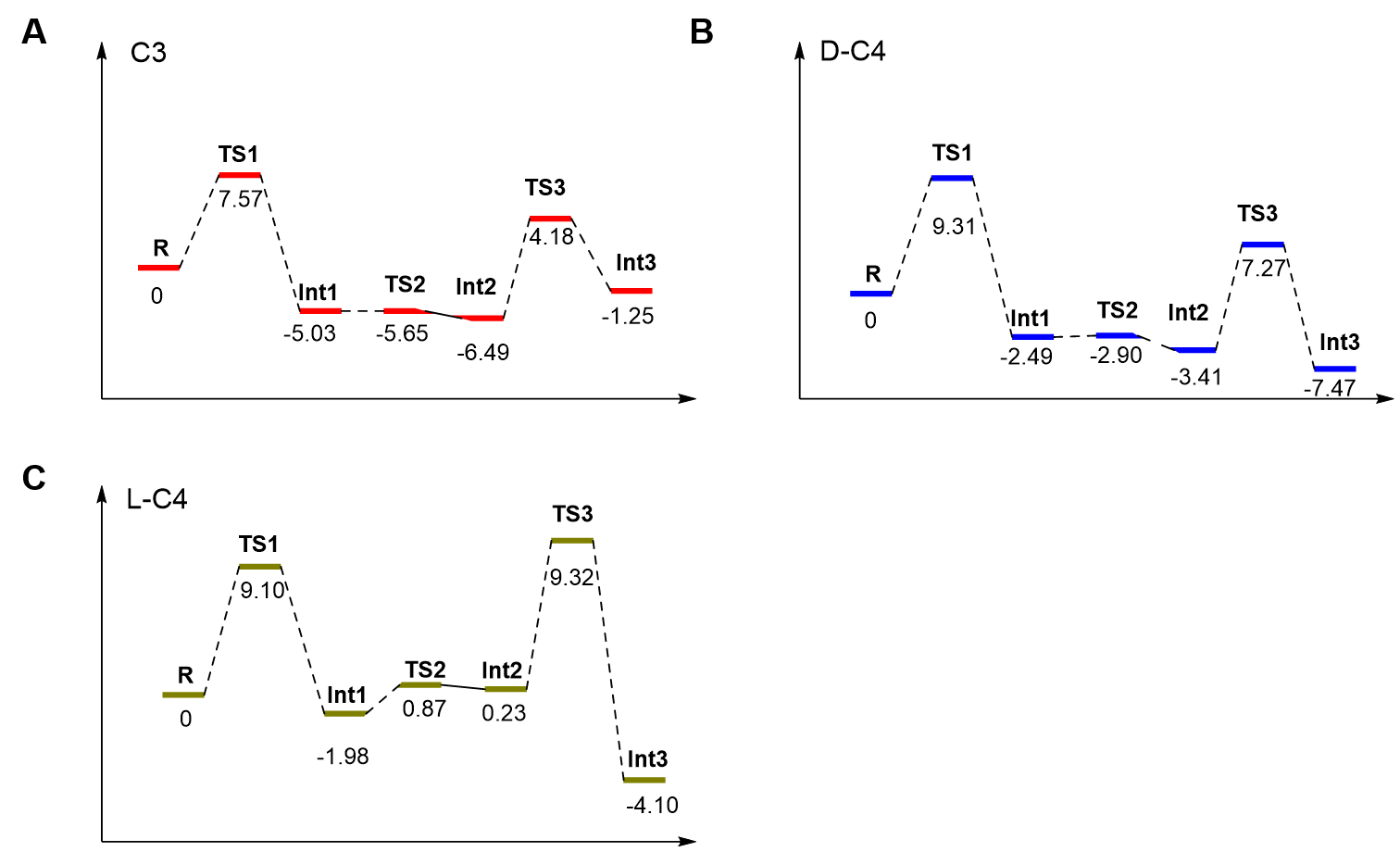

**Supplementary Figure 9. Calculated energy profiles for the forming process of 2-α, β-dihydroxyethylidene-ThDP (DHEThDP) from short chain ketoses.** **A** The energy profiles for 1,3-dihydroxyacetone. **B** The energy profiles for D-erythrulose. **C** The energy profiles for L-erythrulose. Energies are given in kilocalories per mole. Note: After adding the large basis set, solvation, and zero-point energy corrections, the energies of TS2 for 1, 3-dihydroxyacetone and D-erythrulose were calculated to be lower than those of int1. Therefore, intramolecular proton transfer of 1,3-dihydroxyacetone and D-erythrulose can be assumed to be barrierless or to occur with very low barriers.

**Fig. S10.**

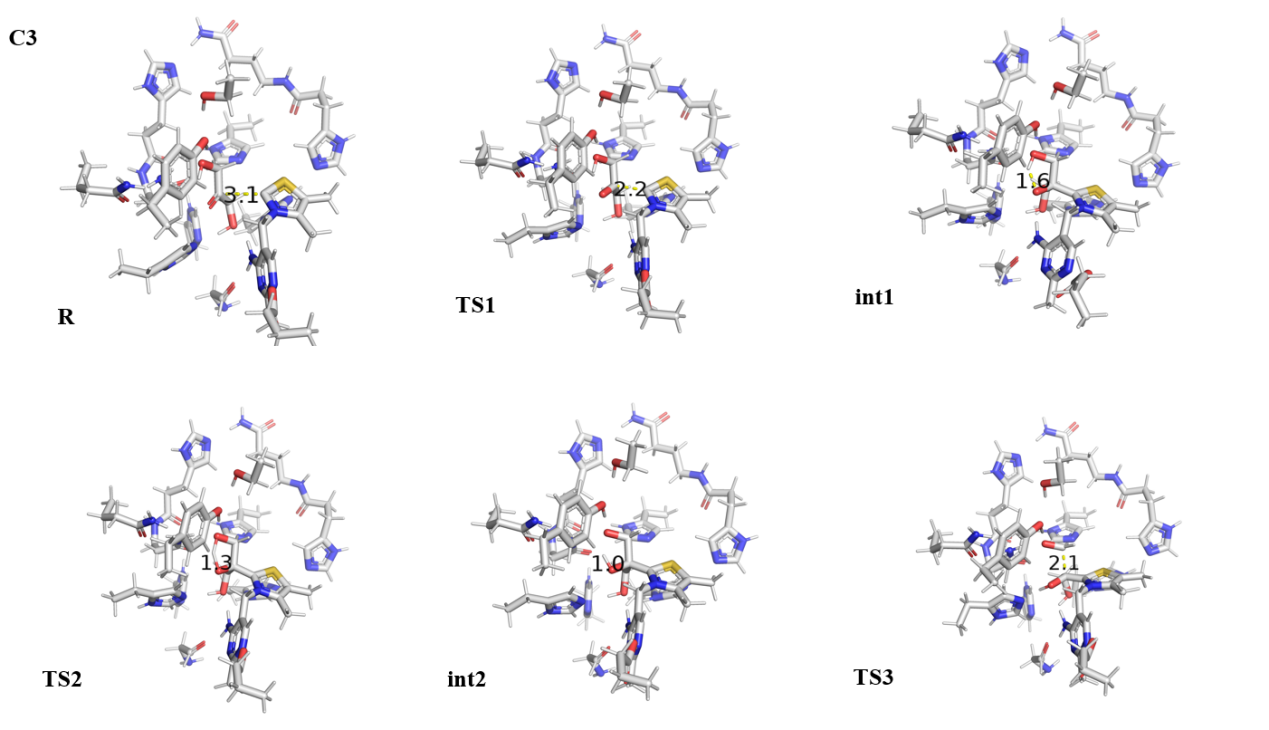

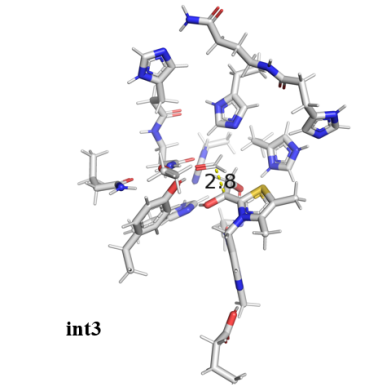

**Supplementary Figure 10. Optimized structures of the transition states and intermediate involved in the forming process of DHEThDP from 1,3-dihydroxyacetone.** The key bond distances change is shown in the figure. Selected distances are given in Å. The distances between C2 of ThDP and carbonyl C of substrate changes from 3.1 Å in Reactant (R) to 2.2 Å in transition state 1 (TS1). The distance between O and H changes from 1.6 Å in intermediate 1 (int1) to 1.3 Å in TS2. The distance of C2 and C3 of substrate changes from 2.1 Å in TS3 to 2.8 Å in int3.

**Fig. S11**

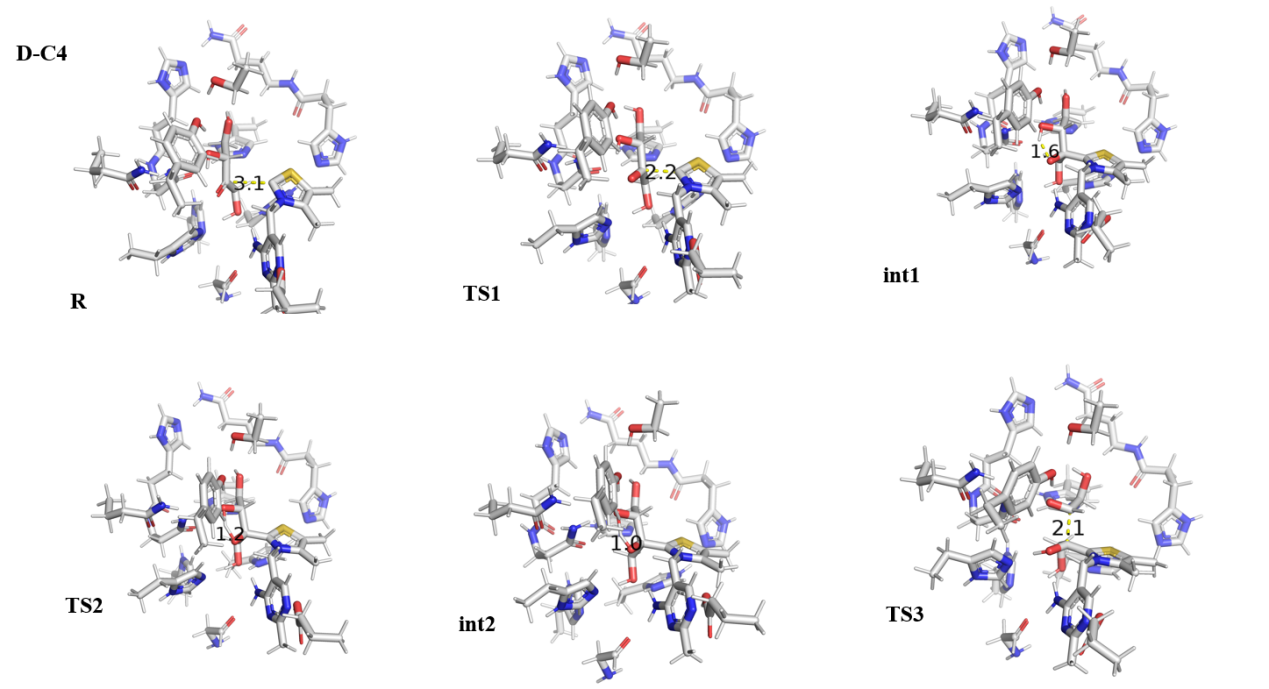

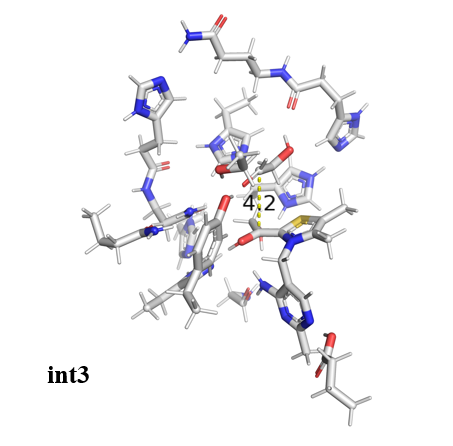

**Supplementary Figure 11. Optimized structures of the transition states and intermediate involved in the forming process of DHEThDP from D-erythrulose.** The key bond distances change is shown in the figure. Selected distances are given in Å. The distances between C2 of ThDP and carbonyl C of substrate changes from 3.1 Å in Reactant (R) to 2.2 Å in transition state 1 (TS1). The distance between O and H changes from 1.6 Å in intermediate 1 (int1) to 1.2 Å in TS2. The distance of C2 and C3 of substrate changes from 2.1 Å in TS3 to 4.2 Å in int3.

**Fig. S12**

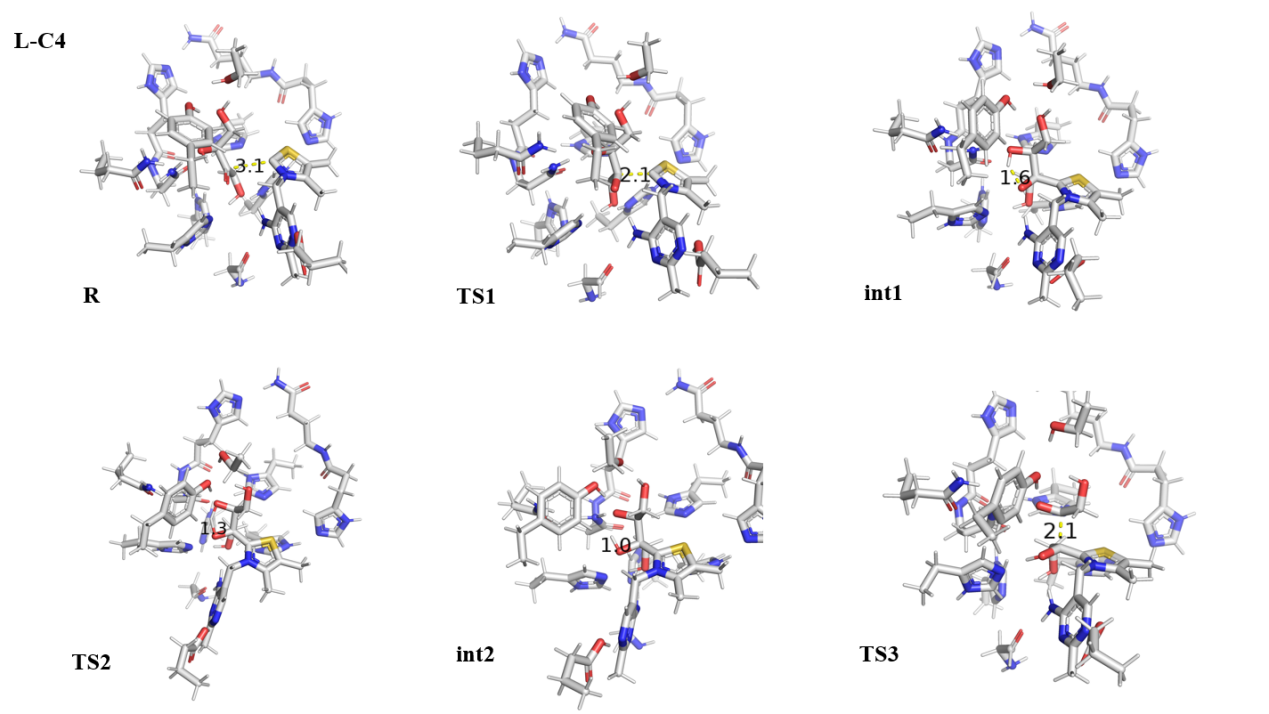

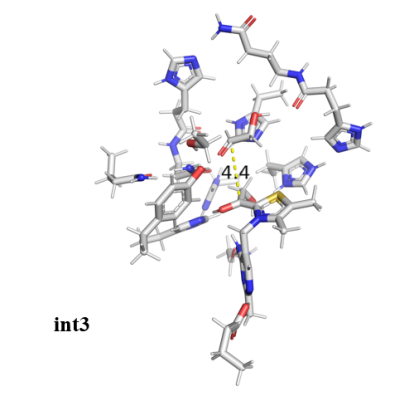

**Supplementary Figure 12. Optimized structures of the transition states and intermediate involved in** **the forming process of DHEThDP from L-erythrulose.** The key bond distances change is shown in the figure. Selected distances are given in Å. The distances between C2 of ThDP and carbonyl C of substrate changes from 3.1 Å in Reactant (R) to 2.1 Å in transition state 1 (TS1). The distance between O and H changes from 1.6 Å in intermediate 1 (int1) to 1.3 Å in TS2. The distance of C2 and C3 of substrate changes from 2.1 Å in TS3 to 4.4 Å in int3.

**Fig. S13**

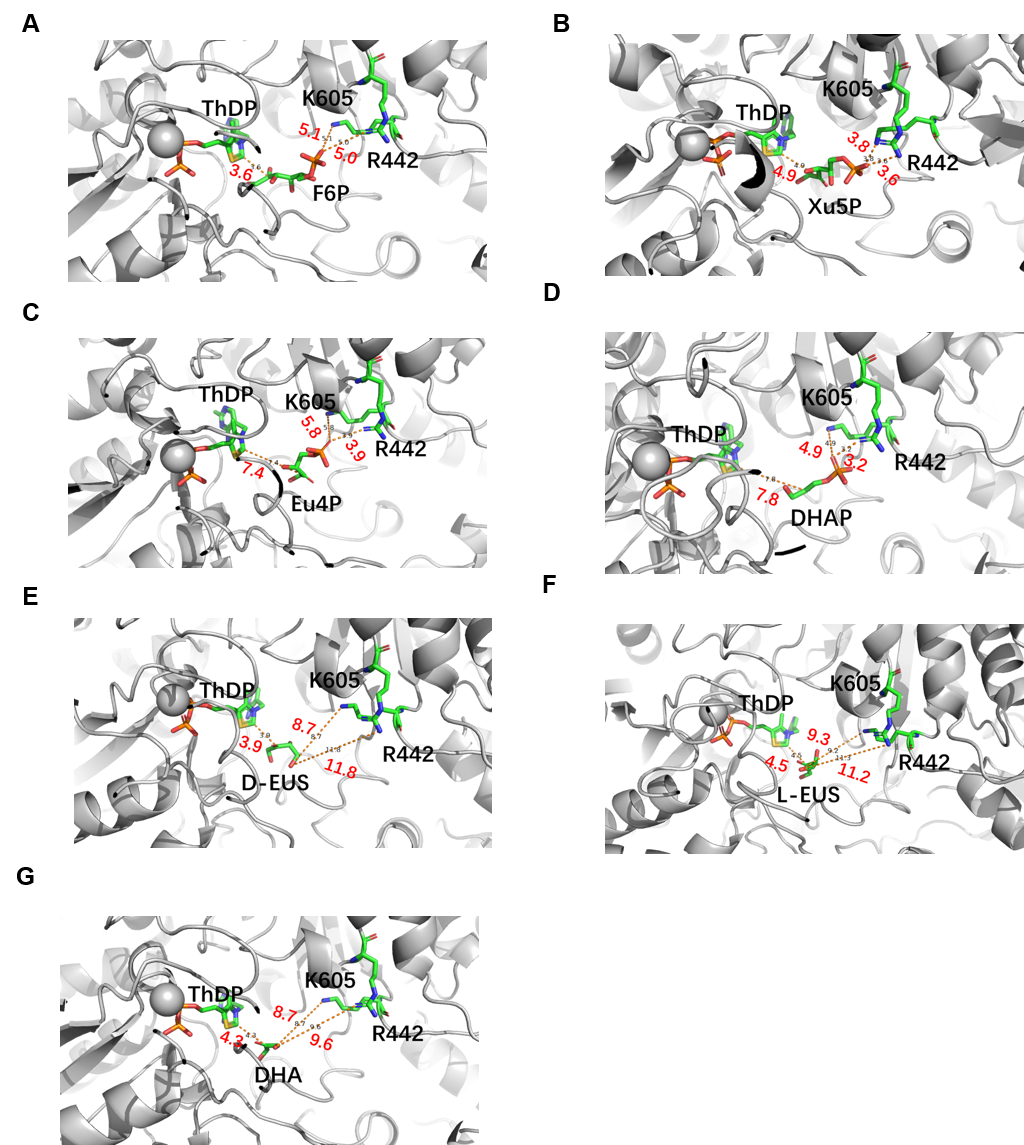

**Supplementary Figure 13. The docking results of different substrates in PK.** The PK was shown in cartoon and colored grey. The ThDP, ligands, and two key residues R442 and K605 were shown in stick. The C, N, O, P, and S atoms were colored green, blue, red, orange, and yellow, respectively. The distance between the substrates and ThDP and the distances between the phosphate moiety of substrates and the key basic residues were shown as dashed lines. Selected distances are given in Å.

**Fig. S14**

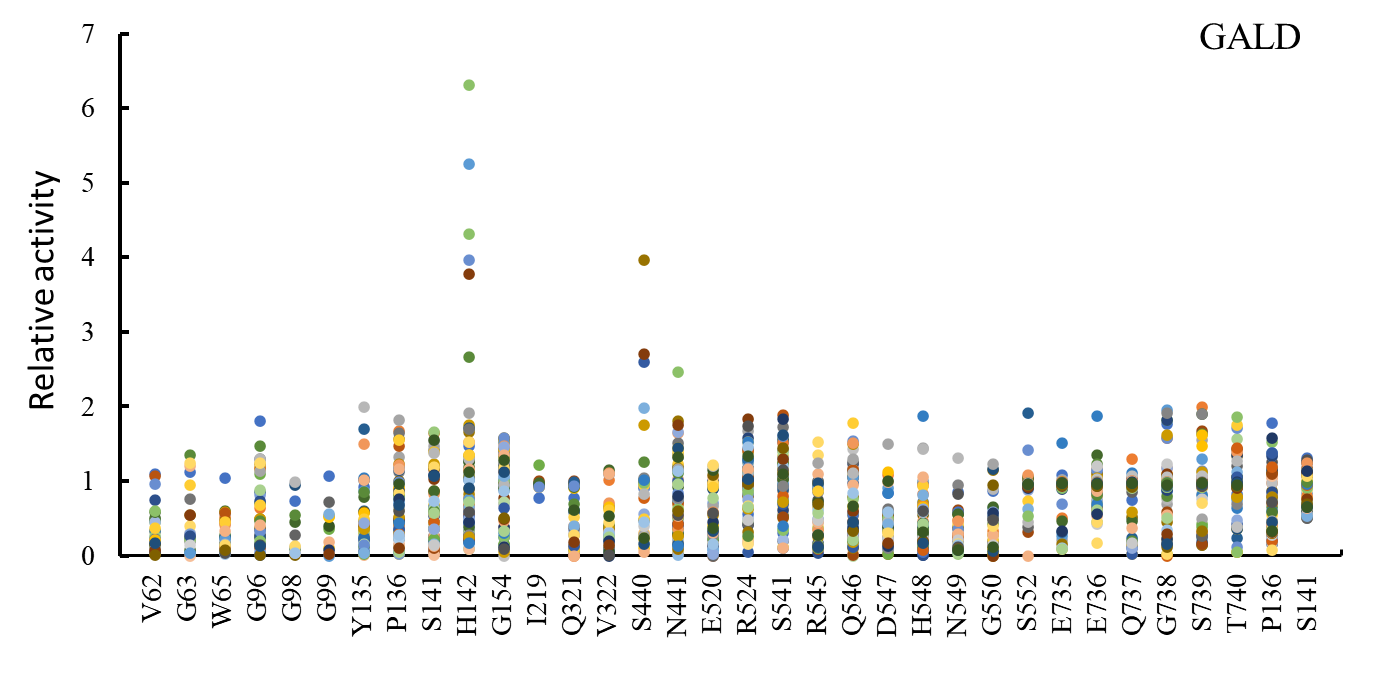

**Supplementary Figure 14. High-throughput screening of glycolaldehyde (GALD).** The x-axis labels represent the selected location in the BbPK. The y-axis labels represent the relative catalytic activities of the different mutants. Relative activity was defined as the ratio of the reduction of substrate for mutants to that of the wild type.

**Fig. S15**

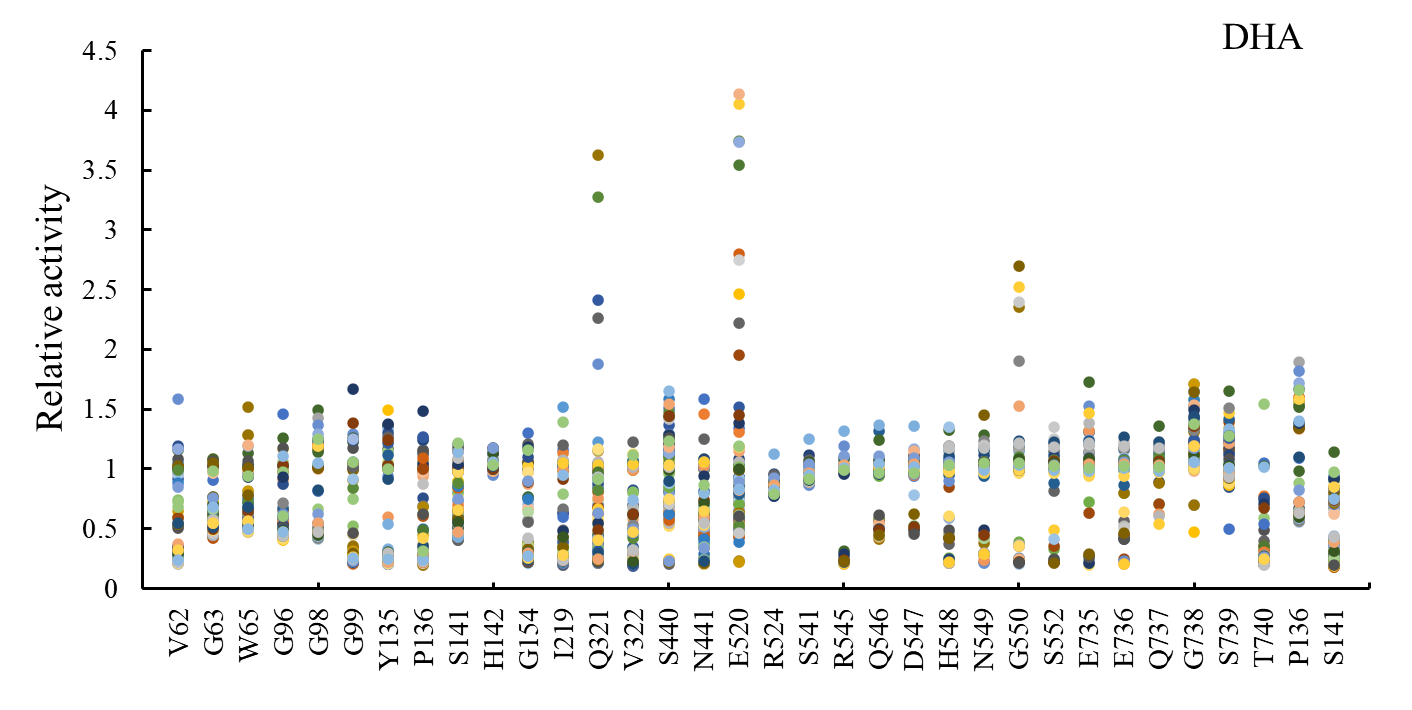

**Supplementary Figure 15. High-throughput screening of dihydroxyacetone (DHA).** The x-axis labels represent the selected location in the BbPK. The y-axis labels represent the relative catalytic activities of the different mutants. Relative activity was defined as the ratio of the titer of formaldehyde for the mutants to that of the wild type.

**Fig. S16**

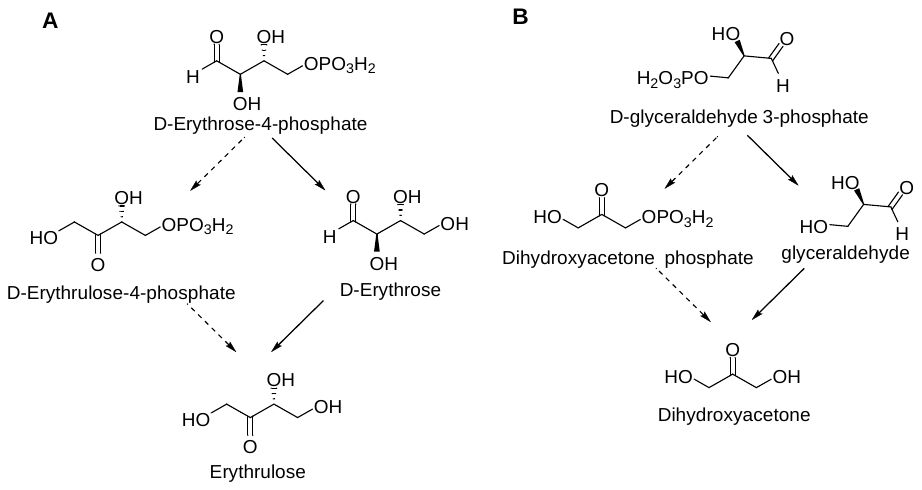

**Supplementary Figure 16. Two different pathways for converting E4P/G3P to EUS/DHA. A** The two conversion pathways of D-erythrulose (D-EUS) from D-erythrose-4-phosphate (E4P). **B** The two conversion pathways of dihydroxyacetone (DHA) from D-glyceraldehyde-3-phosphate (G3P). Dashed arrows represent the pathway that is first isomerized and then dephosphorylated. Solid arrows represent the pathway that is first dephosphorylated and then isomerized.

**Fig. S17**

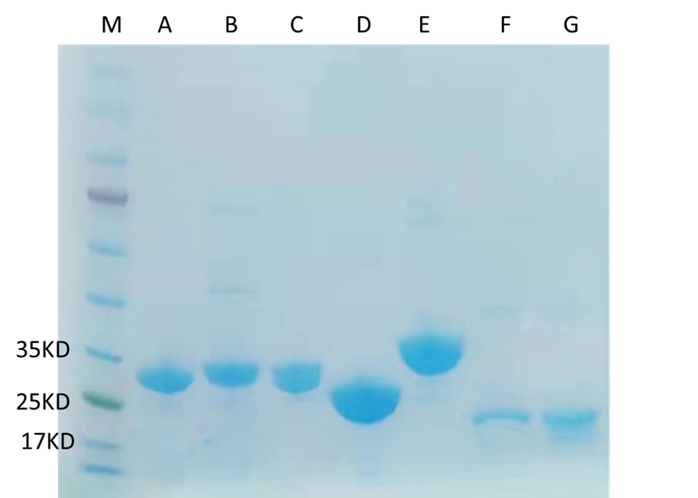

Supplementary Figure 17. SDS-PAGE analysis of phosphatases from different species. A, pfHAD; B, EcHAD; C, NbIMP; D, TmHAD; E, CpHAD; F, XiHAD; G, CgHAD. The detailed information of phosphatases sees Table S7 and Table S8.

**Fig. S18**

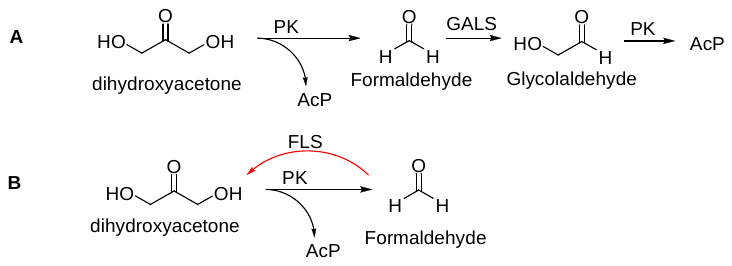

**Supplementary Figure 18. Two different pathways to recycle formaldehyde. A** Formaldehyde is converted to glycolaldehyde by glycolaldehyde synthase (GALS). Glycolaldehyde is then converted to AcP by PK. **B** Formaldehyde is converted to dihydroxyacetone by formolase (FLS). Dihydroxyacetone is then converted to AcP by PK.

**Fig. S19**

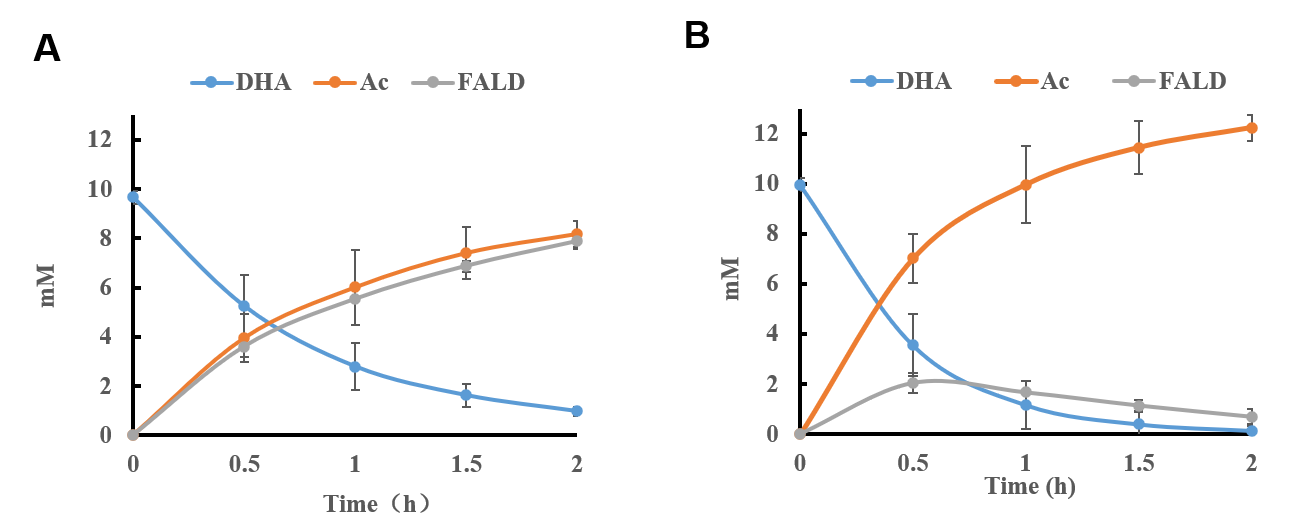

**Supplementary Figure 19. Recycle the formaldehyde generated in the MAG pathway. A** The conversion of DHA to AcP using only BbPK *in vitro*. **B** Formolase (FLS) was used to recycle formaldehyde. The red curve represents the change of acetic acid concentration. The blue curve represents the change of DHA concentration. The grey curve represents the change of formaldehyde concentration. Acetic acid was detected by HPLC. Error bars represent s.d. (standard deviation), n = 3.

**Fig. S20**

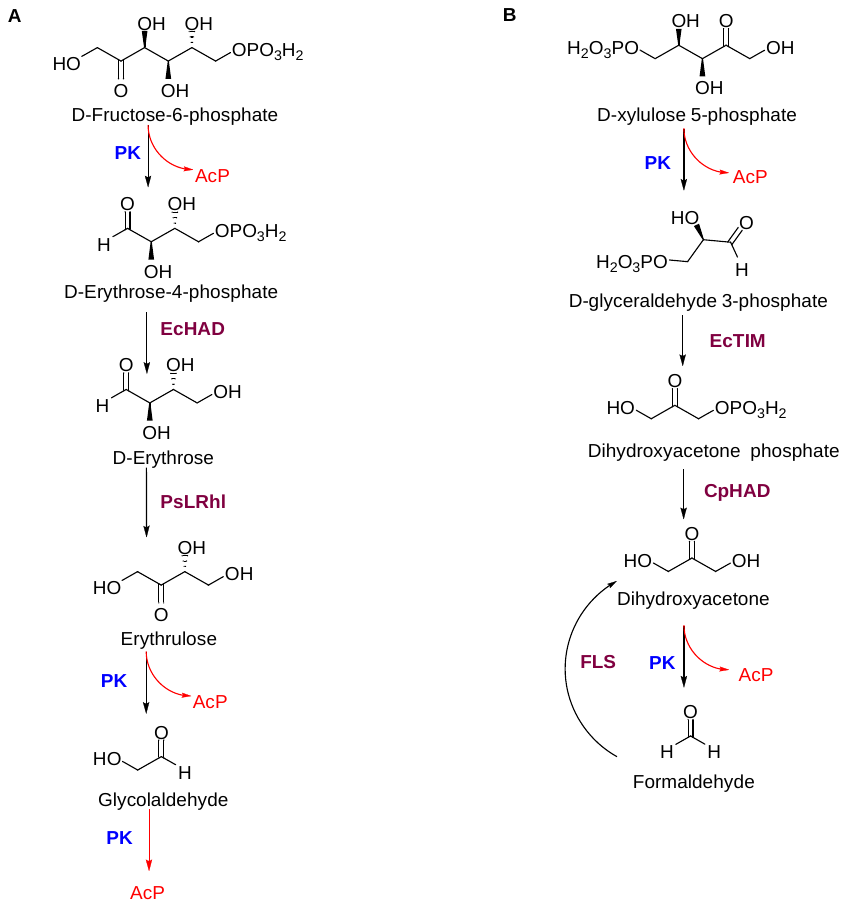

Supplementary Figure 20. The MAG pathway for F6P and Xu5P. A The MAG pathway for F6P include phosphoketolase (PK) from *Bifidobacterium*（BbPK）, HAD-like hydrolase (EcHAD) from *Escherichia coli*, and L-rhamnose isomerase (PsLRhI) from *Pseudomonas stutzeri*. B The MAG pathway for Xu5P consist of BbPK, triose phosphate isomerase (EcTIM) from *E. coli*, sugar phosphatase from *Candida parapsilosis* (CpHAD), and formolase (FLS).

**Fig. S21**

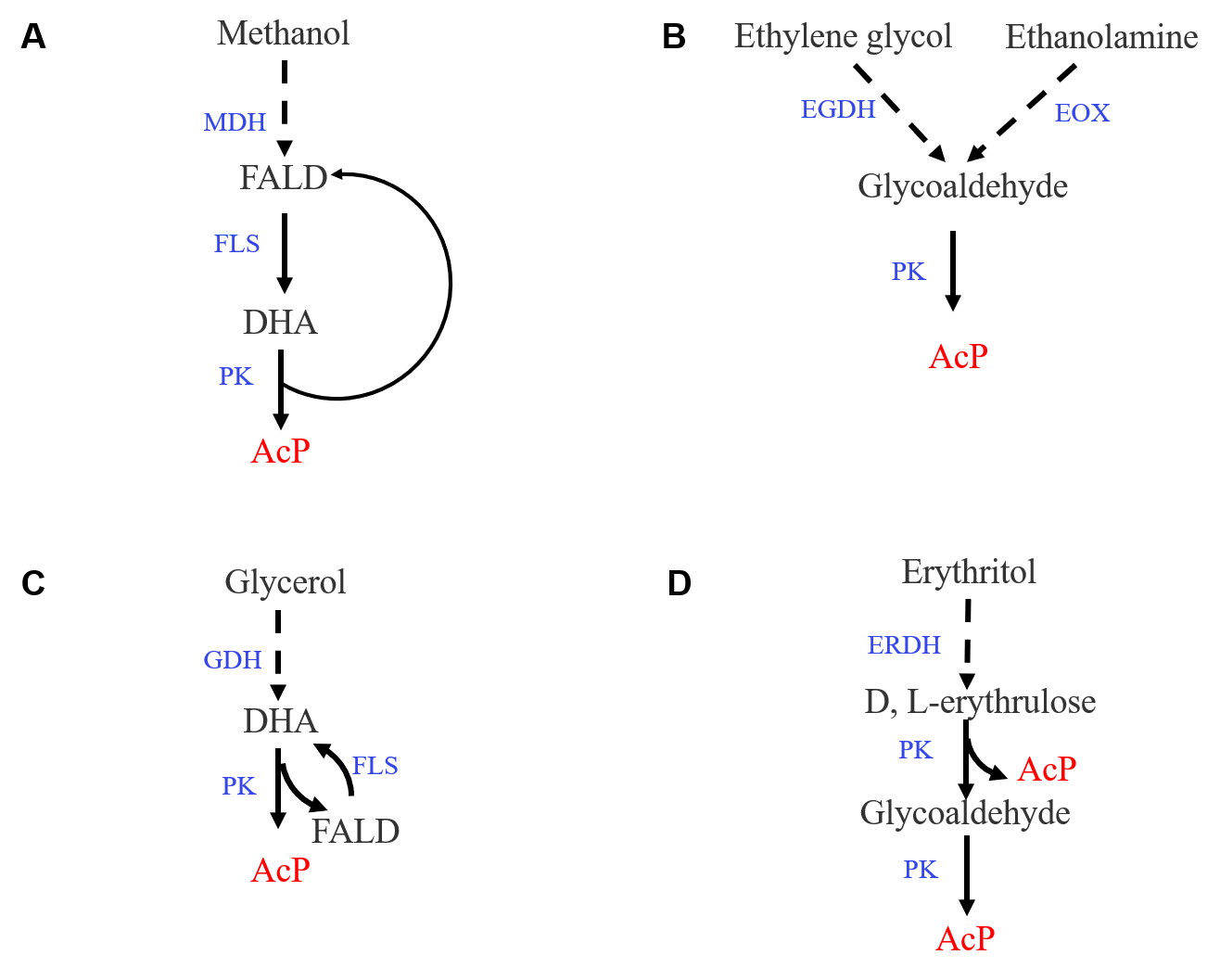

**Supplementary Figure 21. The MAG pathway for the utilization of C1, C2, C3, and C4 carbon sources.** **A** Methanol is converted to DHA and then converted to AcP via MAG pathway. MDH, methanol dehydrogenase. **B** Ethylene glycol or ethanolamine is converted to glycoaldehyde and then converted to AcP via MAG pathway. EGDH, ethylene glycol dehydrogenase; EOX, ethanolamine oxidase. **C** Glycerol is converted to DHA and then converted to AcP via MAG pathway. GDH, glycerol dehydrogenase. **D** Erythritol is converted to erythrulose and then converted to AcP via MAG pathway. ERDH, erythritol dehydrogenase. Dashed arrows represent the process that is not tested.

**Fig. S22**

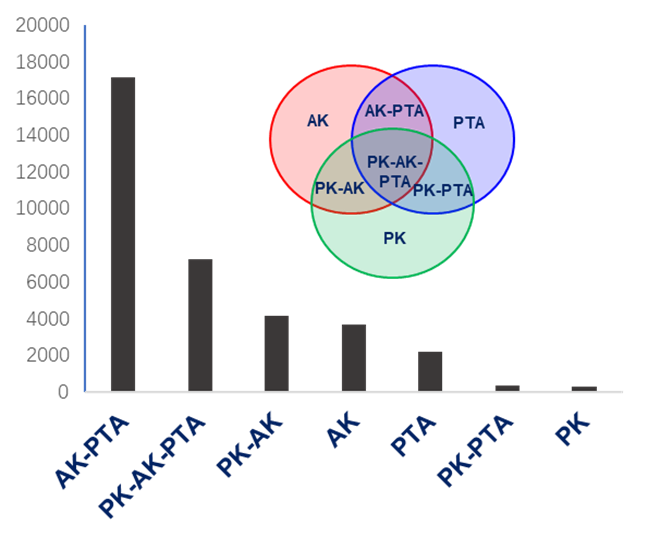

**Supplementary Figure 22. The investigation of the distribution of PK, AK and PTA genes in about 50,000 sequenced genomes.** The numbers in the bar are corresponding to the Venn diagram.

**Fig. S23**

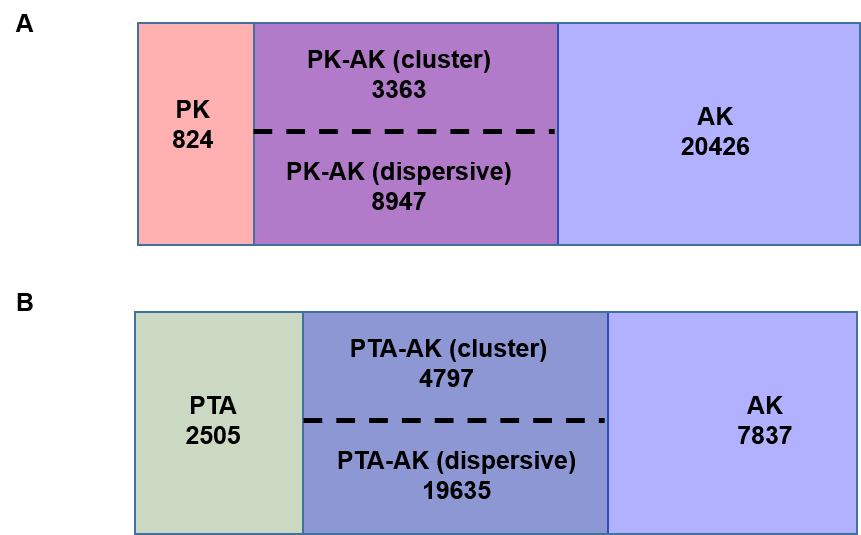

**Supplementary Figure 23. The distribution of PK and AK, and PTA and AK in sequenced genomes.** **A** venn diagram of PK or/and AK genes distributed in sequenced genomes. The ‘PK 824’ represents 824 species with PK genes, but without AK genes; The ‘PK-AK (cluster) 3363’ represents 3363 species including PK-AK gene cluster in the genomes; The ‘PK-AK (dispersive) 8947’ represents 8947 species in which PK and AK genes are dispersively distributed in the genomes; The ‘AK 20426’ represents 20426 species with AK genes, but without PK genes. **B** venn diagram of PTA or/and AK genes distributed in sequenced genomes.

**Fig. S24**

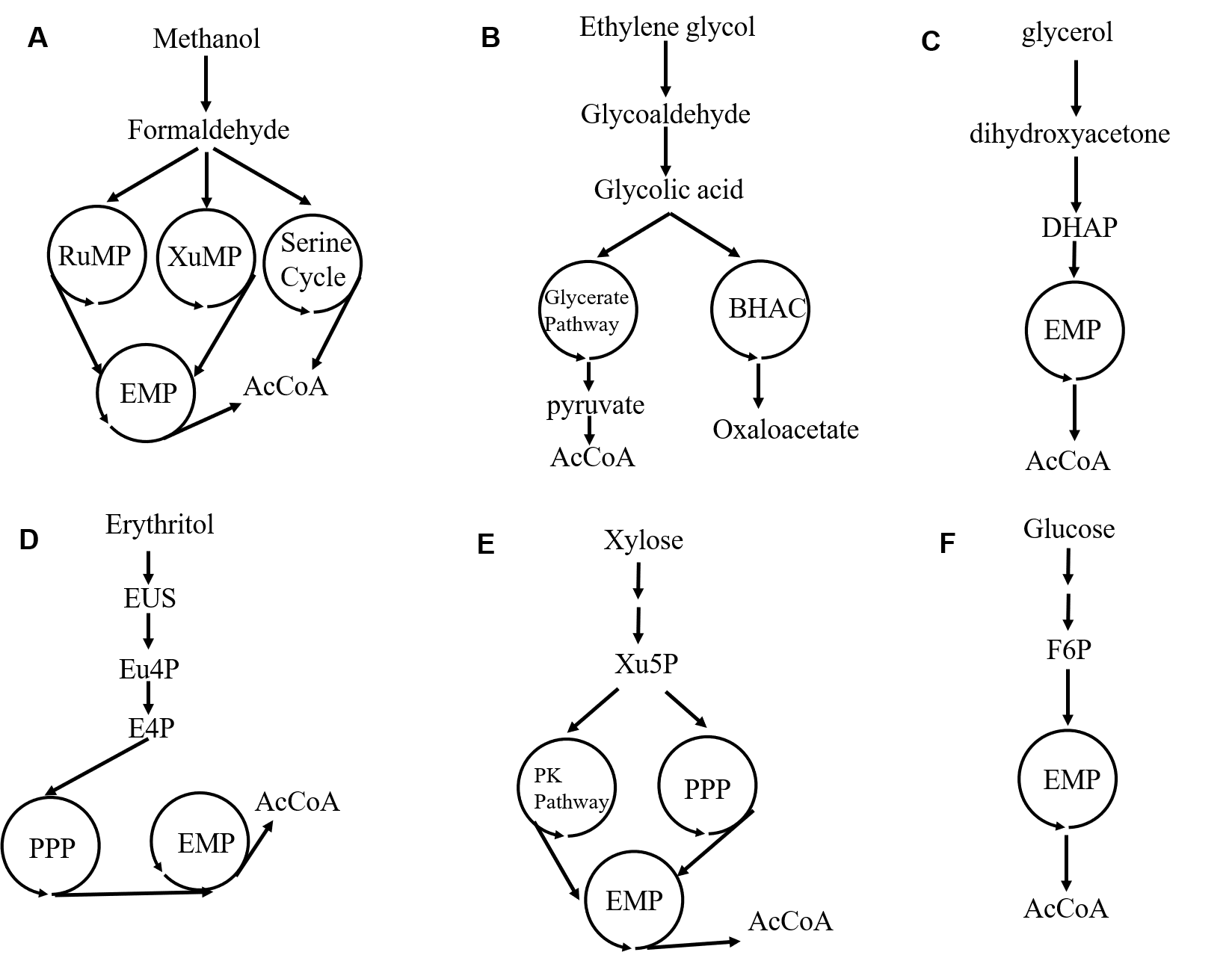

**Supplementary Figure 24. Natural metabolic pathways of C1-C6 carbon sources.** **A** Natural metabolic pathways of methanol. RuMP, ribulose monophosphate pathway; XuMP, xylulose monophosphate pathway; EMP, Embden–Meyerhoff–Parnas pathway. **B** Natural metabolic pathways of ethylene glycol. BHAC, β-hydroxyaspartate cycle. **C** Natural metabolic pathways of glycerol**. D** Natural metabolic pathways of erythritol. PPP, pentose phosphate pathway. **E** Natural metabolic pathways of D-xylose. **F** Natural metabolic pathways of D-glucose.

**Table S1. Energies and energy corrections for the forming process of DHEThDP from DHA.**

| Structure | Eel (a.u.) | Ebb (a.u.) | Esol (a.u.) | ZPE  (a.u.) | Etot (a.u.) | Erel  (kcal/mol) |
| --- | --- | --- | --- | --- | --- | --- |
| R | -5435.47252 | -5436.93303 | -5435.59435 | 1.95631 | -5435.09855 | 0 |
| TS1 | -5435.46527 | -5436.92287 | -5435.58620 | 1.95731 | -5435.08649 | 7.57 |
| Int1 | -5435.48496 | -5436.93945 | -5435.61064 | 1.95857 | -5435.10656 | -5.03 |
| TS2 | -5435.48379 | -5436.93765 | -5435.60860 | 1.95490 | -5435.10756 | -5.65 |
| Int2 | -5435.48624 | -5436.94092 | -5435.61195 | 1.95774 | -5435.10889 | -6.49 |
| TS3 | -5435.46870 | -5436.92532 | -5435.59167 | 1.95640 | -5435.09189 | 4.18 |
| Int3 | -5435.48060 | -5436.93909 | -5435.59669 | 1.95464 | -5435.10054 | -1.25 |

**Table S2. Energies and energy corrections for the forming process of DHEThDP from D-EUS.**

| Structure | Eel (a.u.) | Ebb (a.u.) | Esol (a.u.) | ZPE  (a.u.) | Etot (a.u.) | Erel  (kcal/mol) |
| --- | --- | --- | --- | --- | --- | --- |
| R | -5549. 99773 | -5551.50070 | -5550.12326 | 1.98824 | -5549.63799 | 0 |
| TS1 | -5549.99209 | -5551.48985 | -5550.11568 | 1.99029 | -5549.62315 | 9.31 |
| Int1 | -5550.01044 | -5551.50456 | -5550.13911 | 1.99127 | -5549.64196 | -2.49 |
| TS2 | -5550.00805 | -5551.50189 | -5550.13639 | 1.98762 | -5549.64261 | -2.90 |
| Int2 | - 5550.00965 | -5551.50419 | -5550.13894 | 1.99005 | -5549.64343 | -3.41 |
| TS3 | -5549.99402 | -5551.49043 | -5550.11961 | 1.98988 | -5549.62614 | 7.27 |
| Int3 | -5550.02381 | -5551.52199 | -5550.13862 | 1.98691 | -5549.64989 | -7.47 |

**Table S3. Energies and energy corrections for the forming process of DHEThDP from L-EUS.**

| Structure | Eel (a.u.) | Ebb (a.u.) | Esol (a.u.) | ZPE  (a.u.) | Etot (a.u.) | Erel  (kcal/mol) |
| --- | --- | --- | --- | --- | --- | --- |
| R | -5550.01505 | -5551.51231 | -5550.13620 | 1.98901 | -5549.64445 | 0 |
| TS1 | -5550.00565 | -5551.49968 | -5550.12637 | 1.99045 | -5549.62995 | 9.10 |
| Int1 | -5550.02358 | -5551.51576 | -5550.14764 | 1.99221 | -5549.64761 | -1.98 |
| TS2 | -5550.01195 | -5551.50404 | -5550.13907 | 1.98810 | -5549.64306 | 0.87 |
| Int2 | -5 550.01574 | -5551.50887 | -5550.14089 | 1.98994 | -5549.64408 | 0.23 |
| TS3 | -5549.99869 | -5551.49368 | -5550.12448 | 1.98988 | -5549.62959 | 9.32 |
| Int3 | -5550.02336 | -5551.52258 | -5550.13875 | 1.98698 | -5549.65099 | -4.10 |

Eel = Energy of the optimized geometry at B3LYP/6-31G (d,p) level.

Ebb = Single-point energy of the optimized geometry at B3LYP/6-311++G (2d,2p) level.

Esol = Single-point energy of the optimized geometry at B3LYP/6-31G (d,p) and SMD solvation method with ε = 4.

Etot = Ebb + (Esolv – Eel) + ZPE

Erel = Energy relative to the lowest-energy binding mode.

ZPE =Zero-point correction

**Table S4. Enzyme kinetic parameters**

| Variants | Substrate | *k*_cat_ (s^-1^) | *K*_m_ (mM) | *k*_cat_/*K*_m_  (s^-1^· M ^-1^) | Source |
| --- | --- | --- | --- | --- | --- |
| FLS | FALD | 0.15 | 34.14 | 4.4 | ^13^ |
| GALS | FALD | 1.58 | 170 | 9.3 | ^14^ |
| BbPK | GALD | 0.16 | 51 | 3.1 | ^14^ |
| PK-H142N | GALD | 0.25 | 16.55 | 15.1 | This study |
| BbPK | DHA | 0.03 | 7.80 | 3.8 | This study |
| PK-E520I | DHA | 0.19 | 21.9 | 8.7 | This study |
| PK-Q321A | DHA | 0.07 | 6.49 | 10.8 | This study |
| BbPK | D-EUS | 0.049 | 45 | 1.1 | This study |
| PK-H142N | D-EUS | 0.035 | 8.4 | 4.2 | This study |
| BbPK | F6P | 520 | 10 | 52000 | ^15^ |
| BbPK | Xu5P | 2700 | 45 | 60000 | ^15^ |

**Table S5. Overall reactions and properties of various carbon metabolism**

| Pathway | Product for glucose | Net ATP  for glucose  and DHA | Net redox | Types of metabolic carbon sources | Enzymes | Kinetic trap | Carbon yield  % |
| --- | --- | --- | --- | --- | --- | --- | --- |
| MAG | 3 acetate | 1 and 1.5 | No | C1 C2, C3, C4, C5, C6, | 6 | No | 100 |
| EMP | 2 lactate | 1 and 1 | No | C3, C6 | 11 | No | 67 |
| ED | 2 lactate | 1 and 1 | No | C3, C6 | 11 | No | 67 |
| PPP | 6 CO_2_ | 0 | 6 | C3, C5, C6 | 11 | No | 0 |
| GLYP | - | - | - | C2, C3 | - | No | <67 |
| BHAC | - | - | - | C2 | - | No | <67 |
| RuMP | - | - | - | C1 | - | No | <67 |
| XuMP | - | - | - | C1 | - | No | <67 |
| WL | - | - | - | C1 | - | No | 100 |
| PKP | lactate + ethanol + CO_2_ | 1 | No | C6, C5, C3 | 15 | No | - |
| B-S | 1.5 acetate + lactate | 2.5 | No | C6, C5, C3 | 14 | No | - |
| NOG^16^ | 3 acetate | 2 | No | C6, C5, C3 | 11 | Yes | 100 |
| GATHCYC | 3 acetate | 2 | No | C6 | 8 | No | 100 |

Note: ED, Entner–Doudoroff pathway; GLYP, glycerate pathway; BHAC, β-hydroxyaspartate cycle; WL, Wood−Ljungdahl pathway; PKP, phosphoketolase pathway; B-S, bifid shunt; GATHCYC, Glycolysis AlTernative High Carbon Yield Cycle.

**Table S6. Primers for gene knockout**

| Primers | Sequence (5’→3’) |
| --- | --- |
| Zwf-1-Forward | 5’- gccggaacaccgcccgcaac-3’ |
| Zwf-2-Forward | 5’- gttaacttaaggagaatgactatctgcgcttatccttta-3’ |
| Zwf-1-Forward | 5’- taaaggataagcgcagatagtcattctccttaagttaac-3’ |
| Zwf-2-Reverse | 5’- ggacatgatcgagcgttgcc-3’ |
| epd-1-Forward | 5’- gcgcacgcaacgtcagcg -3’ |
| epd-2-Forward | 5’- gaaaaccttgcaggagatctgacgcaagcagcgtctgc -3’ |
| epd-1- Reverse | 5’- gcagacgctgcttgcgtcagatctcctgcaaggttttc -3’ |
| epd-2- Reverse | 5’- tgttactggatgatgaaaag -3’ |
| glpD-1-Forward | 5’-aaaggggacagtataaagc-3’ |
| glpD-2-Forward | 5’-ctggccaccttacattaaagctgccctcattcactttc-3’ |
| glpD-1- Reverse | 5’-gaaagtgaatgagggcagctttaatgtaaggtggccag-3’ |
| glpD-2- Reverse | 5’-ttcgcaccggtctatttatg-3’ |
| mgsA-1-Forward | 5’- cggtttattcagcagaaccc -3’ |
| mgsA -2-Forward | 5’- gttaactacggatgtacatttattgcacaggtggcaaacg -3’ |
| mgsA -1- Reverse | 5’- cgtttgccacctgtgcaataaatgtacatccgtagttaac -3’ |
| mgsA -2- Reverse | 5’- ctgtatatctccccaccgctg -3’ |
| gldA-1-Forward | 5’- cgcaatgcgcggatcaccac -3’ |
| gldA -2-Forward | 5’- ctcatctctaaaggagcaatttactccaaactcccggcttg -3’ |
| gldA -1- Reverse | 5’- caagccgggagtttggagtaaattgctcctttagagatgag -3’ |
| gldA -2- Reverse | 5’- tcagactatgagtcgtgacgc -3’ |
| ldhA-1-Forward | 5’-cgccgggagctgtggcagc-3 |
| ldhA -2-Forward | 5’-catcactggagaaagtctttcttgccgctcccctgcaac-3’ |
| ldhA -2- Reverse | 5’-gttgcaggggagcggcaagaaagactttctccagtgatg-3’ |
| ldhA -1- Reverse | 5’-catcactggagaaagtctttcttgccgctcccctgcaac-3’ |
| frdBC-1-Forward | 5’- tgcgatgcacaatatcgttg -3’ |
| frdBC -2-Forward | 5’- gccaataagaaggagaaggcgaaaggagcctgagatgatcaatcc -3’ |
| frdBC -1- Reverse | 5’- ggattgatcatctcaggctcctttcgccttctccttcttattggc -3’ |
| frdBC -2- Reverse | 5’- tgccggtaatggcaacgaag -3’ |
| pflB-1-Forward | 5’- caaaaccgttggtgtccaga -3’ |
| pflB-2-Forward | 5’- ccacttaagaaggtaggtgttacttagatttgactgaaatcgtacag -3’ |
| pflB-1- Reverse | 5’- ctgtacgatttcagtcaaatctaagtaacacctaccttcttaagtgg -3’ |
| pflB-2- Reverse | 5’- ggggactaaacgtcctacaaac -3’ |
| frmAB-1-Forward | 5’- gctgttctgcgtaaaaatcatc -3’ |
| frmAB-2-Forward | 5’- cccagggtaaaggcttgatggcagaatttcattactaaatc -3’ |
| frmAB-1- Reverse | 5’- gatttagtaatgaaattctgccatcaagcctttaccctggg -3’ |
| frmAB-2- Reverse | 5’- atgggctgatggcagaagtg -3’ |
| adhE-1-Forward | 5’- tctgtttttgtggccgtaaag -3’ |
| adhE -2-Forward | 5’- ctcttgcctgtacactgacgctgagaaaaaagcgaaaaaatcc -3’ |
| adhE -1- Reverse | 5’- ggattttttcgcttttttctcagcgtcagtgtacaggcaagag -3’ |
| adhE -2- Reverse | 5’- ggtaaaccagctatcggtgtag -3’ |
| ptaI-1-Forward | 5’- tctgggcaaactgggcgtgc -3’ |
| ptaI-2-Forward | 5’- ctggatcaacgatttcaatccgacgatctcttccatcagc -3’ |
| ptaI -1- Reverse | 5’- gctgatggaagagatcgtcggattgaaatcgttgatccag -3’ |
| ptaI -2- Reverse | 5’- ccgcagcgtcgtactgcagc -3’ |
| acs-1-Forward | 5’- gactaactccctgaaatgcgC -3’ |
| acs-2-Forward | 5’- gccttgaccgccatctgcaagcttaatcacggggaggaacc -3’ |
| acs-1-Reverse | 5’- ggttcctccccgtgattaagcttgcagatggcggtcaaggc -3’ |
| acs-2-Reverse | 5’- atgacattgctcgcccctatg -3’ |
| fucO-1-Forward | 5’-ggtcaaccagcggcaacgaa-3’ |
| fucO-1-Reverse | 5’- cggcattatcacatcagcgcatccttctccttgttgctttacgaaattac -3’ |
| fucO-2-Forward | 5’-gtaatttcgtaaagcaacaaggagaaggatgcgctgatgtgataatgccg-3’ |
| fucO-2-Reverse | 5’- ctgttttacgcgtatgacagtctcc -3’ |
| yqhD-1-Forward | 5’-gacggctggaaatgctcatcttc -3’ |
| yqhD -2-Forward | 5’-cgaaagtttgaggcgtaaaaagctacttgctccctttgctgggcc -3’ |
| yqhD -1-Reverse | 5’-ggcccagcaaagggagcaagtagctttttacgcctcaaactttcg -3’ |
| yqhD -2-Reverse | 5’-cgccagtttcatcaatcaggcg -3’ |
| aldA-1-Forward | 5’-ggacaatcccgatgcaccacg -3’ |
| aldA-2-Forward | 5’-cctccgcctcttttactcagggcgactcctgtgatttatatg -3’ |
| aldA-1-Reverse | 5’-catataaatcacaggagtcgccctgagtaaaagaggcggagg -3’ |
| aldA-2-Reverse | 5’-actggcacccagtcactgg -3’ |

| Zwf-N20-Forward | 5’-tagcaaaacgcagcgccaacgttttagagctagaaatagcaag-3’ |
| --- | --- |
| Zwf-N20-Forward | 5’-gttggcgctgcgttttgctaactagtattatacctaggactg-3’ |
| epd-N20-Forward | 5’-gacgggaattatgcaattcggttttagagctagaaatagcaag-3’ |
| epd-N20-Forward | 5’-cgaattgcataattcccgtcactagtattatacctaggactg-3’ |
| glpD-N20-Forward | 5’-acgtgaagtgctactgaaaagttttagagctagaaatagcaag-3’ |
| glpD -N20-Forward | 5’-ttttcagtagcacttcacgtactagtattatacctaggactg-3’ |
| mgsA-N20-Forward | 5’-tcgtgcggcacggcatttaggttttagagctagaaatagcaag-3’ |
| mgsA-N20-Forward | 5’-ctaaatgccgtgccgcacgaactagtattatacctaggactgagc-3’ |
| gldA-N20-Forward | 5’-ctttttcgccttcttccagcgttttagagctagaaatagcaag-3’ |
| gldA-N20-Forward | 5’-gctggaagaaggcgaaaaagactagtattatacctaggactgagc-3’ |
| ldhA-N20-Forward | 5’-aggcggcttcgttcaacagagttttagagctagaaatagcaag-3’ |
| ldhA-N20-Forward | 5’-tctgttgaacgaagccgcctactagtattatacctaggac-3’ |
| frdBC-N20-Forward | 5’-agttgatgcaaccggagaacgttttagagctagaaatagcaag-3’ |
| frdBC-N20-Forward | 5’-gttctccggttgcatcaactactagtattatacctaggac-3’ |
| pflB-N20-Forward | 5’-tgcgttcggaacgatagagagttttagagctagaaatagc-3’ |
| pflB-N20-Forward | 5’-tctctatcgttccgaacgcaactagtattatacctaggactg-3’ |
| ptaI-N20-Forward | 5’-acgcagcattccgcacatgcgttttagagctagaaatagcaag-3’ |
| ptaI-N20-Forward | 5’-gcatgtgcggaatgctgcgtactagtattatacctaggactgagc-3’ |
| acs-N20-Forward | 5’-gtacagcaagtaactgtgtcgttttagagctagaaatagc-3’ |
| acs-N20-Forward | 5’-gacacagttacttgctgtacactagtattatacctaggactg-3’ |
| fucO-N20-Forward | 5’-catgacatgcggtagcaggagttttagagctagaaatagcaag-3’ |
| fucO-N20-Forward | 5’-tcctgctaccgcatgtcatgactagtattatacctaggac-3’ |
| yqhD-N20-Forward | 5’-attcaggaccgtttcgcagagttttagagctagaaatagcaag-3’ |
| yqhD-N20-Forward | 5’-tctgcgaaacggtcctgaatactagtattatacctaggac-3’ |
| aldA-N20-Forward | 5’-gtacagcaagtaactgtgtcgttttagagctagaaatagc-3’ |
| aldA-N20-Forward | 5’-gacacagttacttgctgtacactagtattatacctaggactg-3’ |

**Table S7. List of plasmids.**

| **Plasmids** | **Short discription** | **Source** |
| --- | --- | --- |
| pET28a-RpiB | pET28a based vector, carrying *rpiB* gene from *Brucella abortus*; Kan^r^ | This study |
| pET28a-EcHAD | pET28a based vector, carrying *yidA* gene from *Escherichia coli* MG1655; Kanr | This study |
| pET28a-NbIMP | pET28a based vector, carrying *impase* gene from *Nocardia brasiliensis*; Kan^r^ | This study |
| pET28a-TmHAD | pET28a based vector, carrying *had* gene from *Thermotoga,* Kan^r^ | This study |
| pET28a-XlHAD | pET28a based vector, carrying *had* gene from *Xenopus laevis*; Kan^r^ | This study |
| pET28a-pfHAD | pET28a based vector, carrying *had* gene from *Plasmodium falciparum*; Kan^r^ | This study |
| pET28a-nagD | pET28a based vector, carrying *had* gene from *Corynebacterium*; Kan^r^ | This study |
| pET28a-CpHAD | pET28a based vector, carrying *had* gene from *Candida parapsilosis*; Kan^r^ | This study |
| pET28a-pk-1 | pET28a based vector, carrying *pk* gene fromLactobacillus baoqingensis; Kan^r^ | This study |
| pET28a-pk-2 | pET28a based vector, carrying *pk* gene from Aspergillus niger; Kan^r^ | This study |
| pET28a-pk-3 | pET28a based vector, carrying *pk* gene from *Pseudomonas congelans*; Kan^r^ | This study |
| pET28a-pk-4 | pET28a based vector, carrying *pk* gene from *Bifidobacterium adolescentis;* Kanr | This study |
| pET28a-pk-5 | pET28a based vector, carrying *pk* gene from *Agrobacterium tumefaciens complex*; Kan^r^ | This study |
| pET28a-pk-6 | pET28a based vector, carrying *pk* gene from *Nitrolancea hollandica*; Kan^r^ | This study |
| pET28a-pk-7 | pET28a based vector, carrying *pk* gene from *Sphingomonas sp*; Kan^r^ | This study |
| pET28a-fls | pET28a based vector, carrying *fls* gene from *Methanosarcina thermophila*; Kane^r^ | This study |
| pET28a-PsLrhi | pET28a based vector, carrying *PsLrhi* gene from *Pseudomonas stutzeri*; Kane^r^ | ^17^ |
| pET28a-gals | pET28a based vector, carrying *gals* gene; Kane^r^ | ^14^ |
| pET28a-ack | pET28a based vector, carrying *ackA* gene from *E. coli* Mg1655; Kan^r^ | ^18^ |
| pET28a-TIM | pET28a based vector, carrying *tpiA* gene from *E. coli* Mg1655; Kanr | This study |
| pACYC-DuetI-pk-4- *Pslrhi* | pET28a based vector, carrying *pk4* and *Pslrhi* ; Cm^r^ | This study |
| pACYC-DuetI-pk4-fls | pET28a based vector, carrying *pk4* and *fls*gene from *Myceliophthora thermophila*; Cm^r^ | This study |
| pTargetT-ldhA | pMB1 *ldhA* sgRNA-pMB，Spec^r^ | This study |
| pTargetT-adhE | pMB1 *adhE* sgRNA-pMB1，Spec^r^ | This study |
| pTargetT-pflB | pMB1 *pflB* sgRNA-pMB1，Spec^r^ | This study |
| pTargetT-fuco | pMB1 *fucO* sgRNA-pMB1，Spec^r^ | This study |
| pTargetT-frdBC | pMB1 *frdBC* sgRNA-pMB1，Spec^r^ | This study |
| pTargetT-yqhD | pMB1 *yqhD* sgRNA-pMB1，Spec^r^ | This study |
| pTargetT-aldA | pMB1 *aldA* sgRNA-pMB1，Spec^r^ | This study |
| pTargetT-patI | pMB1 *ptaI* sgRNA-pMB1，Spec^r^ | This study |
| pTargetT-acs | pMB1 *acs* sgRNA-pMB1，Spec^r^ | This study |
| pTargetT-zwf | pMB1 *zwf* sgRNA-pMB1，Spec^r^ | This study |
| pTargetT-epd | pMB1 *epd* sgRNA-pMB1，Spec^r^ | This study |
| pTargetT-mgsA | pMB1 *mgsA* sgRNA-pMB1，Spec^r^ | This study |
| pTargetT-glpD | pMB1 *glpD* sgRNA-pMB1，Spec^r^ | This study |
| pTargetT-gldA | pMB1 *gldA* sgRNA-pMB1，Spec^r^ | This study |
| pFN-Cas9-K | repA101(Ts) kan Pcas-cas9 ParaB-Red lacIq Ptrc-sgRNA-pMB1 pSC10 replication, temperature sensitive replication origin, Para BAD-drivenI-SceI gene, red recombinase expression plasmid, lac-inducible expression; Kan^r^ | This study |

**Table S8. List of synthesized genes and their optimized sequences.**

| **Gene** | **Sequence** |
| --- | --- |
| **rpiB** | ATGAAAGTTGCTGTTGCTGGTGACTCTGCTGGTGAAGGTCTGGCTAAAGTTCTGGCTGACCACCTGAAAGACCGTTTCGAAGTTTCTGAAATCTCTCGTACCGACGCTGGTGCTGACGCTTTCTACGCTAACCTGTCTGACCGTGTTGCTTCTGCTGTTCTGGACGGTACCTACGACCGTGCTATCCTGGTTTGCGGTACCGGTATCGGTGTTTGCATCGCTGCTAACAAAGTTCCGGGTATCCGTGCTGCTCTGACCCACGACACCTACTCTGCTGAACGTGCTGCTCTGTCTAACAACGCTCAGATCATCACCATGGGTGCTCGTGTTATCGGTGCTGAAGTTGCTAAAACCATCGCTGACGCTTTCCTGGCTCAGACCTTCGACGAAAACGGTCGTTCTGCTGGTAACGTTAACGCTATCAACGAAGTTGACGCTAAATACAACAAATTC |
| **pfHAD** | ATGCACGAAATTGTAGATAAGAATGGTAAGAAAGTTCAAAAGAATAATTTGAATGATGAAATAAAAATAATCTTTACGGATTTAGATGGAACATTGTTAAATAGTGAGAATAAGGTTTCAGAACAGAATTTGGAGAGTTTAATAAGAGCTCAAGAAAAAGGCATAAAGGTTGTTATAGCAACAGGTAGATCTATATTTTCTGTAGAGAGTGTTATAGGAGAGCATGTAAAAAAGAATAGAATAAGTTTATTACCAGGGATATATATGAATGGATGTGTAACATTTGATGAAAAAGGTTCAAGGGTGATAGATAGGATTATGAACAATGACTTGAAAATGGAGATACATGAATTTTCTAAACAAATAAATATATCAAAATATGCTATATGGTTTTGTTTAGAAAAAACATATTGTTTTGAAATAAATGATTGTATACGTGAATATATGGAGGTTGAAGCATTAAATCCTGATGTTATTGAAGATAATATGTTAGAAGGTTTGACAGTATATAAAGTATTATTTTCATTACCAGAAAATATATTAGAAAATACGTTAAAATTATGTAGAGAGAAATTTTCTCATCGTATTAATGTAGCTAATACTTTTCAAAGTTATGTTGAATTATTTCATCAACATACTAATAAATTCGAAGGTGTAAAAGAAATTTGTAAATATTATAATATAAGTCTAAACAATGCGCTAGCTATGGGAGATGGAGAAAATGATATTGAAATGTTAAGTGGTTTAACACATTCAGTGGGTGTACATAATGCTTCAGAAAAAGTAAAAAATTCAGCTGCTTATGTTGGACCTTCGAATAATGAACATGCTATATCTCATGTCTTGAAGACATTCTGTGACATATAA |
| **yidA** | ATGGCTATTAAACTCATTGCTATCGATATGGATGGCACCCTTCTGCTGCCCGATCACACCATTTCACCCGCCGTTAAAAATGCGATTGCCGCAGCTCGCGCCCGTGGCGTGAATGTCGTGCTAACGACGGGTCGCCCGTATGCAGGTGTGCACAACTACCTGAAAGAGCTGCATATGGAACAGCCGGGCGACTACTGCATTACTTATAACGGCGCGCTGGTACAGAAGGCCGCTGATGGTAGCACCGTGGCGCAAACTGCTCTCAGCTATGACGACTATCGTTTCCTGGAAAAACTCTCTCGCGAAGTCGGTTCTCATTTCCACGCCCTGGACCGCACCACGCTGTACACCGCCAACCGTGATATCAGCTACTACACGGTGCATGAATCCTTCGTTGCCACCATTCCGCTGGTGTTCTGCGAAGCGGAGAAAATGGACCCCAATACCCAGTTCCTGAAAGTGATGATGATTGATGAACCCGCCATCCTCGACCAGGCTATCGCGCGTATTCCGCAGGAAGTGAAAGAGAAATATACCGTGCTGAAAAGTGCGCCGTACTTCCTCGAAATCCTCGATAAACGCGTTAACAAAGGTACGGGGGTGAAATCACTGGCCGACGTGTTAGGTATTAAACCGGAAGAAATCATGGCGATTGGCGATCAGGAAAACGATATCGCAATGATTGAATATGCAGGCGTCGGTGTGGCGATGGATAACGCTATTCCTTCAGTGAAAGAAGTGGCGAACTTTGTCACCAAATCTAACCTTGAAGATGGCGTGGCGTTTGCTATTGAGAAGTATGTGCTGAATTAA |
| **Impase** | ATGGCTGACCCGTGGCAGGAATGCATGGACTACGCTGTTACCCTGGCTCGTCAGGCTGGTGAAGTTGTTTGCGAAGCTATCAAAAACGAAATGAACGTTATGCTGAAATCTTCTCCGGTTGACCTGGTTACCGCTACCGACCAGAAAGTTGAAAAAATGCTGATCTCTTCTATCAAAGAAAAATACCCGTCTCACTCTTTCATCGGTGAAGAATCTGTTGCTGCTGGTGAAAAATCTATCCTGACCGACAACCCGACCTGGATCATCGACCCGATCGACGGTACCACCAACTTCGTTCACCGTTTCCCGTTCGTTGCTGTTTCTATCGGTTTCGCTGTTAACAAAAAAATCGAATTCGGTGTTGTTTACTCTTGCGTTGAAGGTAAAATGTACACCGCTCGTAAAGGTAAAGGTGCTTTCTGCAACGGTCAGAAACTGCAGGTTTCTCAGCAGGAAGACATCACCAAATCTCTGCTGGTTACCGAACTGGGTTCTTCTCGTACCCCGGAAACCGTTCGTATGGTTCTGTCTAACATGGAAAAACTGTTCTGCATCCCGGTTCACGGTATCCGTTCTGTTGGTACCGCTGCTGTTAACATGTGCCTGGTTGCTACCGGTGGTGCTGACGCTTACTACGAAATGGGTATCCACTGCTGGGACGTTGCTGGTGCTGGTATCATCGTTACCGAAGCTGGTGGTGTTCTGATGGACGTTACCGGTGGTCCGTTCGACCTGATGTCTCGTCGTGTTATCGCTGCTAACAACCGTATCCTGGCTGAACGTATCGCTAAAGAAATCCAGGTTATCCCGCTGCAGCGTGACGACGAAGACCTCG |
| **tm1254** | ATGGAAGCTGTTATCTTCGACATGGACGGTGTTCTGATGGACACCGAACCGCTGTACTTCGAAGCTTACCGTCGTGTTGCTGAATCTTACGGTAAACCGTACACCGAAGACCTGCACCGTCGTATCATGGGTGTTCCGGAACGTGAAGGTCTGCCGATCCTGATGGAAGCTCTGGAAATCAAAGACTCTCTGGAAAACTTCAAAAAACGTGTTCACGAAGAAAAAAAACGTGTTTTCTCTGAACTGCTGAAAGAAAACCCGGGTGTTCGTGAAGCTCTGGAATTCGTTAAATCTAAACGTATCAAACTGGCTCTGGCTACCTCTACCCCGCAGCGTGAAGCTCTGGAACGTCTGCGTCGTCTGGACCTGGAAAAATACTTCGACGTTATGGTTTTCGGTGACCAGGTTAAAAACGGTAAACCGGACCCGGAAATCTACCTGCTGGTTCTGGAACGTCTGAACGTTGTTCCGGAAAAAGTTGTTGTTTTCGAAGACTCTAAATCTGGTGTTGAAGCTGCTAAATCTGCTGGTATCGAACGTATCTACGGTGTTGTTCACTCTCTGAACGACGGTAAAGCTCTGCTGGAAGCTGGTGCTGTTGCTCTGGTTAAACCGGAAGAAATCCTGAACGTTCTGAAAGAAGTTCTGCTCGA |
| **xlapase** | ATGGCTCAGCAGGGTAACGGTAAATCTGTTCTGTTCGTTTGCCTGGGTAACATCTGCCGTTCTCCGATCGCTGAAGCTGTTTTCCAGAAACTGGTTACCGACGCTGGTATCTCTAAAGAATGGTCTATCGACTCTGCTGCTACCTCTGACTGGAACGTTGGTTCTTCTCCGGACTCTCGTGCTCTGAAATGCCTGAAATCTCACTCTATCGAAACCTCTCACCGTGCTCAGCAGATCACCCGTGACGACTTCCTGTCTTACGACTACATCCTGTGCATGGACGAATCTAACCTGCAGGACCTGAAACGTCGTGGTTCTCAGGTTCAGAACTGCAAAGCTAAAATCGAACTGCTGGGTTCTTACGACCCGCAGAAACAGCTGATCATCCAGGACCCGTACTACGGTCGTGACGAAGACTTCGAAACCGTTTACCAGCAGTGCATCCGTTGCTGCAAATCTTTCCTGGAAAAATCTTCTCTCG |
| **nagD** | CATGACAGTGAACATTTCATATCTGACCGACATGGACGGCGTCCTCATCAAAGAGGGCGAGATGATTCCGGGTGCAGATCGTTTTCTTCAGTCTCTCACCGATAACAATGTGGAGTTTATGGTTTTGACCAACAACTCCATTTTCACCCCGAGGGATCTTTCTGCACGTCTTAAGACTTCCGGTTTGGATATCCCGCCGGAGCGTATTTGGACTTCTGCAACCGCCACTGCTCACTTCCTGAAATCCCAGGTCAAGGAGGGCACAGCCTATGTTGTTGGGGAGTCCGGTCTGACCACTGCGTTGCATACCGCGGGTTGGATTTTGACGGATGCAAATCCTGAGTTTGTTGTCCTGGGCGAAACCCGCACGTATTCCTTCGAGGCAATCACCACTGCTATAAATCTGATTTTGGGCGGCGCTCGCTTTATTTGCACCAACCCGGATGTAACAGGACCTTCACCAAGTGGCATTTTGCCTGCTACTGGCTCTGTCGCAGCGCTTATTACCGCAGCTACAGGCGCTGAGCCTTATTACATCGGTAAGCCAAACCCTGTGATGATGCGCAGTGCGCTGAACACCATCGGGGCGCATTCCGAGCACACTGTCATGATCGGCGACCGCATGGACACCGACGTGAAATCTGGTTTGGAAGCCGGCCTGAGCACCGTGCTGGTTCGAAGCGGAATCTCCGACGACGCCGAGATCCGCCGCTACCCCTTCCGCCCAACTCACGTGATCAATTCCATCGCCGATCTTGCCGATTGCTGGGACGATCCATTCGGTGACGGTGCATTTCACGTACCAGATGAGCAGCAGTTCACTGACTAGCTCGA |
| **Cppho** | ATGTCTGTTAAAATCACCGAAAAATCTCAGGTTCAGAACCTGATCCTGGACAAATACGACTACTTCCTGTTCGACTGCGACGGTGTTCTGTGGCTGGGTGACCACCTGCTGCCGTCTATCGGTGAAGCTCTGGACTACCTGAAACAGCAGAACAAAACCGTTATCTTCGTTACCAACAACTCTACCAAATCTCGTACCGACTACCTGTCTAAATTCAACAAAATGGGTATCTCTAACATCACCAAATCTGAAATCTTCGGTTCTTCTTTCGCTTCTGCTGTTTACGTTGAAAAAATCCTGAAACTGCCGAAAGACAAAAAAGTTTGGGTTCTGGGTGAAGAAGGTATCGAAAAAGAACTGCACGAACTGGGTTACTCTACCGTTGGTGGTACCGACCCGAAACTGGTTAAAGAAGGTGTTAAATTCGACCCGAACACCACCCTGTTCGACAACCTGGACCCGAACGTTGGTTGCGTTGTTTGCGGTCTGACCTTCAACATCAACTACCTGAAACTGTCTCTGACCATGCAGTACCTGCTGAAAGACAACAAATCTATCCCGTTCATCGCTACCAACATCGACTCTACCTTCCCGATGAAAGGTAAACTGCTGATCGGTGCTGGTTCTATCATCGAAACCGTTGCTTACGCTTCTGGTCGTCAGCCGGACGCTATCTGCGGTAAACCGAACCAGTCTATGATGAACTCTATCAAAGCTCAGCTGCCGGGTCTGGAAAAAAACCCGAAAAAAGGTCTGATGATCGGTGACCGTCTGAACACCGACATGAAATTCGGTCGTGACGGTGGTCTGGACACCATGCTGGTTCTGACCGGTATCGAAACCGAATCTAACGTTAAACAGCTGTCTAAAGAAGACGCTCCGACCTACTACATCGAAAAACTGGGTGACATCTACGAATTCACCCAC |
| **PK 1** | ATGGCTACAGATTATTCATCTCAAGCATACTTTGACAAGATGACCGCGTATTGGCGCGCGGCAAATTATATTTCTGTTGGACAACTATATCTGGTGGCGAACCCATTGCTGCGCCGGCCTTTGGAATCGGATGATGTGAAATACTATCCGATCGGGCACTGGGGCACGATCTCAGGACAAAACTTCATTTATACGCATTTGAACCGGGTCATCAATAAATATGACTTGAACATGTTTTATTTGGAAGGTCCAGGTCATGGCGGCCAAGTTATGCTGTCGAATGCCTACCTTGATGGCACGTATTCTGAGAAATATCCGAACATTTCTCAAGATGAAAAAGGGATGCAGATCCTATTCAAGCGCTTCTCCTTCCCAGGCGGGGCAGCCAGTCACGCCAATGCGCAGATCCCAGGGTCTATTCATGAAGGCGGCGAGTTGGGGTACACGCTGAGTCATGCCACTGGGGCAGTTTTAGATAATCCTGACGTGATCGCCGCTGCGGTCACTGGGGATGGTGAAACTGAAACCGGACCATTGGCAGCTTCGTGGTTCTCAAATACCTTCATCAACCCGATCAACGATGGGGCGGTATTGCCGATCGTGCATATGAACGGCTTCAAGATCTCCAACCCAACGATCTTGTCGCGTAAATCGGATGATGACTTACGCAAGTACTTCGAAGGGATGGGATGGGACCCATACTTCGTTGAAGGTGATGATCCGAATAAGCTGAATCCGATCATGGCCAAGACGATGGATACTGCGATCGAAAAGATCCAAGCGATCCAAAAGCATGCCCGTGAAACCGGCGATGCCACTATGCCACACTGGCCAGTGTTGATCGTACGCACGCCGAAAGGCTGGACTGGTCCTAAGACATGGAATGGCGAGCCGATCGAAGGCTCCTTCCGTGCGCACCAGATCCCGATCCCAGTCAAGCGCGACGACATGACCCACAAGGAATCTTTGGAAGATTGGCTGAAGTCTTATCATCCAGAAGAGCTGTTTGATGAAAACGGCACGCTGATCCCAGAACTGCAAGCGTTGACGCCTAAAGGCGATAAGCGGATGGCGGCTAACCCGATCACCAATGTCGGCTATCATGCAAAGCCATTAGTGCTCCCTGATTTCAAAGATTATGCCTTAGACAACTCCAAACGCGGCCAAAACGTCAAGCAAGACATGATCATCTGGTCTGATTACCTGCGGGATCTGATCAAGCTCAACCCGCAAAACTTCCGGGTATTCGGTCCCGATGAAACCATGTCCAATCGCTTGTATGGTTTGTTTGAAGCGACTGACCGGCAGTGGTTGGAGCCGATCAACCAGCATGGTTGCGATGAAAAAATGGCGCCAGCTGGGCGGATCATTGACTCACAGCTGTCAGAACATCAGGCTGAAGGCTTCTCTGAAGGCTACACTTTGACTGGTCGGCACAGCCTGTTCACGTCATATGAAGCCTTCTTGCGGGTCGTTGATTCGATGCTGACGCAGCACTTCAAGTGGATGCGTGAAGCGGCTAAGGAGCAGTGGCACAAACCATATCCTTCTTTGAACGTGGTGTCGACGTCAACATCATTCCAACAAGACCACAACGGCTACACGCACCAAGATCCTGGTATTTTAACGCACTTGGCTGAAAAGAAAGGTGAGTTCATTCGCGAGTATCTGCCAGCAGATGCCAACTCGTTGTTAGCGATCTCGCCGAAAGTCTTCAGCAGTCAAAACACCATCAACTTGCTGATCACCTCTAAGCAGCCGCGGCCACAATTTTATTCCATTGAAGAAGCCGAAGTGTTGGCTGAAAATGGCTTAAAGCGTATCGATTGGGCTTCTAACGACGATGGTGTCGCACCCGATGTCGTGATCGCAGCAGCTGGGACGGAACCGAACATGGAAAGCTTAGCGGCCATCAATTTGCTGCATGACAACTTCCCAGAGCTGAAGATTCGCTTTATCAACGTGGTCGATCTCTTGAAGTTGAAGAGTCCAGACCGCGATCCGCGGGGCTTGTCTGACGAAGAATTTGATCGCTACTTTACCACGGATAAGCCGATCTTCTTTGGCTTCCACGGTTTTGAAGATTTGATTCGGGACATCTTCTTTGATCGGCATAACCATAATTTACGCGTGCACGGGTATCGTGAAGAAGGGGCCATTACCACACCGTTTGATATGCGGGTGGTCAATGAACTTGACCGCTTCCATCTGGCTAAAGATATTATCGCCCATGTGCCAGGTTATGAACAAAAAGCGGCGGCCTTCATCCAAAAAATGGATGATACGCTGCAATTCCACCACGATTATATTCGCCAAACCGGCGAAGATATTCCAGAAGTGCAAGCTTGGACTTGGCAAGCGATCAAGTGA |
| **PK2** | ATGCCTGGAGAGGTCATCGACAGGCCGAATCCCAAGGCCGAGCCTTCACACATTCCCGATCTTGTCAATCAATTGCAGGTCAAACTTCAAGAGACGAGTTTGGAGGAAACTGATTACAATGCCCTGCTGAAATTCCGCCGTGCAGCGGCCTACATTGCTGCTGCAATGATCTTTCTCCAAGACAATGTGCTGCTGAAGCAGAATCTAAAGCATGAGGACATCAAGCCCAGGCTTCTTGGCCACTGGGGAACATGTCCCGGGTTGATTCTTGTATACTCTCACTTGAACTACATCATCAGAAAGCATAATCTGGATATGTTGTATGTCGTCGGGCCTGGCCACGGCGCGCCAGCTATTTTGGCCTCACTGTGGCTTGAGGGCTCTTTAGAGAAATTCTACCCCCACTACTCACGAGACATGGATGGTCTCCATGAGCTCATCTCGACCTTCAGCACAAGTGCTGGATTACCAAGCCATATCAATGCGGAAACTCCCGGTGCGATCCATGAAGGTGGTGAACTGGGTTACGCGCTGGCTGTCTCTTTTGGTGCTGTTATGGACAATCCCGACATGATCGTCACCTGCGTGGTTGGTGACGGGGAAGCAGAAACCGGTCCTACCGCGACGTCCTGGCATGCAATCAAGTACATTGACCCCGCAGAATCAGGTGCCGTCCTGCCGATTCTCCACGTTAATGGCTTTAAGATCAGCGAGCGCACCATCTATGGCTGCATGGACAACAAAGAGCTGGTCTCCCTCTTCACGGGTTATGGATACCAGGTGCGCATTGTTGAGAACCTGGATGACATCGACGCAGATCTCCATAGCTCTATGATGTGGGCAGTTGAGGAGATCCACAAGATCCAAAAAGCGGCGCGTTCCGGCAAGCCAATAATGAAGCCTAGATGGCCAATGATTGTTTTGCGCACACCGAAGGGCTGGTCAGGACCTAAAGAGCTCCACGGGTCATTCATAGAGGGATCTTTCCACTCACATCAGGTTCCTCTACCTAATGCAAAGAAGGATAAAGAGGAGCTTCAAGCTCTGCAGAAATGGCTGTCCTCGTATAATCCGCACGAACTTTTCACTGAGACGGGAGACATCATTGACGACATCAAGTCAGTGATCCCTCTGGAAGACACCAAGAAGCTTGGGCAGCGAGCAGAAGCCTACAAGGGCTACAGGGCACCCGATCTCCCAGACTGGCGCAAGTTTGGCGTAGAAAAGGGCTCCCAGCAGAGCGCTATGAAAACAATTGGAAAGTTCATTGACCAAGTGTTTACCCAAAATCCTCATGGCGTCCGTGTATTTTCGCCAGACGAGCTAGAGAGCAACAAGCTGGATGCAGCACTGGCGCACACGGGAAGGAACTTTCAGTGGGATCAATTCTCGAATGCCAAAGGCGGCCGCGTCATCGAGGTGCTCAGTGAGCACCTGTGCCAGGGCTTTATGCAGGGATACACGTTGACGGGCCGGGTGGGCATTTTCCCATCGTACGAAAGCTTCTTGGGAATCATCCATACCATGATGGTGCAATATGCCAAATTTAACAAAATGGCTCAAGAGACGACCTGGCATAAGCCGGTTAGTAGCATCAACTATATCGAAACGAGTACGTGGGCTCGTCAGGAGCACAATGGATTCTCTCACCAGAACCCCTCCTTTATCGGAGCTGTGCTCAGGCTGAAGCCCACCGCCGCGCGAGTTTATCTGCCACCTGATGCTAACACATTTTTGACCACCCTTCACCACTGTCTCAAGTCCAAGAATTATGTCAACCTCATGGTAGGTTCAAAGCAGCCAACTCCCGTGTACTTGAGCCCCGAGGAAGCAGAGAGCCACTGCCGAGCCGGCGCATCGATCTGGAGATTCTGTAGTACCGACAATGGGCTGAACCCGGATGTCGTGCTGGTTGGCATTGGAGTAGAGGTGATGTTCGAGGTCATCTACGCGGCGGCCATCCTCCGCAAGCGTTGTCCAGAACTCCGAGTGCGTGTGGTCAACGTGACCGACTTGATGATTCTGGAGAAGGAAGGTCTACATCCACATGCATTGACGACCGAAGCTTTCGACAGCCTGTTTGGCTCGGACCGGCCGATACACTTCAACTACCACGGATACCCGGGCGAGCTCAAAGGTCTGCTCTTTGGGCGGCCCCGCCTGGACCGAGTTTCAGTGGAAGGATACATGGAGGAAGGAAGCACGACGACGCCGTTCGATATGATGTTGTTGAACCGCGTCTCACGATACCACGTGGCGCAGGCAGCCGTGATCGGGGCGTCCAGACGGAATGAGAAGGTTCAAGTTCGGCAGCACGAACTAGTCAGCGAATTCGGCCACAACATTGTGGAGACACGCAAATACATTCTGGCCAACCGCAAAGACCCGGATGATACGTATGATATGCCCTCGTTTGAATGA |
| **PK 3** | ATGTATCACTTCAACTCAACGCTTTCCAGTGCCGAACAGTCGGACGCTGAATCGGCCACGCAGCCTGAACGGCCGGGGAGCGAAGCAGGCCCCTTGGACGCCGATCTGCTGTCACGCCTGCATCGTTACTGGAATGCCGCCAATTACCTGTGCGTTGGCCAGATCTATCTGAAGGCCAATGCACTGCTGCACGAGCCGTTGCTGGCCGAGCACATCAAGCCACGACTGCTGGGGCATTGGGGTACGTCGGTTGGGCAGAACTTTATCTATGTGCACCTCAATCGCCTGATTTGCGACCGACGCATCGAGACGATTTTCATCTCCGGTCCAGGTCATGGCGGCCCGACCATGAACGCCTGCGCCTGGCTTGAAGGCACGTACAGCGAGGTTCACCCCGACATTCCGGCCGATGAAGAGGGCATGCTGGCTTTCTTTCGCAGCTTTTCCACCCCCGGCGGTATCCCCAGCCATTGCGGCCCGCACACACCCAACTCCCTGCATGAAGGCGGAGAGCTGGGCTATTCGCTGATGCACGCCTTTGGCGCTGTCTTCGACAATCCCCACCTGCTGGTCGCCTGCGTCATTGGTGACGGGGAAGCTGAAACCTGTCCGCTGGAAGGCAGCTGGAAAAGTGTGCACTTTCTCGATCCCCGTCGGGATGGTGCCGTGCTGCCGATCCTGCACCTCAATGGCTACAAGATTTCCGGGCCGACGGTCGAAGCACGTTTGCCGGACGAGCAGTTGATTGAGCTGTACCGAGGGCGGGGTTATCAGCCCGTTATCGTCGCGGGGGATGACCTGCCAGGCATGCATCAACGCTTCGCAGCGGCGCTCAATATCTGTCATGACGCGATCCGCGAACAGCAGGCGCGCTCCAGAGCGGAAGGGGGCGCTGCGCGTGCGCGGTGGCCGATGATTATCCTGCGCAGTCCCAAGGGCTGGACCGGACCGAAAGTCGTGGACGGTGTGCCGGTGGAGGGCACCTTTCGCGCTCATCAGGTGCCGTTGGCCAATGTCATCGGTAACCCGGAGCACCTCCAGCAACTGGAGCGCTGGCTGCGCAGTTACTCGCCCGAGACGTTGTTCGATCAGAACGGCCGGTTGCTGCCCGAGCTTCAGGCGTTGACCCCGCCGGCTGCATTGCGCATGGGCGCCGTACCTTACGTGAACGGCGGACGCGTGCTGGTGGCGCTGGACCTGCCGAACTTTGCCGACTATGGACTGGAAGTGCCGGGCCCCGGTCAGGTCATCGCCGAAGCACCACGTCGGCTGGGCGAATACCTGCGTGATGTGATGCGCAATAACCCGCACAACTTTCGTGTCTTCGGGCCTGACGAGACCAACTCTAACCGACTCAACGCCGTGTTCGAAGCCAGCAACCGCACTGCACCAGGCCCGTGTCTGGAGATTGACGATCACCTGGCAAGCGATGGTCGGGTGATGGAGGTGCTCAGTGAACACCTGTGTGAAGGGTGGCTGGAGGGTTATCTGCTCACCGGACGACACGGTATGTGGTCGACCTACGAAGCATTTGCTCAGGTGGTCGACTCCATGGTCACTCAGCACGCCAAATGGTTGCAGCAGAGCCGCGAGTTTGCCTGGCGTCGTCCGCTGGCCTCACTCAACATTCTGATCAGCAGCCATGCCTGGCGTAACGATCACAATGGTTTCAGCCATCAGTCCACCGGGTTCGTCGACAACGTCTTGCAGCGCCGAGCCGACGTGGTGAGGGTGTATTACCCGCCGGATTCCAATTGCCTGGTCAACGTTTTCGACCATTGTCTGCGCAGCCGCAACTACATAAACGTTGTGACCTGCGGCAAGCAACCGGACTTTCAATGGCTGGACTTCGACGCGGCGCTCAAACATTGCTCGCAAGGGGCGTCGATCTGGGACTTCGCAGGCAGTGACGACAGTGAGACGCCGGATGTCGTGCTGGGCTGTGCGGGCGATGTACCGACCACCGAAGCAGTGGCGGCAGCCTGGTTGCTGCAAAAACACGTTCCGGGTATTCGCGTGAGGCTGGTCAATGTCGTCGATCTGGGCATTCTGAGCGCCCCTGAAGATCGGCCTCATGGCATGGACCATGTGTCGTTCGAGGCCTTGTTTACCCGTGACGCGCCGGTGATGTTCGCCTTTCACGGTTCGACCTGGGTGATCCATTCCATGGTTCATGGCCGCGCCAACGAGGCGCGCTTTCATGTCCGTGGTTTCAGTGATCGTGGGACCACGACGACGCCATTCGACATGGTGGTGCTCAATCGCCTTAGCCGTTATCAGCTGGCCATCGACGCATTGAGCCACGTACCGCGTTTACGCGCGCAGAGTCTGGATGCGGTGCATTTTTTCGAAAGTCAGTTACGCAAGCATCACACCTGGATACGCGAACACTTCGAAGACATGCCGGAGATTCGCAATTGGTGCTGGACGGCTGATTTCAGCGAATCGCAGACGCCTCCGCCATTGGCCAAGGGGCACAGCCGAGGGCAGACGTTTACGGATGCGTGA |
| **PK 4** | ATGACGAGTCCTGTTATTGGCACCCCTTGGAAGAAGCTGAACGCTCCGGTTTCCGAGGAAGCTATCGAAGGCGTGGATAAGTACTGGCGCGCAGCCAACTACCTCTCCATCGGCCAGATCTATCTGCGTAGCAACCCGCTGATGAAGGAGCCTTTCACCCGCGAAGACGTCAAGCACCGTCTGGTCGGTCACTGGGGCACCACCCCGGGCCTGAACTTCCTCATCGGCCACATCAACCGTCTCATTGCTGATCACCAGCAGAACACTGTGATCATCATGGGCCCGGGCCACGGCGGCCCGGCTGGTACCGCTCAGTCCTACCTGGACGGCACCTACACCGAGTACTTCCCGAACATCACCAAGGATGAGGCTGGCCTGCAGAAGTTCTTCCGCCAGTTCTCCTACCCGGGTGGCATCCCGTCCCACTACGCTCCGGAGACCCCGGGCTCCATCCACGAAGGCGGCGAGCTGGGTTACGCCCTGTCCCACGCCTACGGCGCTGTGATGAACAACCCGAGCCTGTTCGTCCCGGCCATCGTCGGCGACGGCGAAGCTGAGACCGGCCCGCTGGCCACCGGCTGGCAGTCCAACAAGCTCATCAACCCGCGCACCGACGGTATCGTGCTGCCGATCCTGCACCTCAATGGCTACAAGATCGCCAACCCGACCATCCTGTCCCGCATCTCCGACGAAGAGCTCCACGAGTTCTTCCACGGCATGGGCTATGAGCCGTACGAGTTCGTCGCTGGCTTCGACAACGAGGATCACCTGTCGATCCACCGTCGTTTCGCCGAGCTGTTCGAGACCGTCTTCGACGAGATCTGCGACATCAAGGCCGCCGCTCAGACCGACGACATGACTCGTCCGTTCTACCCGATGATCATCTTCCGTACCCCGAAGGGCTGGACCTGCCCGAAGTTCATCGACGGCAAGAAGACCGAGGGCTCCTGGCGTTCCCACCAGGTGCCGCTGGCTTCCGCCCGCGATACCGAGGCCCACTTCGAGGTCCTCAAGAACTGGCTCGAGTCCTACAAGCCGGAAGAGCTGTTCGACGAGAACGGCGCCGTGAAGCCGGAAGTCACCGCCTTCATGCCGACCGGCGAACTGCGCATCGGTGAGAACCCGAACGCCAACGGTGGCCGCATCCGCGAAGAGCTGAAGCTGCCGAAGCTGGAAGACTACGAGGTCAAGGAAGTCGCCGAGTACGGCCACGGCTGGGGCCAGCTCGAGGCCACCCGTCGTCTGGGCGTCTACACCCGCGACATCATCAAGAACAACCCGGACTCCTTCCGTATCTTCGGACCGGATGAGACCGCTTCCAACCGTCTGCAGGCCGCTTACGACGTCACCAACAAGCAGTGGGACGCCGGCTACCTGTCCGCTCAGGTCGACGAGCACATGGCTGTCACCGGCCAGGTCACCGAGCAGCTTTCCGAGCACCAGATGGAAGGCTTCCTCGAGGGCTACCTGCTGACCGGCCGTCACGGCATCTGGAGCTCCTATGAGTCCTTCGTGCACGTGATCGACTCCATGCTGAACCAGCACGCCAAGTGGCTCGAGGCTACCGTCCGCGAGATTCCGTGGCGCAAGCCGATCTCCTCCATGAACCTGCTCGTCTCCTCCCACGTGTGGCGTCAGGATCACAACGGCTTCTCCCACCAGGATCCGGGTGTCACCTCCGTCCTGCTGAACAAGTGCTTCAACAACGATCACGTGATCGGCATCTACTTCCCGGTGGATTCCAACATGCTGCTCGCTGTGGCTGAGAAGTGCTACAAGTCCACCAACAAGATCAACGCCATCATCGCCGGCAAGCAGCCGGCCGCCACCTGGCTGACCCTGGACGAAGCTCGCGCCGAGCTCGAGAAGGGTGCTGCCGAGTGGAAGTGGGCTTCCAACGTGAAGTCCAACGATGAGGCTCAGATCGTGCTCGCCGCCACCGGTGATGTTCCGACTCAGGAAATCATGGCCGCTGCCGACAAGCTGGACGCCATGGGCATCAAGTTCAAGGTCGTCAACGTGGTTGACCTGGTCAAGCTGCAGTCCGCCAAGGAGAACAACGAGGCCCTCTCCGATGAGGAGTTCGCTGAGCTGTTCACCGAGGACAAGCCGGTCCTGTTCGCTTACCACTCCTATGCCCGCGACGTGCGTGGTCTGATCTACGATCGCCCGAACCACGACAACTTCAACGTTCACGGCTACGAGGAGCAGGGCTCCACCACCACCCCGTACGACATGGTTCGCGTGAACAACATCGATCGCTACGAGCTCCAGGCTGAAGCTCTGCGCATGATCGACGCTGACAAGTACGCCGACAAGATCAACGAGCTCGAGGCCTTCCGTCAGGAAGCCTTCCAGTTCGCTGTCGACAACGGCTACGATCACCCGGATTACACCGACTGGGTCTACTCCGGTGTCAACACCAACAAGCAGGGTGCTATCTCCGCTACCGCCGCAACCGCTGGCGATAACGAGTGA |
| **PK 5** | ATGAACGAGATCGCAAGAGCATCAAATCCTCTCACTGGTGAGGAACTGCGTAAGATCGATGCCTGGTGGCGCGCTGCGAACTATCTGAATGTCGGGCAGATCTATCTCTCGGCAAATCCGCTGTTGCGCGACAAGCTAACAGTCGAACACATCAAGCCACGTCTGCTCGGGCACTGGGGCACATCGCCCGGCCTGAACCTCATCTACGCTCACATGAACCGGCTGATCAGGAAATACGACCTCGATACGATCTATATGGCCGGTCCCGGCCATGGCGGCCCGGCCCTAGTAGCCAATGTCTACCTCGAAGGCAGTTATACCGAATTCTACCCCGAGGTGACACAGGACGAGGCCGGCCTTCTCAGGCTCTTCCGTCAATTCTCCACCCCCGGCGGCATTCCGAGCCATGTGAGCGTGCCAACACCCGGCTCGATCCATGAGGGTGGCGAACTTGGCTACGTGCTGGTGCACGCCTTCGGCGCCGTCATGGACAATCCCGACCTGATCGTCACGGCGGTTGTCGGCGACGGAGAAGCAGAGACAGGACCACTGGCTGGAAGCTGGAAAAGCATCGATTTCATCAATCCCGCCCGCGACGGGGCGGTGCTGCCGATCCTTCACCTCAACGGTTACAAGATTGCCGGTCCGACGGTGCTTGCGCGACATAGCGATGCGGATCTCGCGAAGTTTTTCGAAGGGCAAGGTTACGAACCTCACTTTGTCGAAGGCGATGTGCCCGAGATCGTTCATCAGCAATTCGCGGCCGTCCTGGAAAGCGCGATTATCAAGATCCGTGCTATTCAGTCGGATGCGCGCGAGAATGGCATGAGCAAGAGGCCGCTGTGGCCGATGATCGTGCTGCGCACGCCGAAAGGTTGGACCGGGCCAAAAATTGTCGATGGTTTGCCGGTCGAAGGGACATTCCGCGCGCACCAGGTGCCCGTGTCAGAAGTGCTGACCAGGCCCGGACATCTCCAGATCCTGGAAGACTGGATGCGCAGTTACAAGCCTGAAGAGCTGTTTGATGAGCACGGGGCCTTCCAGCCGGAATATGCCGCGCTGGCCCCGGCCGGCGACCGGCGTATGGGGTCGAACCCGCATGCCAACGGCGGCAAGCTGATCGTGCCGTTGAACCTGCCCGACTTCAATACATATGCCGTGACGCTTGAGCAGCGGGCGCGAGAGCGGATGGGCTCGACCGCTGTGCTCGGAGAATATCTCCGCGACATCTACACACGCAATCCCGACAACTTCCGGCTGTTCTGCCCCGACGAAACCAACTCCAACCGTCTCGGCGCGATTTTTGAAGTTTCCGATCGCTGCCTGGTCAGCCATATTCTGCCCGGTGACGACCATATCTCGCATGAAGGCCGGGTCATGGAGGTTCTGAGCGAGCATTGCTGCCACGGGTGGCTGGAAGGCTATACGCTGACCGGACGCCACGGTCTTTTTGCCACCTACGAGGCGTTCGCCATGATCGTCGATTCCATGTCGATGCAGCATGGCAAGTGGATGGAACATGCCAGACATGTTCCCTGGCGGGCCGATATTCCATCGCTCAACTATCTGCTCACGTCGACATGCTGGAGGAACGATCACAATGGTTTCAGTCACCAGGGGCCGGGCTTCATCGACACGATCATCCACCGCAAGCCGGCCGTCGCGCGTGTCTACCTTCCGCCGGACGCGAACTGCCTTCTCTCCGTTGCAGACCATTGCTTCCAAAGCCGGAATTACCTCAATCTGATTGTCATCGACAAACAGCCCCAGTTGCAGTGGCTGACGATGGAAGAGGCCAAAGCCCATTGCGCCAAGGGCGCCGGAATATGGGATATGTACAGCAACGAACCAGGTGAACCGGATGTCGTTCTGGCCTGCGCTGGCGACATACCAACGCAAGAGACCATTGCCGCAGCGTGGTTGCTGCGCCAGCATGCTCCCGATCTGAAAGTCCGCGTCGTGAACGTCGTCGACCTGATGCGGCTATGCCCGGCTGACCGCCATCCGCACGGCATGAGCGATACCGATTTCACCGCGATCTTTACCACGACCGCTCCGGTGATATTCACGTTTCACGGATATCCGGGCGTGATCCATGACCTGTTGCATGGGCGCGAAGCGCATGATCGTTTCCACGTCCGCGGCTATCTCGAGGAGGGCACGACGACGACGCCCTTCGACATGGTTGTGCTTAACAGGATCAGTCGTCTTCATCTGTGCCTGGATGTGCTGCGTTATGTTCCCGGCCTGCTGGTCAGCAATGCTGGCCTGGTTACCTTCTGCACGGACATGCTTTCGGAACATGAGCGTTATATCCGGGAGCATTTTGACGATCTGCCGGTGATCAAGGAATGGGTATGGTCGGAATAA |
| **PK 6** | ATGCCGCGGAAGCTCCAGAACCACCCGCAAGCGAAGGAATCCGTTTCTCCGAGCGGGGTGAGCGAGGAACTGCAGGATATCGCCCGCTATCGCCGCGCCGCCAACTATCTATCGGCCGCCCAGATCTACCTCAAGGATAATTGCCTGCTCGAACGGCCGCTCCTACCGGACGATATCAAAGACCGGCTACTCGGCCACTGGGGCACGGCGCCAGGCATCAACCTCGTCTACGCGCATCTCAACCGGCTAATCCGCGACCACCAGGCCAGTATTCTCCTGATTACCGGCCCGGGTCACGGGGCCGTGGCCAATCTCGCGAATCTCTACCTCGAAGGCACGCTGGCCGAATTCTATCCCGACCTGACCCTTGACCGTGCCGGACTGACGCGCTTCTGTCGCAGTTTCTCCTGGCCGGGGGGCTTCCCCAGCCATCTCAATCCGCAGATCCCCGGCACGATTCACGAGGGGGGAGAGCTTGGCTACGCGCTCTCGACCGCCTTTGGAGCCGCGCTGGATAATCCGGACCTTATCGTGGCCTGTATTGTCGGAGATGGCGAAGCCGAGACCGGACCGACCGCCACCGCCTGGCACAGCAATAAGTTCCTCGATCCGGCCACCTCGGGCGCCGTCCTGCCGATCCTCCACCTGAACGAGTACAAGATCTCGGGGCCGACCATCTTCGGCAGGATGTCGGACGGGGAACTCCTCTCCCTCTTCGAGGGCTATGGCTACGCCCCGCTGCTGGTCCAGGGTCCGGACCTCGACACAGCAACTTATCAGGCGATGGAGTGGGCCTACCAGCGCATCCGGGAGATCCAACGCCAGGCGAGAGCCGGCGTGCGCTTGGAACGCCCGGCGTGGCCGTTACTCATCCTGCGAAGCCCGAAGGGTTGGACCGGCATCAAGGAAATCGGGGGCAAGCCGATCGAAGGCTCGTTCCGCGCTCACCAGGTCCCGGCTGCCGATGCCAAGACGAATCCGGCAAGCCTGAAGCTCGTCGAAGCCTGGCTCCGGTCCTATCACCCGGAGGAACTCTTCGATAGCGAAGGCCACCCCGCGCCGGATATCCTCGCAACGTGCCCAACGGGCGATCGCCGGATGGGCATGAACCCCCACGCCTACGGCGGGGACATCCGCCGGAATCCGAGCCTACCGCCACACTGGGAGGATTACGGACTCGACGTGACGCATCGTGGCGCACCGATGGCGAGCAGCGTGGCCCAGTTCGGCAACTACTTGCGAGACACGATTGAACGGAGCAAGACGGCGTGGAACTTCCGCATCGCCTCGCCGGATGAGCTGACATCCAACCGCCTGGGCGCGGTACTTGAGGCGACCGACCGCGCCTTCGTCTGGCCGATTAAGCCGACGGACGACCACCTGGCTCCGGATGGACTGGTAATGGAGATCCTGAGCGAGCATACCTGCCAGGGATGGCTCCAGGGCTATCTCCTCACCGGCCGCCACGGGCTCTTCCCCTGCTATGAAGCCTTCATCATGATCGTCGACTCGATGCTCAACCAGTACGCCAAATTCATGAAAGTGGCGGCGGAGATTCCCTGGCGCAAGCCCATTTCTTCCTTGAACTACCTCCTCACCTCGCACAGCTGGCGGCAAGACCACAACGGCTACTCCCATCAGGGTCCAGGATTCATCGACCAGCTCCTCACCAAAAAGGCGTGGATGGTCCGGATCTACCTCCCGCCCGATGCCAACTGCCTGCTCCAGACGATGGATCACTGCTTCTGGAGCAAGAACTACATCAATCTGGTCATCGCCTCGAAGCAGCCGATGCCGCAATGGCTCACCAAGGAAGAAGCGCTGGAGCACTGCCGGATGGGCGCCTCGATCTGGCGCTGGGCCAGCACCGATGACGGCAAGAACCCCGACGTCGTGCTGGCGGCGGCTGGCGACATTTTGACTTTGGAGATGATGGCAGCGGTGAGGCTGTTGCAGCAAGACCTCCCGGATCTCCGCGTCAGGGTGGTGAATGTAACCGATCTCATGGTGTTGGGTCTCGATACCGAGCACCCGCACGGCCTGACACCCGAGGCATTCGACGAACTCTTCACGCCGGATCGGCTGGTCATCTTCAACTTCCACGGCTATCCGGGCGCGGTGAAACAACTCCTCTTCGGGAGACCGAGCCCCGGCCGGTTCCAGGTTAATGGCTACCAGGAAGAGGGGACCACCACGACACCGTTCGACATGCACGTCCGCAACGGCACGAGCCGGTATCACCTGGTGATGCAAGCCGCCCAAGCCTGCTCCGAGAAGCTCAGCCCGGGAACGATCTGCGACATCGAGGAACGGTACCGGCGGAAACTGGCTGAGCACCGGAAATACATAGAGGAGCACGGGATCGATCCGCCCGAGATCCGGGATTGGACATGGTAG |
| **PK 7** | ATGAACGAGATCGCAAGAGCATCAAATCCTCTCACTGGTGAGGAACTGCGTAAGATCGATGCCTGGTGGCGCGCTGCGAACTATCTGAATGTCGGGCAGATCTATCTCTCGGCAAATCCGCTGTTGCGCGACAAGCTAACAGTCGAACACATCAAGCCACGTCTGCTCGGGCACTGGGGCACATCGCCCGGCCTGAACCTCATCTACGCTCACATGAACCGGCTGATCAGGAAATACGACCTCGATACGATCTATATGGCCGGTCCCGGCCATGGCGGCCCGGCCCTAGTAGCCAATGTCTACCTCGAAGGCAGTTATACCGAATTCTACCCCGAGGTGACACAGGACGAGGCCGGCCTTCTCAGGCTCTTCCGTCAATTCTCCACCCCCGGCGGCATTCCGAGCCATGTGAGCGTGCCAACACCCGGCTCGATCCATGAGGGTGGCGAACTTGGCTACGTGCTGGTGCACGCCTTCGGCGCCGTCATGGACAATCCCGACCTGATCGTCACGGCGGTTGTCGGCGACGGAGAAGCAGAGACAGGACCACTGGCTGGAAGCTGGAAAAGCATCGATTTCATCAATCCCGCCCGCGACGGGGCGGTGCTGCCGATCCTTCACCTCAACGGTTACAAGATTGCCGGTCCGACGGTGCTTGCGCGACATAGCGATGCGGATCTCGCGAAGTTTTTCGAAGGGCAAGGTTACGAACCTCACTTTGTCGAAGGCGATGTGCCCGAGATCGTTCATCAGCAATTCGCGGCCGTCCTGGAAAGCGCGATTATCAAGATCCGTGCTATTCAGTCGGATGCGCGCGAGAATGGCATGAGCAAGAGGCCGCTGTGGCCGATGATCGTGCTGCGCACGCCGAAAGGTTGGACCGGGCCAAAAATTGTCGATGGTTTGCCGGTCGAAGGGACATTCCGCGCGCACCAGGTGCCCGTGTCAGAAGTGCTGACCAGGCCCGGACATCTCCAGATCCTGGAAGACTGGATGCGCAGTTACAAGCCTGAAGAGCTGTTTGATGAGCACGGGGCCTTCCAGCCGGAATATGCCGCGCTGGCCCCGGCCGGCGACCGGCGTATGGGGTCGAACCCGCATGCCAACGGCGGCAAGCTGATCGTGCCGTTGAACCTGCCCGACTTCAATACATATGCCGTGACGCTTGAGCAGCGGGCGCGAGAGCGGATGGGCTCGACCGCTGTGCTCGGAGAATATCTCCGCGACATCTACACACGCAATCCCGACAACTTCCGGCTGTTCTGCCCCGACGAAACCAACTCCAACCGTCTCGGCGCGATTTTTGAAGTTTCCGATCGCTGCCTGGTCAGCCATATTCTGCCCGGTGACGACCATATCTCGCATGAAGGCCGGGTCATGGAGGTTCTGAGCGAGCATTGCTGCCACGGGTGGCTGGAAGGCTATACGCTGACCGGACGCCACGGTCTTTTTGCCACCTACGAGGCGTTCGCCATGATCGTCGATTCCATGTCGATGCAGCATGGCAAGTGGATGGAACATGCCAGACATGTTCCCTGGCGGGCCGATATTCCATCGCTCAACTATCTGCTCACGTCGACATGCTGGAGGAACGATCACAATGGTTTCAGTCACCAGGGGCCGGGCTTCATCGACACGATCATCCACCGCAAGCCGGCCGTCGCGCGTGTCTACCTTCCGCCGGACGCGAACTGCCTTCTCTCCGTTGCAGACCATTGCTTCCAAAGCCGGAATTACCTCAATCTGATTGTCATCGACAAACAGCCCCAGTTGCAGTGGCTGACGATGGAAGAGGCCAAAGCCCATTGCGCCAAGGGCGCCGGAATATGGGATATGTACAGCAACGAACCAGGTGAACCGGATGTCGTTCTGGCCTGCGCTGGCGACATACCAACGCAAGAGACCATTGCCGCAGCGTGGTTGCTGCGCCAGCATGCTCCCGATCTGAAAGTCCGCGTCGTGAACGTCGTCGACCTGATGCGGCTATGCCCGGCTGACCGCCATCCGCACGGCATGAGCGATACCGATTTCACCGCGATCTTTACCACGACCGCTCCGGTGATATTCACGTTTCACGGATATCCGGGCGTGATCCATGACCTGTTGCATGGGCGCGAAGCGCATGATCGTTTCCACGTCCGCGGCTATCTCGAGGAGGGCACGACGACGACGCCCTTCGACATGGTTGTGCTTAACAGGATCAGTCGTCTTCATCTGTGCCTGGATGTGCTGCGTTATGTTCCCGGCCTGCTGGTCAGCAATGCTGGCCTGGTTACCTTCTGCACGGACATGCTTTCGGAACATGAGCGTTATATCCGGGAGCATTTTGACGATCTGCCGGTGATCAAGGAATGGGTATGGTCGGAATAA |
| **fls** | ATGGCGATGATAACTGGAGGGGAACTGGTGGTCCGGACCCTGATTAAAGCTGGCGTAGAACAACTGTTTGGCCTGCATGGCATTCATATTGACACCATTTTTCAGGCTTGCCTGGACCACGACGTCCCAATCATTGATACTCGCCACGAAGCGGCGGCAGGCCACGCTGCGGAAGGTTATGCCCGCGCGGGCGCTAAACTGGGTGTTGCCCTGGTGACCGCTGGCGGTGGCTTTACCAATGCCGTTACGCCGATCGCGAACGCTCGGACCGATCGCACTCCGGTTCTGTTCCTGACCGGTTCTGGTGCTCTTCGTGATGACGAAACCAACACCCTGCAGGCCGGTATTGATCAGGTGGCCATGGCGGCCCCGATCACGAAATGGGCTCATCGTGTTATGGCAACTGAACACATCCCGCGTCTGGTTATGCAGGCCATTCGTGCCGCTCTGAGCGCCCCACGTGGCCCGGTGCTGCTGGATCTGCCATGGGACATCCTGATGAACCAAATCGATGAAGATTCCGTTATCATCCCAGACCTGGTGCTGTCTGCTCACGGTGCCCATCCAGACCCGGCTGACCTGGACCAGGCTCTGGCACTGCTGCGTAAAGCCGAACGCCCAGTTATCGTACTGGGCTCCGAGGCGTCCCGCACCGCACGCAAGACCGCACTGAGCGCATTCGTAGCGGCGACCGGTGTACCGGTTTTCGCTGACTATGAAGGCCTGTCCATGCTGAGCGGCCTGCCGGACGCTATGCGTGGCGGCCTGGTGCAGAACCTGTACTCCTTTGCAAAAGCTGATGCAGCTCCGGACCTGGTACTGATGCTGGGTGCTCGTTTCGGTCTGAACACCGGTCATGGTTCCGGTCAACTGATCCCGCATTCTGCTCAGGTGATCCAGGTGGATCCAGACGCGTGTGAACTGGGTCGCCTGCAAGGCATCGCGCTGGGTATCGTGGCTGATGTAGGTGGCACCATTGAAGCGCTGGCTCAGGCGACCGCACAGGACGCCGCGTGGCCGGACCGCGGCGACTGGTGCGCCAAGGTAACTGACCTGGCCCAGGAGCGTTACGCTTCCATCGCGGCTAAATCCAGCTCTGAACATGCGCTGCACCCGTTCCACGCTTCTCAGGTTATCGCGAAACACGTGGACGCAGGCGTGACCGTCGTTGCGGATGGTGGCCTGACTTATCTGTGGCTGTCCGAAGTTATGTCTCGTGTCAAACCAGGCGGCTTCCTGTGCCACGGCTATCTGAACAGCATGGGTGTAGGCTTCGGTACTGCCCTGGGTGCGCAGGTTGCGGATCTGGAGGCAGGTCGTCGTACCATCCTGGTGACCGGCGACGGCTCTGTTGGTTATTCCATTGGCGAATTCGACACCCTGGTACGCAAACAGCTGCCGCTGATTGTAATTATCATGAACAACCAGTCTTGGGGCTGGACCCTGCACTTTCAGCAGCTGGCCGTTGGTCCTAACCGTGTCACCGGCACCCGCCTGGAAAATGGTTCCTATCACGGCGTTGCTGCGGCATTCGGTGCTGATGGTTACCACGTCGACTCTGTCGAGAGCTTCAGCGCCGCTCTGGCTCAGGCACTGGCACACAACCGCCCGGCATGCATCAACGTTGCTGTGGCCCTGGACCCGATCCCGCCGGAGGAACTGATCCTGATTGGCATGGACCCGTTTGCG |
| **gals** | ATGGCTTCTGTTCACGGTACCACCTACGAACTGCTGCGTCGTCAGGGTATCGACACCGTTTTCGGTAACCCGGGTTCTAACGAACTGCCGTTCCTGAAAGACTTCCCGGAAGACTTCCGTTACATCCTGGCTCTGCAGGAAGCTTGCGTTGTTGGTATCGCTGACGGTTACGCTCAGGCTTCTCGTAAACCGGCTTTCATCAACCTGCACTCTGCTGCTGGTACCGGTAACGCTATGGGTGCTCTGTCTAACGCTCGTACCTCTCACTCTCCGCTGATCGTTACCGCTGGTCAGCAGACCCGTGCTATGATCGGTGTTGAAGCTGGTGAAACCAACGTTGACGCTGCTAACCTGCCGCGTCCGCTGGTTAAATGGTCTTACGAACCGGCTTCTGCTGCTGAAGTTCCGCACGCTATGTCTCGTGCTATCCACATGGCTTCTATGGCTCCGCAGGGTCCGGTTTACCTGTCTGTTCCGTACGACGACTGGGACAAAGACGCTGACCCGCAGTCTCACCACCTGTTCGACCGTCACGTTTCTTCTTCTGTTCGTCTGAACGACCAGGACCTGGACATCCTGGTTAAAGCTCTGAACTCTGCTTCTAACCCGGCTATCGTTCTGGGTCCGGACGTTGACGCTGCTAACGCTAACGCTGACTGCGTTATGCTGGCTGAACGTCTGAAAGCTCCGGTTTGGGTTGCTCCGTCTGCTCCGCGTTGCCCGTTCCCGACCCGTCACCCGTGCTTCCGTGGTCTGATGCCGGCTGGTATCGCTGCTATCTCTCAGCTGCTGGAAGGTCACGACGTTGTTCTGGTTATCGGTGCTCCGGTTTTCCGTTACGTTTTTTACGACCCGGGTCAGTACCTGAAACCGGGTACCCGTCTGATCTCTGTTACCTGCGACCCGCTGGAAGCTGCTCGTGCTCCGATGGGTGACGCTATCGTTGCTGACATCGGTGCTATGGCTTCTGCTCTGGCTAACCTGGTTGAAGAATCTTCTCGTCAGCTGCCGACCGCTGCTCCGGAACCGGCTAAAGTTGACCAGGACGCTGGTCGTCTGCACCCGGAAACCGTTTTCGACACCCTGAACGACATGGCTCCGGAAAACGCTATCTACCTGAACGAATCTACCTCTACCACCGCTCAGATGTGGCAGCGTCTGAACATGCGTAACCCGGGTTCTTACTACTTCTGCGCTGCTGGTGGTCTGGGTTTCGCTCTGCCGGCTGCTATCGGTGTTCAGCTGGCTGAACCGGAACGTCAGGTTATCGCTGTTATCGGTGACGGTTCTGCTAACTACTCTATCTCTGCTCTGTGGACCGCTGCTCAGTACAACATCCCGACCATCTTCGTTATCATGAACAACGGTACCTACGGTATGCTGCGTTGGTTCGCTGGTGTTCTGGAAGCTGAAAACGTTCCGGGTCTGGACGTTCCGGGTATCGACTTCCGTGCTCTGGCTAAAGGTTACGGTGTTCAGGCTCTGAAAGCTGACAACCTGGAACAGCTGAAAGGTTCTCTGCAGGAAGCTCTGTCTGCTAAAGGTCCGGTTCTGATCGAAGTTTCTACCGTTTCTCCGGTTAAA |
| **PsLrhi** | ATGGCTGAATTCCGTATCGCTCAGGACGTTGTTGCTCGTGAAAACGACCGTCGTGCTTCTGCTCTGAAAGAAGACTACGAAGCTCTGGGTGCTAACCTGGCTCGTCGTGGTGTTGACATCGAAGCTGTTACCGCTAAAGTTGAAAAATTCTTCGTTGCTGTTCCGTCTTGGGGTGTTGGTACCGGTGGTACCCGTTTCGCTCGTTTCCCGGGTACCGGTGAACCGCGTGGTATCTTCGACAAACTGGACGACTGCGCTGTTATCCAGCAGCTGACCCGTGCTACCCCGAACGTTTCTCTGCACATCCCGTGGGACAAAGCTGACCCGAAAGAACTGAAAGCTCGTGGTGACGCTCTGGGTCTGGGTTTCGACGCTATGAACTCTAACACCTTCTCTGACGCTCCGGGTCAGGCTCACTCTTACAAATACGGTTCTCTGTCTCACACCGACGCTGCTACCCGTGCTCAGGCTGTTGAACACAACCTGGAATGCATCGAAATCGGTAAAGCTATCGGTTCTAAAGCTCTGACCGTTTGGATCGGTGACGGTTCTAACTTCCCGGGTCAGTCTAACTTCACCCGTGCTTTCGAACGTTACCTGTCTGCTATGGCTGAAATCTACAAAGGTCTGCCGGACGACTGGAAACTGTTCTCTGAACACAAAATGTACGAACCGGCTTTCTACTCTACCGTTGTTCAGGACTGGGGTACCAACTACCTGATCGCTCAGACCCTGGGTCCGAAAGCTCAGTGCCTGGTTGACCTGGGTCACCACGCTCCGAACACCAACATCGAAATGATCGTTGCTCGTCTGATCCAGTTCGGTAAACTGGGTGGTTTCCACTTCAACGACTCTAAATACGGTGACGACGACCTGGACGCTGGTGCTATCGAACCGTACCGTCTGTTCCTGGTTTTCAACGAACTGGTTGACGCTGAAGCTCGTGGTGTTAAAGGTTTCCACCCGGCTCACATGATCGACCAGTCTCACAACGTTACCGACCCGATCGAATCTCTGATCAACTCTGCTAACGAAATCCGTCGTGCTTACGCTCAGGCTCTGCTGGTTGACCGTGCTGCTCTGTCTGGTTACCAGGAAGACAACGACGCTCTGATGGCTACCGAAACCCTGAAACGTGCTTACCGTACCGACGTTGAACCGATCCTGGCTGAAGCTCGTCGTCGTACCGGTGGTGCTGTTGACCCGGTTGCTACCTACCGTGCTTCTGGTTACCGTGCTCGTGTTGCTGCTGAACGTCCGGCTTCTGTTGCTGGTGGTGGTGGTATCATCTAACTCGAG |
